## Supplementary Figures with caption for "A Boolean network model of hypoxia, mechanosensing and TGF-β signaling captures the role of phenotypic plasticity and mutations in tumor metastasis"

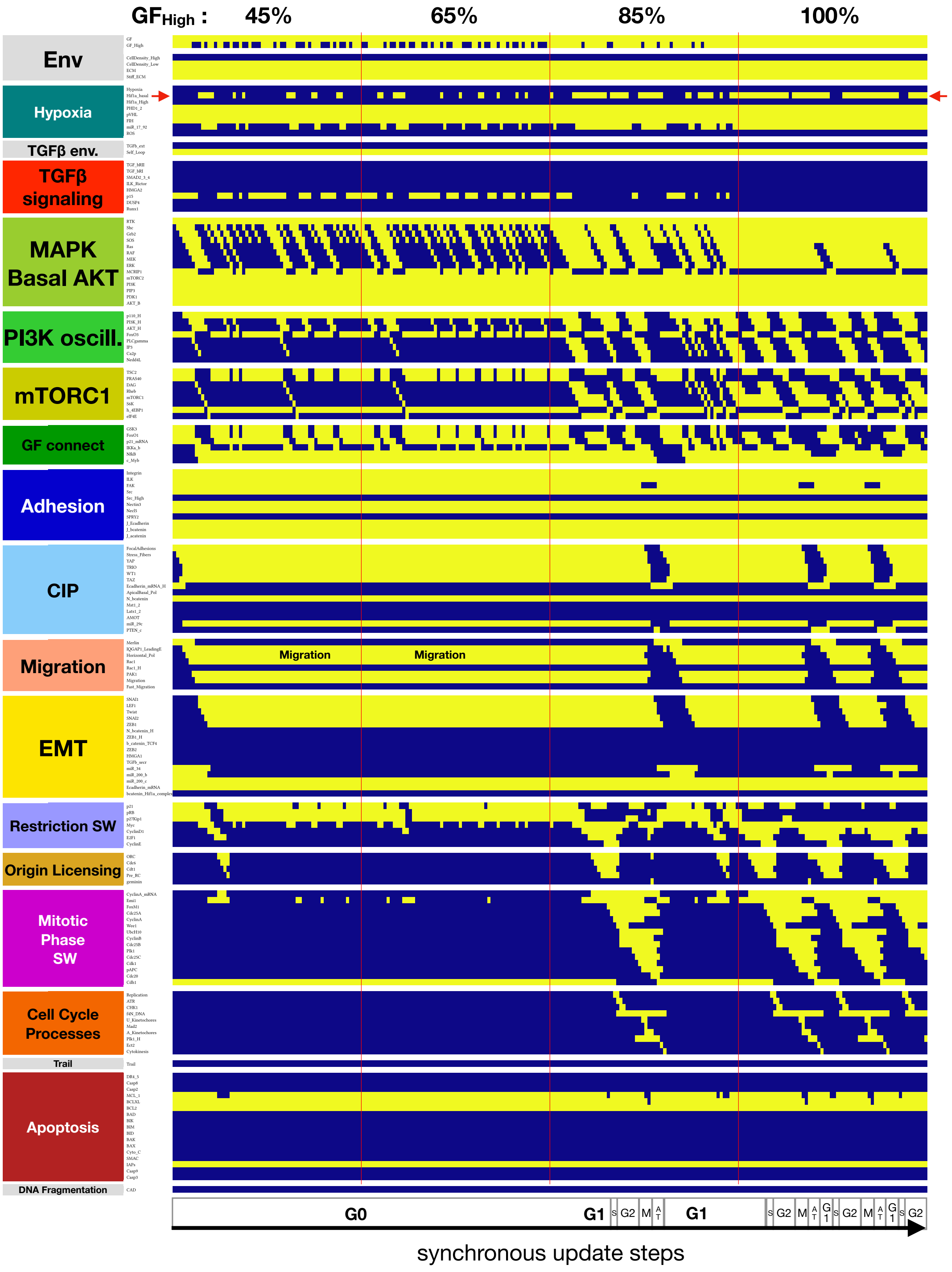


**SM Figure 1. Hif-1α is required for proliferation under normoxia (full version of Fig. 2A)**. A) Dynamics of relevant regulatory molecule expression/activity during exposure of a quiescent cell to increasing mitogen signaling (60 update-steps of 45%, 65%, 85% and 100% high GF; full network dynamics in SM Fig. 1A). *X-axis:* update steps annotated by cell cycle phase (G0: quiescence; G1: start of cell cycle entry; S: DNA synthesis; G2: growth phase 2; M: metaphase; A T: anaphase, telophase, cytokinesis); *y-axis*: nodes organized by regulatory module; *yellow/blue*: ON/OFF; *red arrow:* Hif1a_basal node.

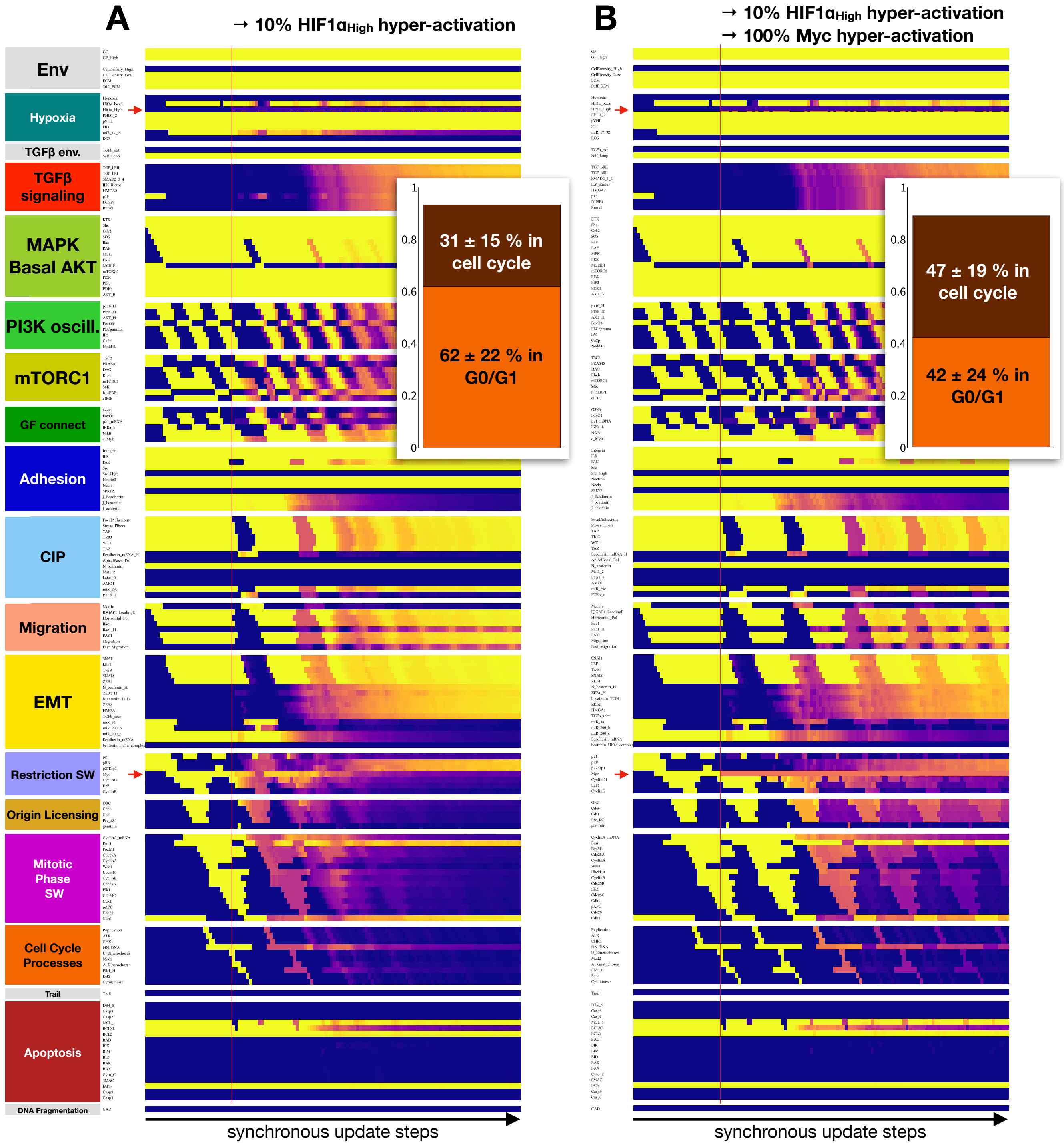


**SM Figure 2. Hif-1α hyper-activation blocks the cell cycle by repressing Myc. A)** Dynamics of average expression/activity of all regulatory molecules across 1000 cycling cells (*left side of vertical red line*) exposed to 10% Hif-1α hyper-activation (*right side of vertical red line*). *Inset:* fraction of time cells spend in quiescence (*orange*) vs. in cell cycle (*dark red*) following Hif-1α hyper-activation. **B)** Dynamics of average expression/activity of all regulatory molecules across 1000 cycling cells (*left side of vertical red line*) simultaneously exposed to 10% Hif-1α and 100% Myc hyper-activation (*right side of vertical red line*). *Inset:* fraction of time cells spend in quiescence (*orange*) vs. in cell cycle (*dark red*) following Hif-1α/Myc hyper-activation. *X-axis:* synchronous update steps; *y-axis*: nodes organized by regulatory module; *yellow/purple/blue color scale*: ON/50%/OFF; *red arrows:* Hif1a_High / Myc node. *Initial condition*: epithelial cells in GF:1, CellDensity_Low:1, GF_High:1, Stiff_ECM:1, Trail:0, Self_Loop:1, TGFb_ext:0, Hypoxia:0; *autocrine TGF-β*: 5% TGFb_secr knockdown.

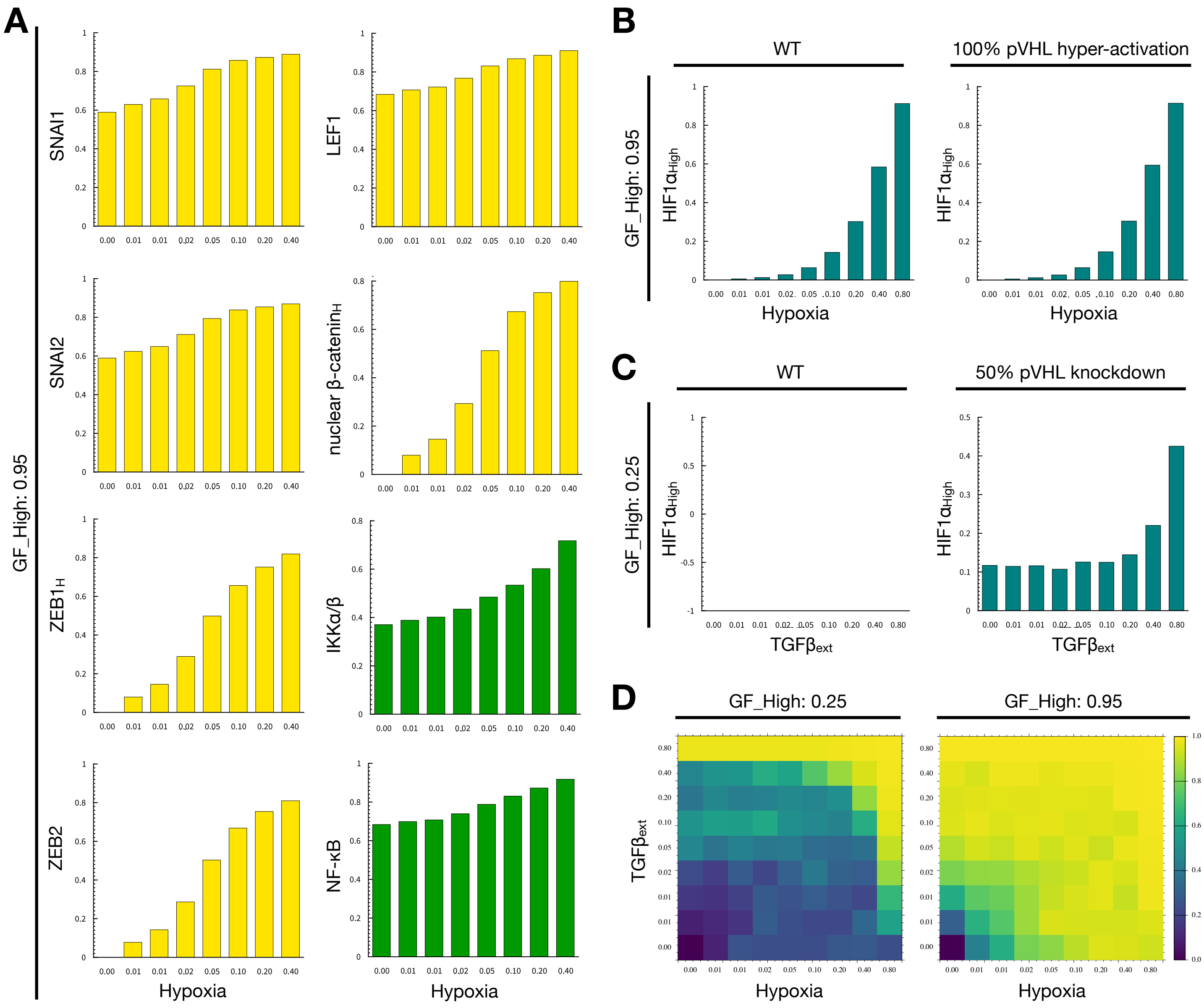


**SM Figure 3. Hypoxia induces EMT independently of TFG-β. A)** Average activation of mesenchymal factors SNAI1, SNAI2, ZEB1 (high expression node *ZEB1_H*), ZEB2, LEF1, nuclear nuclear β-catenin (high expression node *N_bcatenin_H*), IKKα/β and NF-κB in an ensemble of initially epithelial cells in response to increasing hypoxia exposure (log 2 hypoxia scale; *environment:* GF_High:0.95, TGFb_ext:0). **B)** Average Hif-1$\alpha_{high}$ activation in an ensemble of initially epithelial cells in response to increasing hypoxia exposure (log 2 hypoxia scale; *environment:* GF_High:0.95, TGFb_ext:0). *Left:* wild-type; *right:* 100% pVHL hyper-activation. **C)** Average Hif-1$\alpha_{high}$ activation in an ensemble of initially epithelial cells in response to increasing hypoxia exposure (log 2 hypoxia scale; *environment:* GF_High:0.25, Hypoxia:0). *Left:* wild-type; *right:* 50% pVHL knockdown. **D)** Fraction of time initially epithelial cells spend a mesenchymal state, as a function of hypoxia (*x axis*, log2 scale) and external TFG-β (*y axis*, log2 scale). *Left:* 25% GF_High; *right:* 95% GF_High. *Length of time-window for continuous runs:* 100 steps (~5 wild-type cell cycle lengths); *total sampled live cell time:* 100,000 steps; *update*: synchronous; *condition for all sampling runs*: CellDensity_Low:1, Stiff_ECM:1, Trail:0, Self_Loop:1; *autocrine TGF-β*: 5% TGFb_secr knockdown.

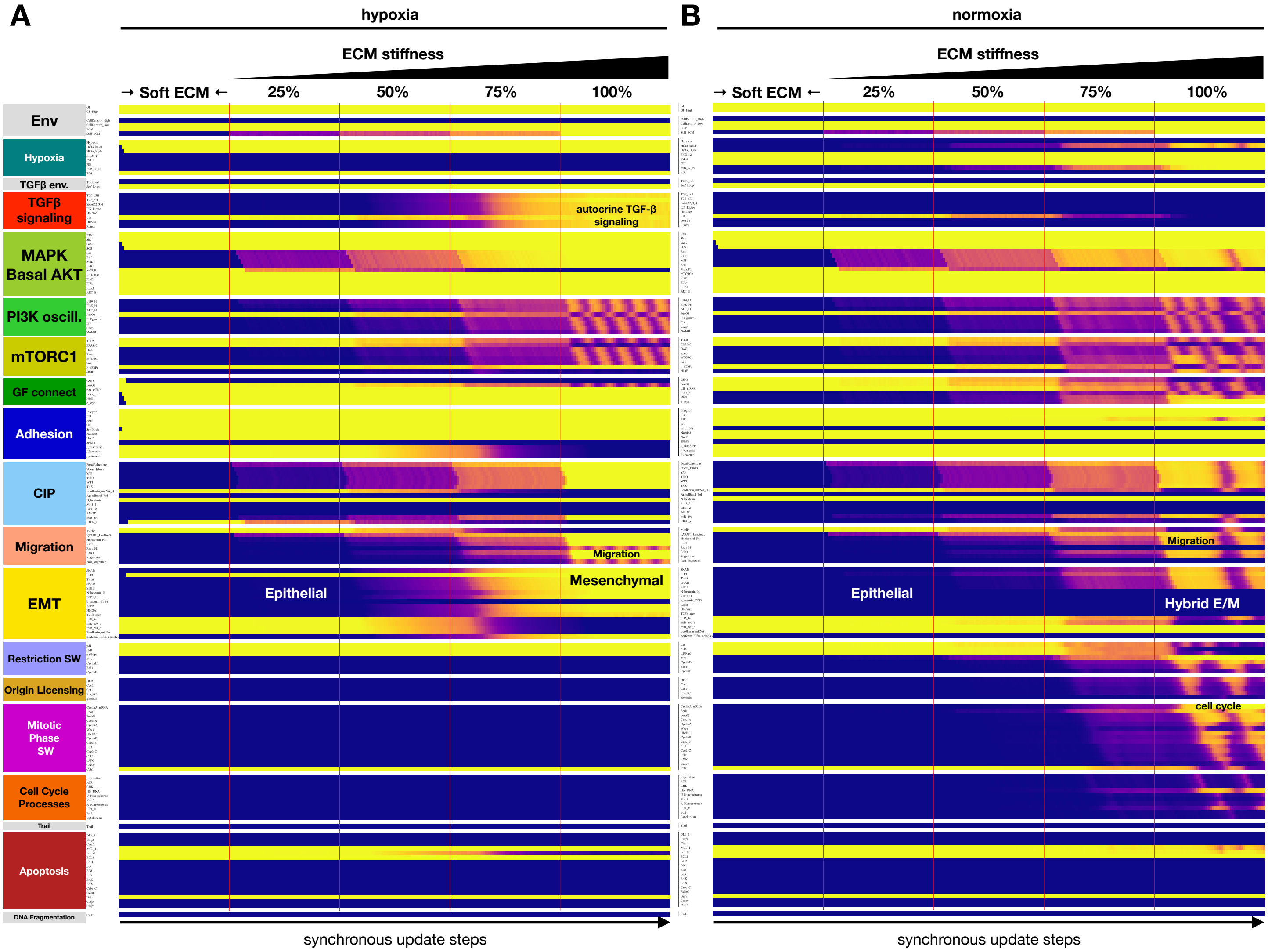


**SM Figure 4. A stiff ECM aids hypoxia-induced EMT, compared to hybrid E/M under normoxia. A)** Dynamics of regulatory molecule expression in a quiescent cell on a soft ECM under hypoxia, in response to increasing ECM stiffness (50 update-steps of 0,25,50,75 ans 100% Stiff_ECM = ON at saturating mitogen exposure and moderate cell density). Full time-course version of Fig. 4A. **B)** Dynamics of regulatory molecule expression in a quiescent cell on a soft ECM under normoxia, in response to increasing ECM stiffness (50 update-steps of 0,25,50,75 ans 100% Stiff_ECM = ON at saturating mitogen exposure and moderate cell density). *X-axis:* update steps; *y-axis*: nodes organized by regulatory module; *yellow/purple/ blue scale*: 100% ON/ 50% on / 100% OFF; *black/white labels:* relevant phenotypes; *update*: synchronous; *autocrine TGF-β*: 5% TGFb_secr knockdown.

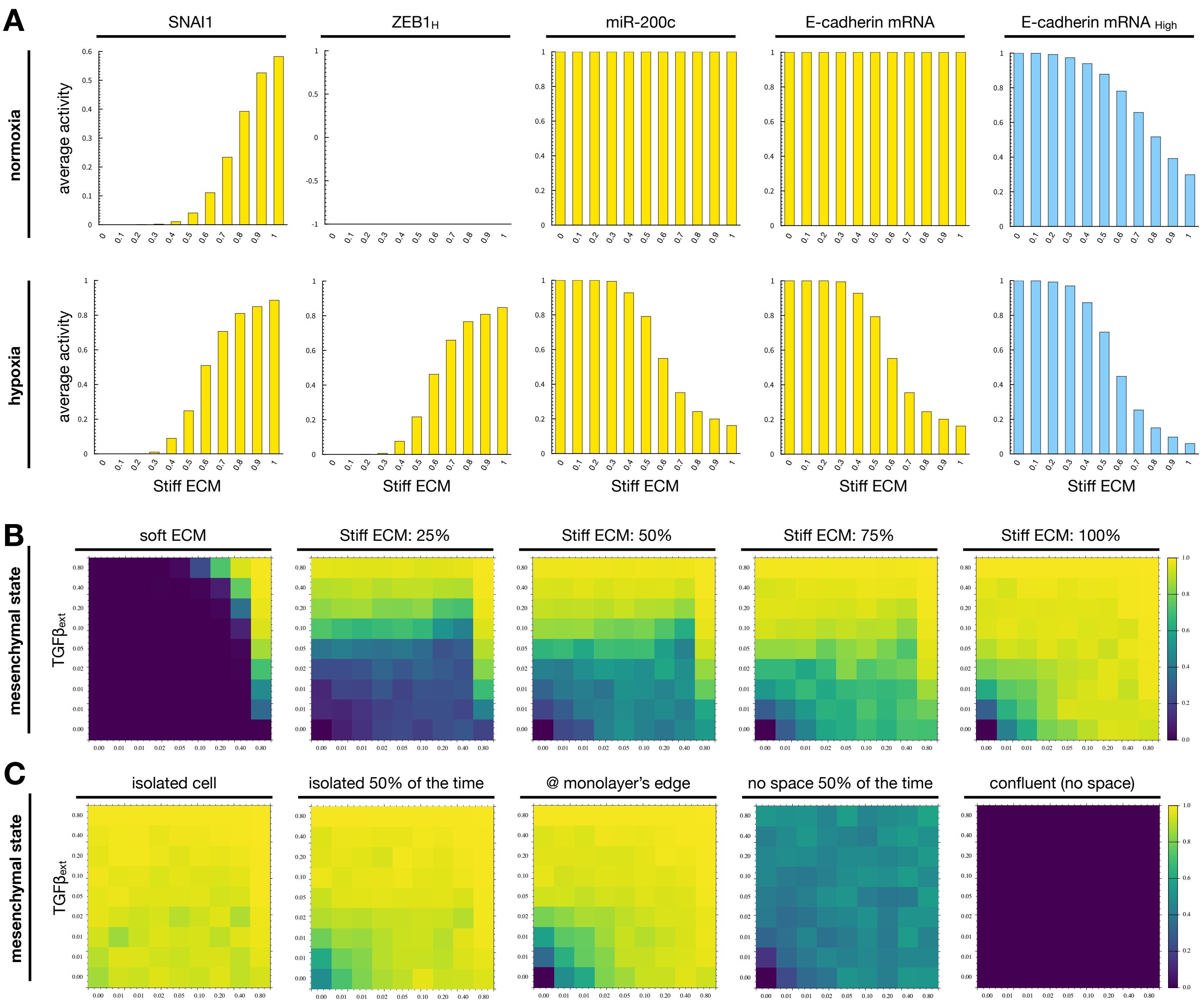


**SM Figure 5. ECM stiffness boosts both hypoxia- and TGF-β-induced EMT; high levels of both are required to drive EMT on very soft ECMs. A)** Average expression of SNAI1, high ZEB1, miR-200c and E-cadherin mRNA (*yellow*: baseline expression; *light blue:* high expression) in normoxic (*top*) vs. hypoxic (*bottom*) cells at moderate density and saturating mitogen exposure, plated on ECM of increasing stiffness (CellDensity_Low:1). **B)** Fraction of time initially epithelial cells spend a mesenchymal state, as a function of hypoxia (*x axis*) and external TFG-β (*y axis*) plated on ECMs of increasing stiffness (soft ECM, 25%, 50%, 75% and 100% Stiff_ECM; CellDensity_Low:1). **C)** Fraction of time initially epithelial cells spend a mesenchymal state, as a function of hypoxia (*x axis*) and external TFG-β (*y axis*) plated on ECMs of increasing cell density (isolated, 50% CellDensity_Low, 100% CellDensity_Low, 50% CellDensity_High, 100% CellDensity_High; Stiff_ECM:1). *Length of time-window for continuous runs:* 100 steps (~5 wild-type cell cycle lengths); *total sampled live cell time:* 100,000 steps; *update*: synchronous; *initial condition for all sampling runs*: epithelial cells in GF:1, CellDensity_Low:1, Stiff_ECM:1, Trail:0, Self_Loop:1, TGFb_ext:0, Hypoxia:0; *environment of sampling runs:* GF_High:0.95, *autocrine TGF-β loop:* 95% (5% TGFb_secr knockdown).

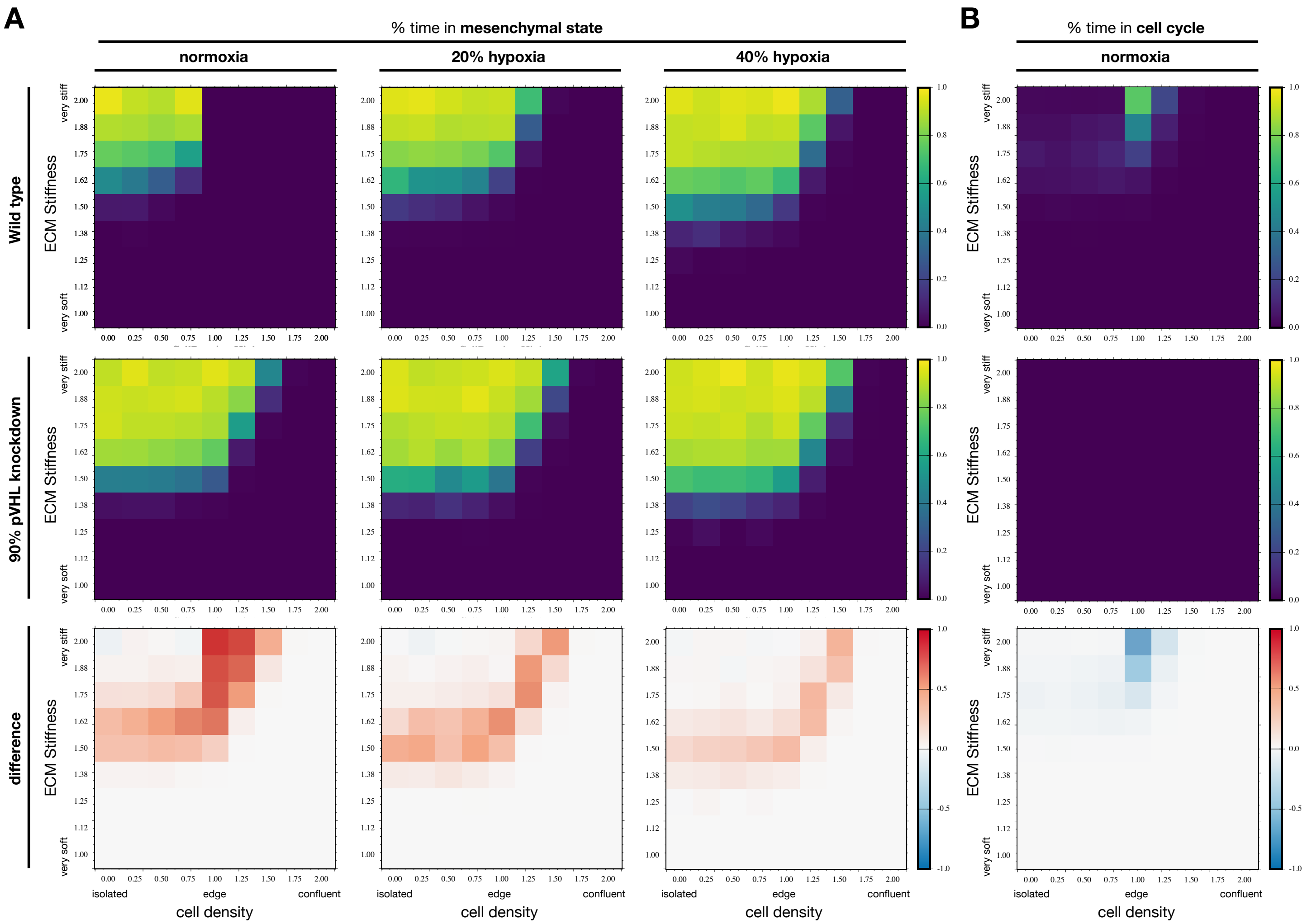


**SM Figure 6. VHL deficiency boosts bio mechanically induced EMT in normoxia as well as hypoxia at moderate ECM stiffness and medium-high density, but abolishes cell cycle entry. A)** Fraction of time initially epithelial cells spend in a mesenchymal state as a function of cell density (*x axis*) and ECM stiffness (*y axis*) in normoxia (*left column*), 20% hypoxia (*middle column*) and 40% hypoxia (*right column*). **B)** Fraction of time cells spend in the cell cycle under normoxic conditions, as a function of cell density (*x axis*) and ECM stiffness (*y axis*). (A-B) *Top row*: wild-type cells; *middle row*: VHL deficient cells (90% pVHL knockdown); *bottom row:* difference (pVHL - wild-type). *Length of time-window for continuous runs:* 100 steps (~5 wild-type cell cycle lengths); *total sampled live cell time:* 100,000 steps; *update*: synchronous; *condition for all sampling runs*: GF_High:0.95, TGF_ext:0, Trail:0, Self_Loop:1; *autocrine TGF-$\beta$*: 5% TGFb_secr knockdown.

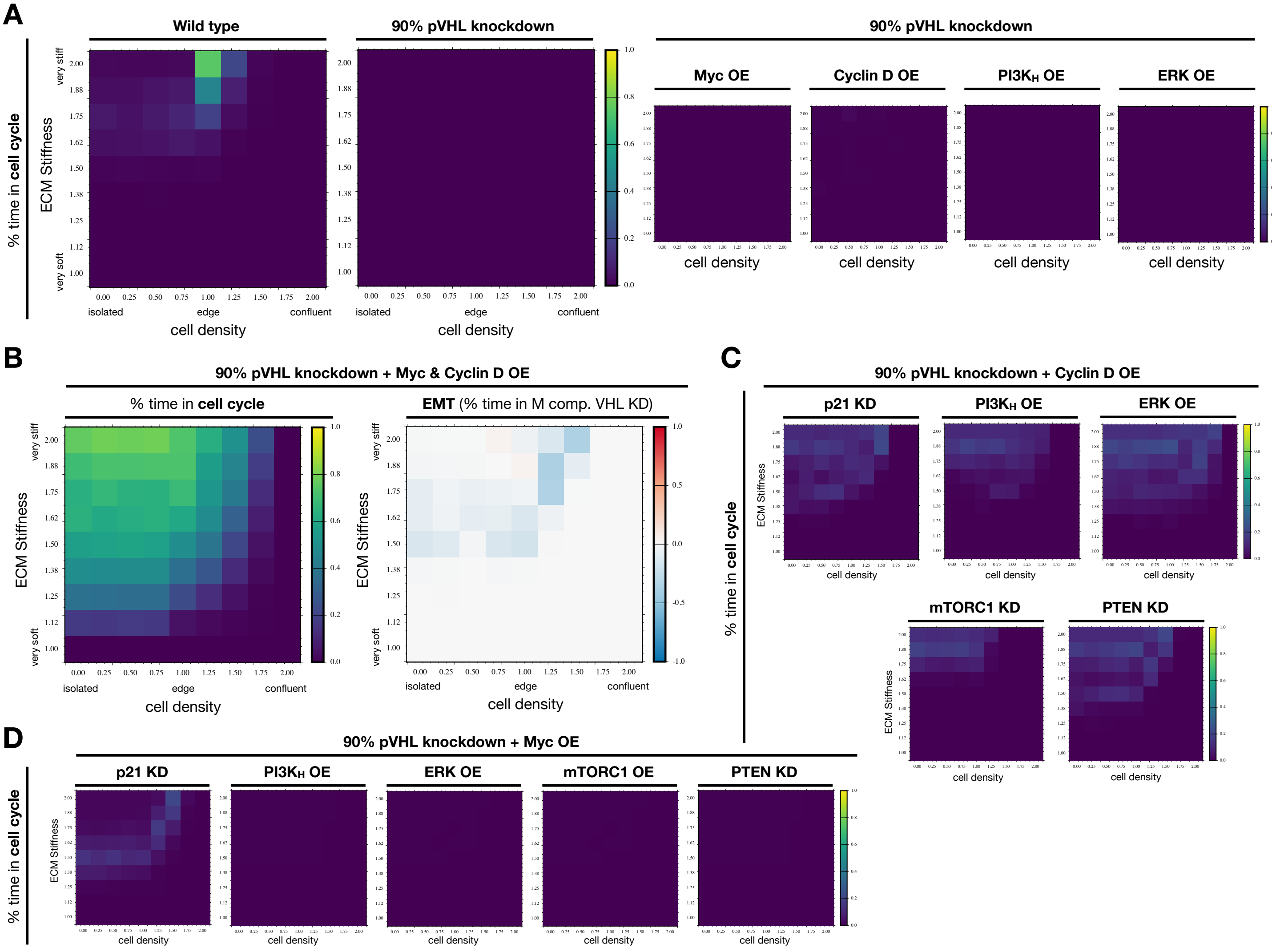


**SM Figure 7. VHL deficiency-induced cell cycle arrest is broken by cooperative hyper-activation of Myc and Cyclin D. A-B)** Fraction of time normoxic cells spend in the cell cycle as a function of cell density (*x axis*), ECM stiffness (*y axis*), and genetic background (wild-type, 90% pVHL knockdown, and 90% pVHL knockdown paired with a 100% ON lock of Myc, CyclinD1, PI3K_H or ERK). **B)** *Left:* Fraction of time normoxic cells with a 90% VHL deficiency and dual Myc/CyclinD1 hyper-activation (100% ON) spend in the cell cycle as a function of cell density (*x axis*), ECM stiffness (*y axis*). *Right:* Difference in the fraction of time normoxic cells with a 90% VHL deficiency and dual Myc/CyclinD1 hyper-activation (100% ON) spend in a mesenchymal state, compared to cells with VHL deficiency deficiency only, as a function of cell density (*x axis*), ECM stiffness (*y axis*). *Red/blue:* more/less time in M state in triple-mutant cells. **C-D)** Fraction of time normoxic cells with a 90% VHL deficiency and 100% CyclinD1 hyper-activation (*C*) or 100% Myc hyper-activation (*D*) spend in the cell cycle as a function of cell density (*x axis*), ECM stiffness (*y axis*), and additional genetic alterations (100% p21_mRNA:0, PI3K_H:1, ERK:1, mTORC1:1 or PTEN_c:0). (A-D) *Length of time-window for continuous runs:* 100 steps (~5 wild-type cell cycle lengths); *total sampled live cell time:* 100,000 steps; *update*: synchronous; *condition for all sampling runs*: GF_High:0.95, TGF_ext:0, Hypoxia:0, Trail:0, Self_Loop:1; *autocrine TGF-β*: 5% TGFb_secr knockdown.

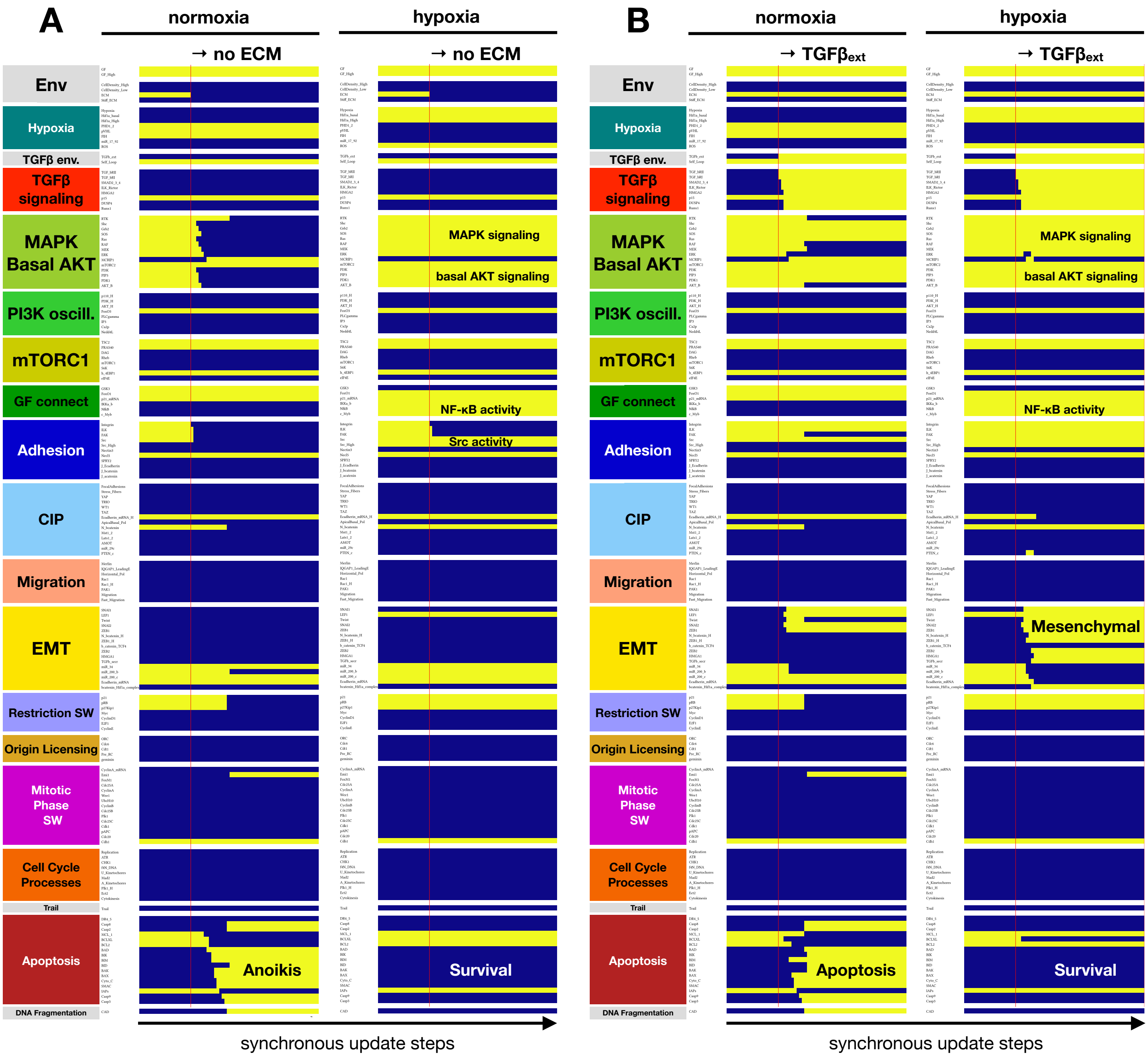


**SM Figure 8. Hypoxia confers anoikis resistance and protection from TGF-β induced epithelial cell apoptosis on a soft ECM (full version of Fig. 5). A)** Dynamics of regulatory molecule expression in a quiescent cell detached from a soft ECM in normoxia (left) vs. hypoxia (right) (20 update-steps on soft ECM, 50 update-steps detached, 100% high GF). **B)** Dynamics of regulatory molecule expression in a quiescent cell on a soft ECM, exposed to exogenous TGF-β in normoxia (left) vs. hypoxia (right) (20 update-steps no TGF-β, 50 update-steps 100% TGF-β). X-axis: update steps; y-axis: nodes organized by regulatory module; yellow/blue: ON/OFF; black/white labels: relevant phenotypes; update: synchronous.

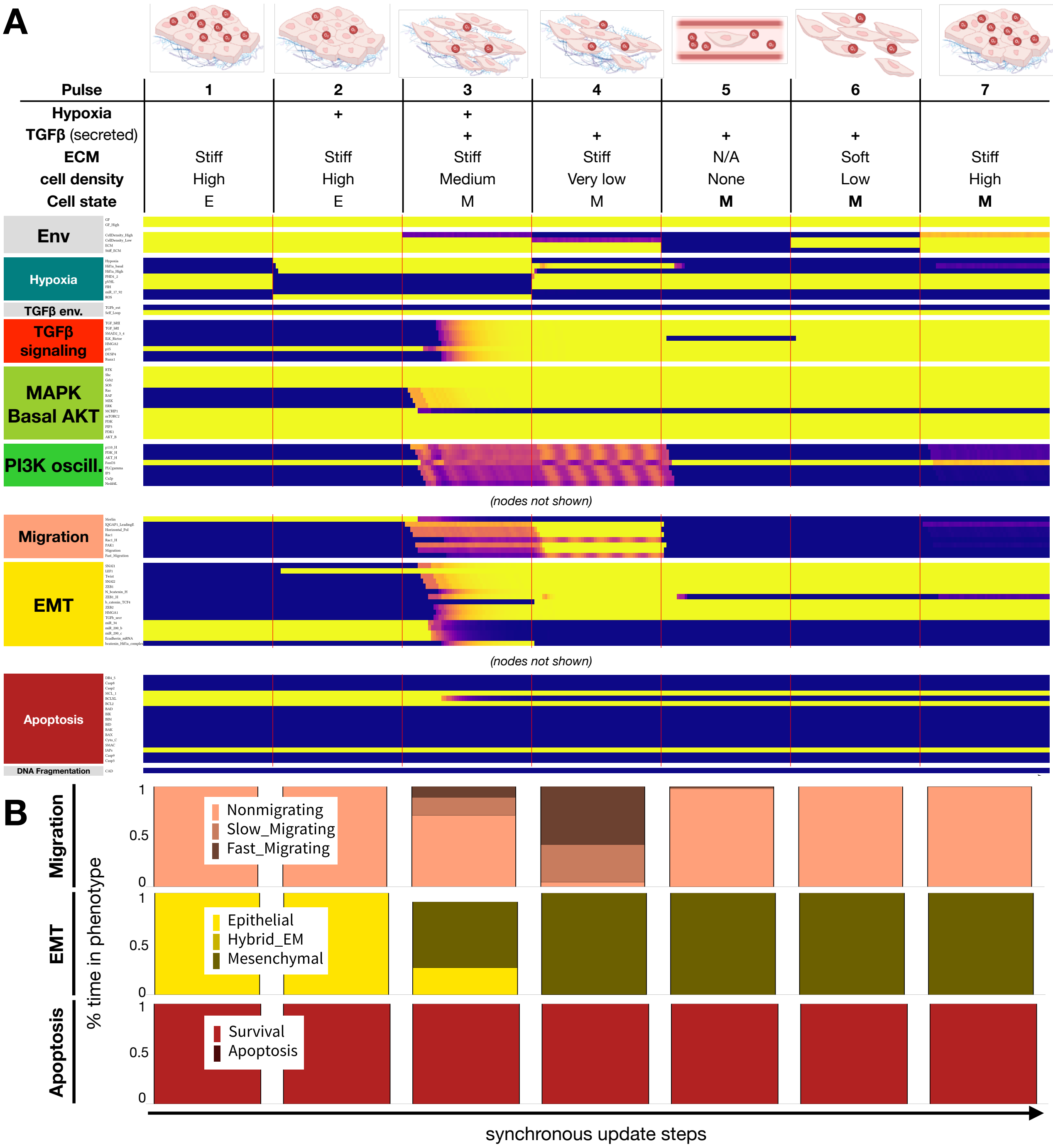


**SM Figure 9. Saturating autocrine TGF-β signaling protects all mesenchymal cells from anoikis, but also blocks MET on a stiff ECM. A)** Dynamics of relevant regulatory molecule expression in an ensemble of 1000 quiescent epithelial cell exposed to a sequence of microenvironments along the metastatic cascade (100 update-steps/pulse). *Pulse 1:* normoxia, stiff ECM, high density; *Pulse 2:* hypoxia, stiff ECM, high density; *Pulse 3:* hypoxia, stiff ECM, medium density; *Pulse 4* (invasion): normoxia, stiff ECM, very low density; *Pulse 5* (intravasation): normoxia, no ECM, no neighbors; *Pulse 6* (extravasation): normoxia, soft ECM, low density; *Pulse 7* (MET): normoxia, stiff ECM, high density. *X-axis:* update steps; *y-axis:* nodes organized by regulatory module; *yellow/blue:* ON/OFF; *black/white labels:* relevant phenotypes; *update:* synchronous; *autocrine TGF-β:* saturating. **B)** Average fraction of time an ensemble of 1000 cells spend in *top*: non-migrating (*light pink*), slow-migrating (*dark pink*) and fast-migrating (*brow-pink*) states; *middle*: epithelial (*yellow*), hybrid E/M (*dark yellow*) and mesenchymal (*mustard*) states; *bottom*: survival (*red*) vs. in apoptotic (*dark red*) states for each pulse along the metastatic cascade in panel A.

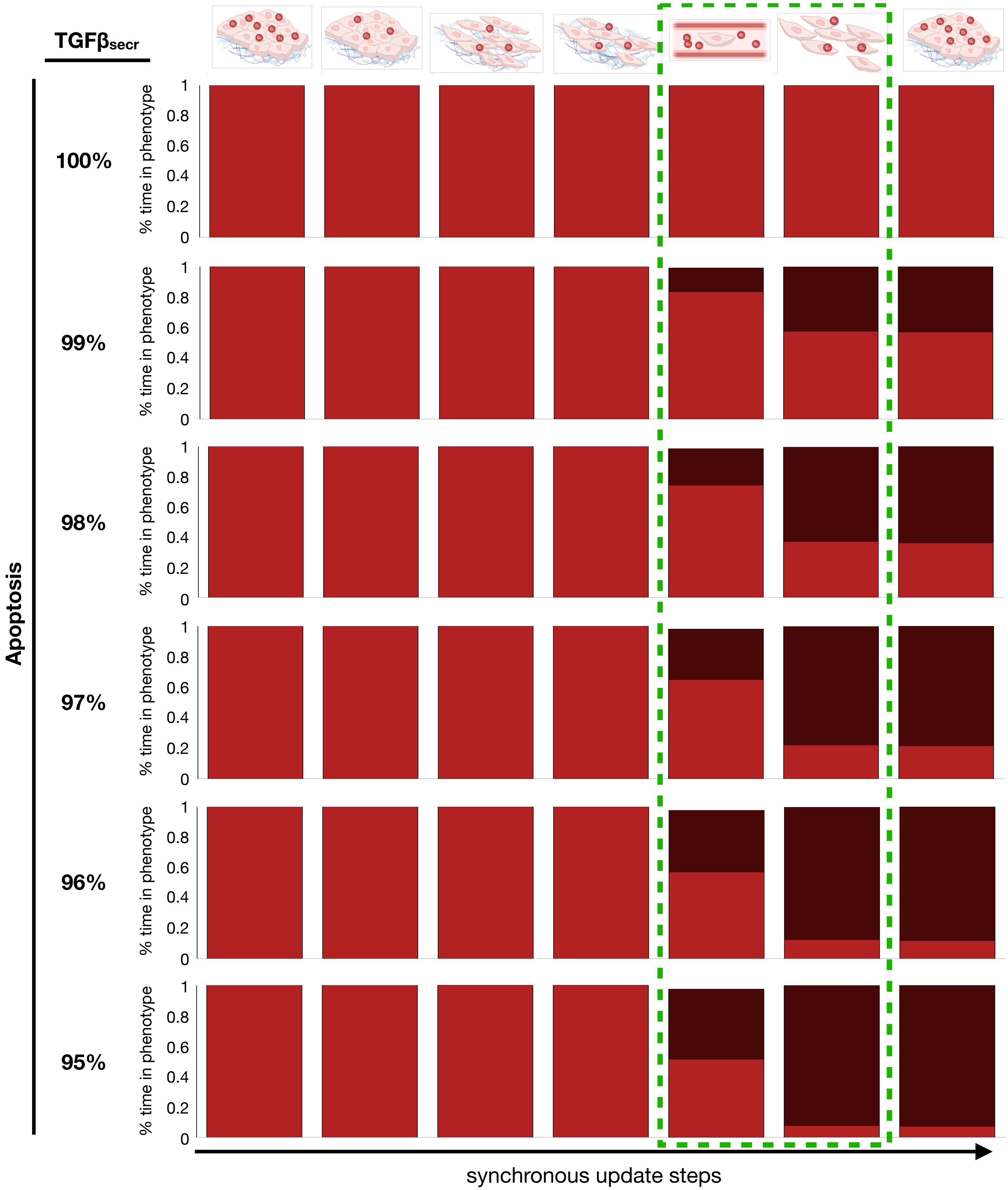


**SM Figure 10. Slight breaks in the autocrine TGF-β signaling loop are potent inducers of anoikis and apoptosis on soft ECM.** Average fraction of time an ensemble of 1000 cells spend in survival (*red*) vs. in apoptotic (*dark red*) states for each pulse along the metastatic cascade in Fig. 6 / SM Fig. 9, as a function of a weakening autocrine loop (1 to 5% TGFb_secr knockdown).

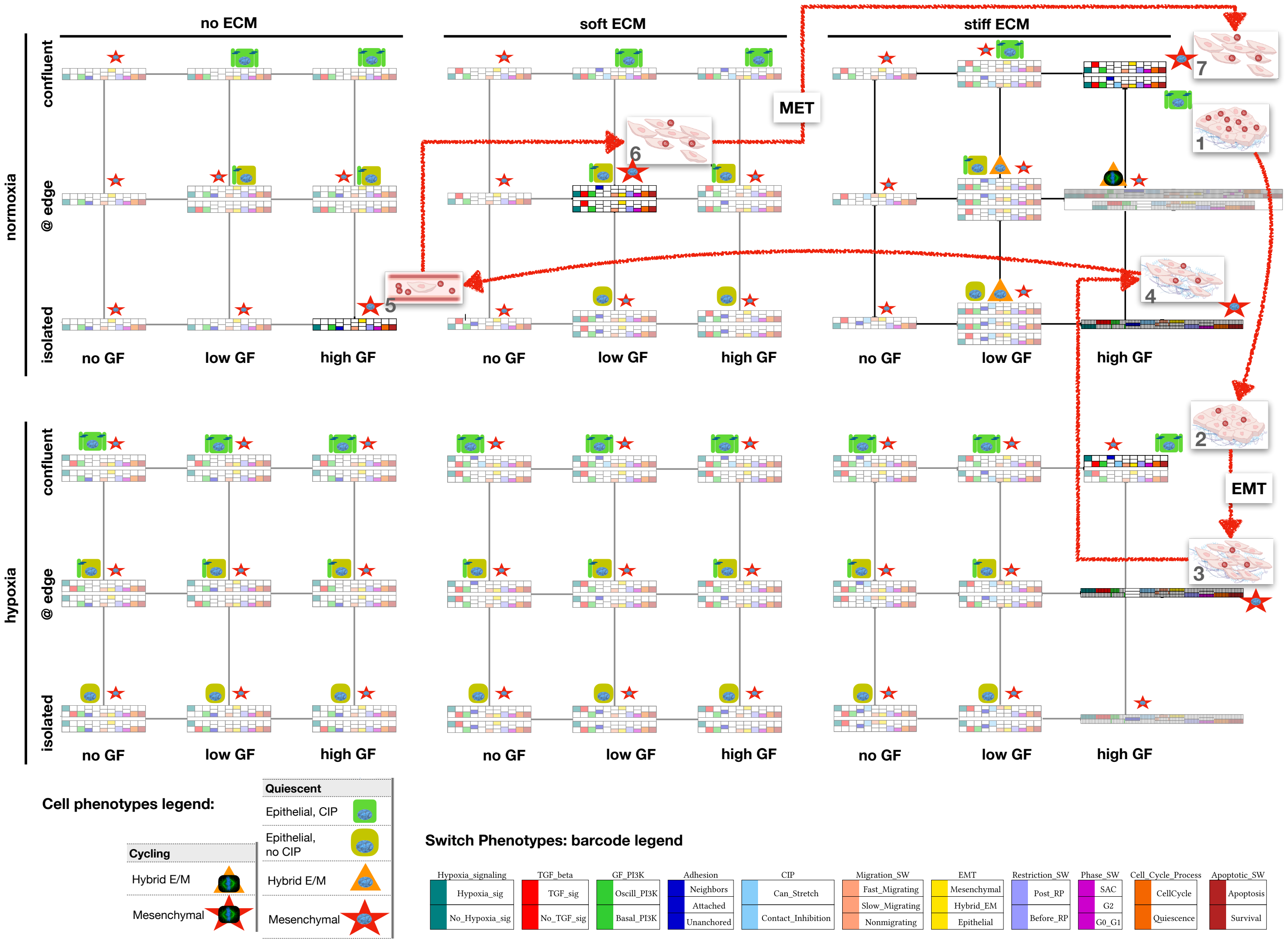


**SM Figure 11. A changing mix of model cell phenotypes as a function of mitogen, cell density, ECM stiffness and hypoxia allows us to track TME-driven EMT and MET during the metastatic cascade.** Summary of model cell states detected in every combination of low/ high growth-factor (*x axis*), isolated cells / moderate density / high cell density (*y axis*), no ECM / very soft ECM / very stiff ECM (*left to right*) in normoxia vs. hypoxia (*top vs. bottom*). Apoptotic and tetraploid quiescent cell states were omitted for clarity. Visual summaries of epithelial (*green/mustard sybmols*) / hybrid EM (*orange triangle*) / mesenchymal (*red star*) cell phenotypes (see cell phenotype legend, *bottom left*). Barcodes provide detailed phenotypic descriptions for each cell state (attractor); derived by comparing the expression of nodes in relevant modules to predetermined molecular signatures (e.g., apoptosis vs. survival), encoded in the *dmms* model file (barcode legend, *bottom right*). Oscillatory phenotypes have expanded barcodes that mark the transitions their regulatory switches undergo during the cycle (*high GF, stiff ECM*). *State transition arrows:* environment and cell phenotype (attractor) changes along the steps of the metastatic cascade shown in **Fig. 6**. *Saturated barcodes:* cell states along the cascade; *transparent barcodes:* all other diploid live cell states.
