## Supplementary File 2 for "A Boolean network model of hypoxia, mechanosensing and TGF-β signaling captures the role of phenotypic plasticity and mutations in tumor metastasis": Hypoxia Model Validation Script.py

```
import csv
import pandas as pd
import matplotlib.pyplot as plt
from mpl_toolkits import mplot3d
import seaborn as sns
from statannotations.Annotator import Annotator
from matplotlib.backends.backend_pdf import PdfPages
from mpl_toolkits.axes_grid1 import make_axes_locatable
import textwrap

p_ch_TH = 0.05  
# validation threshold: difference / average >= p_ch_TH

paper_list_firstAuthor = ["Wels", "Tang", "Gregory", "Casas", "Koshiji M", "Green SL", "Schwab LP", "Cummins EP", "Mallikarjuna P", "Lv Y", "Schokrpur S", "Whelan KA", "Zhang L"]

model_folder = ""
validation_csv = "ST2 - Hypoxia model validation list.csv"


with open(validation_csv, newline='\n') as csvfile:
	ST3 = csv.DictReader(csvfile, delimiter=',')
	csv_cols = []
	Stat_tests_output = []
	Stat_tests_output.append(['Environment','Env. change', 'perturbed node', 'perturbation type', 'bkgr (perturb or env)', 'bkgr type', 'downstream node in model','downstream phenotype in model','expected change direction','direction match','p-value'])

	BAD_tests_output = []
	BAD_tests_output.append(['Environment','Env. change', 'perturbed node', 'perturbation type', 'bkgr (perturb or env)', 'bkgr type', 'downstream node in model','downstream phenotype in model','expected change direction','direction match','p-value'])
	
	line_OK = 0
	some_in_line_OK = 0
	tests_OK = 0
	N_lines = 0
	N_tests =0
	
	fail_fig_IDs = []
	fig_ID = 0
	for exp_ID, col in enumerate(ST3):
		paper = col['paper']
		if(paper.split(',')[0] in paper_list_firstAuthor):
			change_dir = col['change direction']
			#print(pert_type)
			exp_name = "EXP "
			control_name = "CT "
	
			#perturbed node in model	
			node_st = col['perturbed node in model'].replace(" ", "")
			nodes_perturbed = [str(n) for n in node_st.split(',')]
			pert_type_str = col['perturbation type'].replace(" ", "")
			if (len(nodes_perturbed) > 0 and nodes_perturbed[0] != ''):
				pert_type = [int(n) for n in pert_type_str.split(',')]
				if (len(nodes_perturbed) == len(pert_type)):
					for i in range(0, len(nodes_perturbed)):
						if(pert_type[i] == 0):
							exp_name = exp_name + nodes_perturbed[i] + "_KD "
						else: exp_name = exp_name + nodes_perturbed[i] + "_OE "
				else: print("Line: " + node_st + "\n\t number of nodes perturbed does not match length of \'perturbation type\' list!")
			
			BKGR_str = col['bkgr (perturb or env: same in ct. vs. exp of comparison)'].replace(" ", "")
			BKGR_perturbed = [str(n) for n in BKGR_str.split(',')]
			BKGR_type_str = col['BKGR type'].replace(" ", "")
			if (len(BKGR_perturbed) > 0 and BKGR_perturbed[0] != ''):
				BKGR_type = [int(n) for n in BKGR_type_str.split(',')]
				if (len(BKGR_perturbed) == len(BKGR_type)):
					for i in range(0, len(BKGR_perturbed)):
						if(BKGR_type[i] == 0):
							exp_name = exp_name + BKGR_perturbed[i] + "_low "
							control_name = control_name + BKGR_perturbed[i] + "_low "
						else: 
							exp_name = exp_name + BKGR_perturbed[i] + "_high "
							control_name = control_name + BKGR_perturbed[i] + "_high "
				else: print("Line: " + BKGR_str +"\n\t number of background nodes  does not match length of \'perturbation type\' list!")
			
			
			#env change	 
			env_st = col['env name'].replace(" ", "")
			envs_perturbed = [str(n) for n in env_st.split(',')]
			env_type_str = col['Env change'].replace(" ", "")
			#print(envs_perturbed)
			if (len(envs_perturbed) > 0 and envs_perturbed[0] != ''):
				env_type = [int(n) for n in env_type_str.split(',')]
				#print(env_type)
				if(len(env_type) != 2):
					print("Env change should be 2 values\n" + env_st)
				else: 
					for i in range(0, len(envs_perturbed)):
						control_name = control_name + "__" + envs_perturbed[i] + ":" + str(env_type[0])
						exp_name = exp_name +  "__" + envs_perturbed[i] + ":" + str(env_type[1])
			
			
			
	
			#print("Control name: " + control_name)
			#print("Experiment name: " + exp_name)
		
			
			output_nodes_str = col['downstream node in model'].replace(" ", "")
			if(len(output_nodes_str)>0):
				output_nodes = [s for s in output_nodes_str.split(',')]
			else:
				output_nodes = []
			#print("output nodes string:" + output_nodes_str)
			#print(output_nodes)
						
			output_phen_str = col['downstream phenotype in model'].replace(" ", "")
			if(len(output_phen_str)>0):
				output_phen = [s for s in output_phen_str.split(',')]
			else:
				output_phen = []
			#print("output phenotypes string:" + output_phen_str)
			#print(output_phen)
				
			expected = col['change direction']
			ctr = col['In silico control name']
			exp = col['In silico experiment name']
			times_str = col['Time windows to compare'].replace(" ", "")
			times = [int(n) for n in times_str.split(',')]
			#print(times)
			attr_code = col['Attractor_code']
			csv_cols.append(col)
				
			if(output_nodes):
				N_lines = N_lines + 1
				#print(output_nodes)
				#print("N_lines = " + str(N_lines))
				fig, axes = plt.subplots(1, 1, figsize=(2.5*len(output_nodes)+len(exp_name)*0.2, 5), sharex=False, sharey=False)		
				df_nodes_list = []
				ctr_node_file = model_folder + "_EXP/General_Time_Series/"+ ctr + "/NodeBCh/" + attr_code + "_" + ctr + ".csv"
				#print(ctr_node_file)
				ok=0
				with open(ctr_node_file, newline='\n') as ct_file:
					CT = csv.reader(ct_file, delimiter=',', quotechar='"')
					#CT_cols = []
					for i in CT:
						#print(i[0])
						#print("ok = " + str(ok))
						if(ok==1 and (str(i[0]) in output_nodes)):
							#print("adding line:" + i[0])
							#CT_cols.append(i[0:10])
							for e in i[1:]:
								df_nodes_list.append([control_name,i[0],float(e)])
								#print("\t\tadding:" + i[0] + e)
						window=i[0][:-1]
						windows=window.split()
						#print(len(windows), windows)
						if(len(windows)>1):
							if(int(windows[1]) == times[0]-1): 
								ok=1
							else: ok=0
				#print(CT_cols)
				exp_node_file = model_folder + "_EXP/General_Time_Series/"+ exp + "/NodeBCh/" + attr_code + "_" + exp + ".csv"
				#print(ctr_node_file)
				ok=0
				with open(exp_node_file, newline='\n') as ct_file:
					EXP = csv.reader(ct_file, delimiter=',', quotechar='"')
					#EXP_cols = []
					for i in EXP:
						#print(i[0])
						#print(output_nodes)
						#print("ok = " + str(ok))
						if(ok==1 and (str(i[0]) in output_nodes)):
							#EXP_cols.append(i[0:10])
							#print("adding line:" + i[0])
							for e in i[1:]:
								df_nodes_list.append([exp_name,i[0],float(e)])
						window=i[0][:-1]
						windows=window.split()
						#print(len(windows), windows)
						if(len(windows)>1):
							if(int(windows[1]) == times[1]-1): 
								ok=1
							else: ok=0
				#print("df_nodes_list: ")
				#print(df_nodes_list)
				
				x = "Node output"
				y = "Av. molecule expression/activity"
				hue = "Experiment"
				df_nodes = pd.DataFrame(df_nodes_list, columns=[hue, x, y])
				hue_order=[control_name, exp_name]
				pairs = []
				for gname in output_nodes:
					pairs.append(((gname, control_name), (gname, exp_name)))
					N_tests = N_tests + 1
				axes = sns.barplot(data=df_nodes, x=x, y=y, hue=hue, hue_order=hue_order, seed=2021, ci="sd", edgecolor="black", errcolor="black", errwidth=1.5, capsize = 0.1, alpha=0.5)
				#sns.stripplot(x=x, y=y, hue=hue, data=df_nodes, dodge=True, alpha=0.6, ax=ax)
					
				annot = Annotator(None, pairs)
				annot.new_plot(axes, pairs, plot='barplot', data=df_nodes, x=x, y=y, hue=hue, hue_order=hue_order, seed=2021)
				annot.configure(test='t-test_welch', text_format='star', loc='inside', verbose=2)
				_, test_results = annot.apply_test().annotate()
				
				axes.set_title("\n".join(textwrap.wrap(paper, 30*len(output_nodes))), fontsize = 8)
				plt.text(x=len(output_nodes), y=-0.1, s="\n".join(textwrap.wrap("Control sim.: " + ctr, 100)), fontsize=6, color='gray',rotation=90)
				plt.text(x=len(output_nodes)+0.1, y=-0.1, s="\n".join(textwrap.wrap("Perturb. sim.: " + exp, 100)), fontsize=6, color='gray',rotation=90)
				plt.legend(loc='upper left', bbox_to_anchor=(1.03, 1), title=hue)
				plt.tight_layout()
				
				ii = []
				for result in test_results:
					ii.append(output_nodes.index(result.data.group1[0]))
				
				stat_results = []
				for indnow in ii:
					stat_results.append(test_results[indnow].data.pvalue)
	
	
				dir_ok =[]
	#			print(axes.containers)
				this_line_OK = 1
				some_OK =0
	
				for indnow, gname in enumerate(output_nodes):
					ct_av = axes.containers[0][indnow].get_height()
					exp_av = axes.containers[1][indnow].get_height()
					#print(str(ct_av) + " vs. " + str(exp_av))
					if((exp_av-ct_av > 0) and change_dir == '1'):
						if(stat_results[indnow]<= 5.00e-02 and 2*(exp_av-ct_av)/(exp_av+ct_av) >= p_ch_TH):
							dir_ok.append("Significant")
							some_OK = 1
							tests_OK = tests_OK + 1
						else:
							dir_ok.append("Dir. OK; ns. or < "+ str(p_ch_TH*100) +"% of av. increase")
							this_line_OK = 0
							
					if((exp_av-ct_av > 0) and change_dir == '0'):
						if(stat_results[indnow]<= 5.00e-02 and 2*(exp_av-ct_av)/(exp_av+ct_av) >= p_ch_TH):
							dir_ok.append("Mismatch (opposite & sig.)")
							this_line_OK = 0
						else:
							dir_ok.append("Match (ns. or < "+ str(p_ch_TH*100) +"% of av. in opposite dir.)")
							some_OK = 1
							tests_OK = tests_OK + 1
							
					if((exp_av-ct_av > 0) and change_dir == 'no change'):
						if(stat_results[indnow]<= 5.00e-02 and 2*(exp_av-ct_av)/(exp_av+ct_av) >= p_ch_TH):
							dir_ok.append("Mismatch, increase")
							this_line_OK = 0
						else:
							dir_ok.append("Match (ns. or < "+ str(p_ch_TH*100) +"% of av. increase)")
							some_OK = 1
							tests_OK = tests_OK + 1
	
					if((exp_av-ct_av < 0) and change_dir == '0'):
						if(stat_results[indnow]<= 5.00e-02 and (-2)*(exp_av-ct_av)/(exp_av+ct_av) >= p_ch_TH):
							dir_ok.append("Significant")
							some_OK = 1
							tests_OK = tests_OK + 1
						else:
							dir_ok.append("Dir. OK; ns. or < "+ str(p_ch_TH*100) +"% of av. decrease")
							this_line_OK = 0
							
					if((exp_av-ct_av < 0) and change_dir == '1'):
						if(stat_results[indnow]<= 5.00e-02 and (-2)*(exp_av-ct_av)/(exp_av+ct_av) >= p_ch_TH):
							dir_ok.append("Mismatch (opposite & sig.)")
							this_line_OK = 0
						else:
							dir_ok.append("Mismatch (ns. or < "+ str(p_ch_TH*100) +"% of av. in opposite dir.)")
							this_line_OK = 0
	
					if((exp_av-ct_av < 0) and change_dir == 'no change'):
						if(stat_results[indnow]<= 5.00e-02 and (-2)*(exp_av-ct_av)/(exp_av+ct_av) >= p_ch_TH):
							dir_ok.append("Mismatch, decrease")
							this_line_OK = 0
						else:
							dir_ok.append("Match (ns. or < "+ str(p_ch_TH*100) +"% of av. decrease)")
							some_OK = 1
							tests_OK = tests_OK + 1
							
					if((exp_av-ct_av == 0) and change_dir == 'no change'):
						dir_ok.append("Match (no change)")
						some_OK = 1
						tests_OK = tests_OK + 1
						
					if((exp_av-ct_av == 0) and change_dir == '1'):
						dir_ok.append("Mismatch (inc. not reproduced)")
						this_line_OK = 0
						
					if((exp_av-ct_av == 0) and change_dir == '0'):
						dir_ok.append("Mismatch (dec. not reproduced)")
						this_line_OK = 0
	
				line_OK = line_OK + this_line_OK
				some_in_line_OK = some_in_line_OK + some_OK
				
				#print(change_dir)
				#print(stat_results)
				#print(dir_ok)
				#print(output_nodes)
				bad=0
				for ind in range(len(output_nodes)):
					Stat_tests_output.append([env_st,env_type_str,node_st, pert_type_str, BKGR_str, BKGR_type_str, output_nodes[ind], '', change_dir, dir_ok[ind], stat_results[ind]])
					if(dir_ok[ind] == "Dir. OK; ns. or < "+ str(p_ch_TH*100) +"% of av. increase" or dir_ok[ind] == "Dir. OK; ns. or < "+ str(p_ch_TH*100) +"% of av. decrease"  or dir_ok[ind] == "Mismatch (ns. or < "+ str(p_ch_TH*100) +"% of av. in opposite dir.)" or dir_ok[ind] == "Mismatch (opposite & sig.)" or dir_ok[ind] == "Mismatch, increase" or dir_ok[ind] == "Mismatch, decrease"  or dir_ok[ind] == "Mismatch (inc. not reproduced)"  or dir_ok[ind] == "Mismatch (dec. not reproduced)"):
						BAD_tests_output.append([env_st,env_type_str,node_st, pert_type_str, BKGR_str, BKGR_type_str, output_nodes[ind], '', change_dir, dir_ok[ind], stat_results[ind]])
						bad=1
				if(bad):		
					fail_fig_IDs.append(fig_ID)
				fig_ID =  fig_ID + 1
					
	# 			plt.show()
	# 			exit()	
				
			if(output_phen):
				N_lines = N_lines + 1
				#print(output_phen)
				#print("N_lines = " + str(N_lines))
				
				fig, axes = plt.subplots(1, 1, figsize=(2.5*len(output_phen)+len(exp_name)*0.4, 5), sharex=False, sharey=False)		
	#			fig, axes = plt.subplots(1, 1, figsize=(2.5*len(output_nodes)+len(exp_name)*0.2, 5), sharex=False, sharey=False)		
				df_phen_list = []
				ctr_phen_file = model_folder + "_EXP/General_Time_Series/"+ ctr + "/PhBCh/" + attr_code + "_" + ctr + ".csv"
				#print(ctr_node_file)
				ok=0
				with open(ctr_phen_file, newline='\n') as ct_file:
					CT = csv.reader(ct_file, delimiter=',', quotechar='"')
					#CT_cols = []
					for i in CT:
						#print(i[0])
						#print("ok = " + str(ok))
						if(ok==1 and (str(i[0]) in output_phen)):
							#print("adding line:" + i[0])
							#CT_cols.append(i[0:10])
							for e in i[1:]:
								df_phen_list.append([control_name,i[0],float(e)])
								#print("\t\tadding:" + i[0] + e)
						window=i[0][:-1]
						windows=window.split()
						#print(len(windows), windows)
						if(len(windows)>1):
							if(int(windows[1]) == times[0]-1): 
								ok=1
							else: ok=0
				#print(CT_cols)
				ctr_phen_file = model_folder + "_EXP/General_Time_Series/"+ exp + "/PhBCh/" + attr_code + "_" + exp + ".csv"
				#print(ctr_node_file)
				ok=0
				with open(ctr_phen_file, newline='\n') as ct_file:
					EXP = csv.reader(ct_file, delimiter=',', quotechar='"')
					#EXP_cols = []
					for i in EXP:
						#print(i[0])
						#print(output_phen)
						#print("ok = " + str(ok))
						if(ok==1 and (str(i[0]) in output_phen)):
							#EXP_cols.append(i[0:10])
							#print("adding line:" + i[0])
							for e in i[1:]:
								df_phen_list.append([exp_name,i[0],float(e)])
								#print("\t\tadding:" + i[0] + e)
						window=i[0][:-1]
						windows=window.split()
						#print(len(windows), windows)
						if(len(windows)>1):
							if(int(windows[1]) == times[1]-1): 
								ok=1
							else: ok=0
				#print("df_nodes_list: ")
				#print(df_nodes_list)
				
				x = "Phenotype"
				y = "Average time in phenotype"
				hue = "Experiment"
				df_phen = pd.DataFrame(df_phen_list, columns=[hue, x, y])
				hue_order=[control_name, exp_name]
				pairs = []
				for gname in output_phen:
					pairs.append(((gname, control_name), (gname, exp_name)))
					N_tests = N_tests + 1
	
				axes = sns.barplot(data=df_phen, x=x, y=y, hue=hue, hue_order=hue_order, seed=2021, ci="sd", edgecolor="black", errcolor="black", errwidth=1.5, capsize = 0.1, alpha=0.5, dodge=True)
				
				#sns.stripplot(x=x, y=y, hue=hue, data=df_nodes, dodge=True, alpha=0.6, ax=ax)
				annot = Annotator(None, pairs)
				annot.new_plot(axes, pairs, plot='barplot', data=df_phen, x=x, y=y, hue=hue, hue_order=hue_order, seed=2021)
				annot.configure(test='t-test_welch', text_format='star', loc='inside', verbose=2)
				_, test_results = annot.apply_test().annotate()
	
				#annot.apply_test().annotate()
				axes.set_title("\n".join(textwrap.wrap(paper, 30*len(output_phen))), fontsize = 8)
				plt.text(x=len(output_phen),     y=-0.1, s="\n".join(textwrap.wrap("Control sim.: "  + ctr, 100)), fontsize=6, color='gray',rotation=90)
				plt.text(x=len(output_phen)+0.1, y=-0.1, s="\n".join(textwrap.wrap("Perturb. sim.: " + exp, 100)), fontsize=6, color='gray',rotation=90)
				plt.legend(loc='upper left', bbox_to_anchor=(1.03, 1), title=hue)
				plt.tight_layout()
				#plt.show()
				
				ii = []
				for result in test_results:
					ii.append(output_phen.index(result.data.group1[0]))
				
				stat_results = []
				for indnow in ii:
					stat_results.append(test_results[indnow].data.pvalue)
	
	
				dir_ok =[]
	#			print(axes.containers)
				this_line_OK = 1
				some_OK =0
	
				for indnow, gname in enumerate(output_phen):
					ct_av  = axes.containers[0][indnow].get_height()
					exp_av = axes.containers[1][indnow].get_height()
					#print(str(ct_av) + " vs. " + str(exp_av))
					if((exp_av-ct_av > 0) and change_dir == '1'):
						if(stat_results[indnow]<= 5.00e-02 and 2*(exp_av-ct_av)/(exp_av+ct_av) >= p_ch_TH):
							dir_ok.append("Significant")
							some_OK = 1
							tests_OK = tests_OK + 1
						else:
							dir_ok.append("Dir. OK; ns. or < "+ str(p_ch_TH*100) +"% of av. increase")
							this_line_OK = 0
							
					if((exp_av-ct_av > 0) and change_dir == '0'):
						if(stat_results[indnow]<= 5.00e-02 and 2*(exp_av-ct_av)/(exp_av+ct_av) >= p_ch_TH):
							dir_ok.append("Mismatch (opposite & sig.)")
							this_line_OK = 0
						else:
							dir_ok.append("Match (ns. or < "+ str(p_ch_TH*100) +"% of av. in opposite dir.)")
							some_OK = 1
							tests_OK = tests_OK + 1
							
					if((exp_av-ct_av > 0) and change_dir == 'no change'):
						if(stat_results[indnow]<= 5.00e-02 and 2*(exp_av-ct_av)/(exp_av+ct_av) >= p_ch_TH):
							dir_ok.append("Mismatch, increase")
							this_line_OK = 0
						else:
							dir_ok.append("Match (ns. or < "+ str(p_ch_TH*100) +"% of av. increase)")
							some_OK = 1
							tests_OK = tests_OK + 1
	
					if((exp_av-ct_av < 0) and change_dir == '0'):
						if(stat_results[indnow]<= 5.00e-02 and (-2)*(exp_av-ct_av)/(exp_av+ct_av) >= p_ch_TH):
							dir_ok.append("Significant")
							some_OK = 1
							tests_OK = tests_OK + 1
						else:
							dir_ok.append("Dir. OK; ns. or < "+ str(p_ch_TH*100) +"% of av. decrease")
							this_line_OK = 0
							
					if((exp_av-ct_av < 0) and change_dir == '1'):
						if(stat_results[indnow]<= 5.00e-02 and (-2)*(exp_av-ct_av)/(exp_av+ct_av) >= p_ch_TH):
							dir_ok.append("Mismatch (opposite & sig.)")
							this_line_OK = 0
						else:
							dir_ok.append("Mismatch (ns. or < "+ str(p_ch_TH*100) +"% of av. in opposite dir.)")
							this_line_OK = 0
	
					if((exp_av-ct_av < 0) and change_dir == 'no change'):
						if(stat_results[indnow]<= 5.00e-02 and (-2)*(exp_av-ct_av)/(exp_av+ct_av) >= p_ch_TH):
							dir_ok.append("Mismatch, decrease")
							this_line_OK = 0
						else:
							dir_ok.append("Match (ns. or < "+ str(p_ch_TH*100) +"% of av. decrease)")
							some_OK = 1
							tests_OK = tests_OK + 1
							
					if((exp_av-ct_av == 0) and change_dir == 'no change'):
						dir_ok.append("Match (no change)")
						some_OK = 1
						tests_OK = tests_OK + 1
						
					if((exp_av-ct_av == 0) and change_dir == '1'):
						dir_ok.append("Mismatch (inc. not reproduced)")
						this_line_OK = 0
						
					if((exp_av-ct_av == 0) and change_dir == '0'):
						dir_ok.append("Mismatch (dec. not reproduced)")
						this_line_OK = 0
	
				line_OK = line_OK + this_line_OK
				some_in_line_OK = some_in_line_OK + some_OK
				
				#print(change_dir)
				#print(stat_results)
				#print(dir_ok)
				#print(output_phen)
				bad=0
				for ind in range(len(output_phen)):
					Stat_tests_output.append([env_st, env_type_str,node_st, pert_type_str, BKGR_str, BKGR_type_str, output_phen[ind], '', change_dir, dir_ok[ind], stat_results[ind]])
					if(dir_ok[ind] == "Dir. OK; ns. or < "+ str(p_ch_TH*100) +"% of av. increase" or dir_ok[ind] == "Dir. OK; ns. or < "+ str(p_ch_TH*100) +"% of av. decrease"  or dir_ok[ind] == "Mismatch (ns. or < "+ str(p_ch_TH*100) +"% of av. in opposite dir.)" or dir_ok[ind] == "Mismatch (opposite & sig.)" or dir_ok[ind] == "Mismatch, increase" or dir_ok[ind] == "Mismatch, decrease"  or dir_ok[ind] == "Mismatch (inc. not reproduced)"  or dir_ok[ind] == "Mismatch (dec. not reproduced)"):
						BAD_tests_output.append([env_st, env_type_str,node_st, pert_type_str, BKGR_str, BKGR_type_str, output_phen[ind], '', change_dir, dir_ok[ind], stat_results[ind]])
						bad=1
				if(bad):		
					fail_fig_IDs.append(fig_ID)
				fig_ID =  fig_ID + 1
	

def save_multi_image(filename):
	pp = PdfPages(filename)
	fig_nums = plt.get_fignums()
	figs = [plt.figure(n) for n in fig_nums]
	for fig in figs:
		fig.savefig(pp, format='pdf')
	pp.close()


def save_multi_image_failed_tests(filename):
	pp = PdfPages(filename)
	#fig_nums = plt.get_fignums()
	#print(fail_fig_IDs)
	figs = [plt.figure(n+1) for n in fail_fig_IDs]
	for fig in figs:
		fig.savefig(pp, format='pdf')
	pp.close()

save_multi_image(model_folder + "Suppl File 4 - Hypoxia Model Validation Figures.pdf")
save_multi_image_failed_tests(model_folder + "Failed Validation Figures.pdf")
df = pd.DataFrame(Stat_tests_output)
print(df.to_string())
df.to_csv('Hypoxia Model Validation Statistics.csv')

print("\n\nLines that match for all output nodes/phenotypes: " + str(line_OK) + " of " + str(N_lines) + " => " + str("%.2f" % float(line_OK/N_lines))+ "% match.")
print("Lines in which at least one output node/phenotype matches: " + str(some_in_line_OK) + " of " + str(N_lines) + " => " + str("%.2f" % float(some_in_line_OK/N_lines))+ "% match.")	
print("Number of unique statistical tests that match: " + str(tests_OK) + " of " + str(N_tests) + " => " + str("%.2f" % float(tests_OK/N_tests))+ "% match.")	

print("\n\nFailed tests:")
df2 = pd.DataFrame(BAD_tests_output)
print(df2.to_string())
print(fail_fig_IDs)
```
