## Supplementary File 3 for "A Boolean network model of hypoxia, mechanosensing and TGF-β signaling captures the role of phenotypic plasticity and mutations in tumor metastasis"

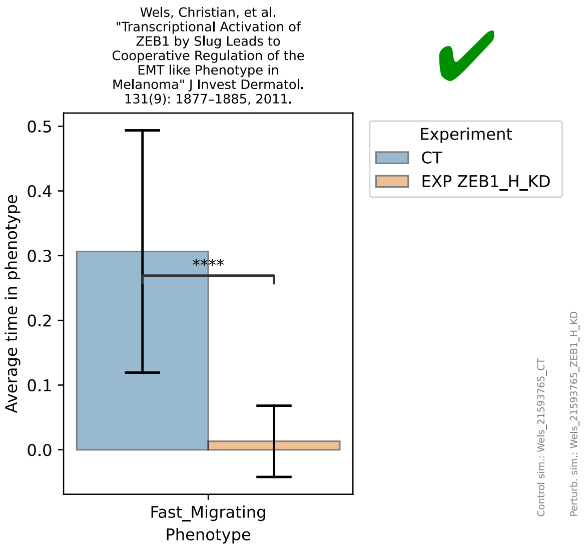
Wels, Christian, et al. "Transcriptional Activation of ZEB1 by Slug Leads to Cooperative Regulation of the EMT like Phenotype in Melanoma" J Invest Dermatol. 131(9): 1877–1885, 2011.
Experiment
CT
EXP ZEB1_H_KD
Average time in phenotype
****
Fast_Migrating
Phenotype
Control sim.: Wels_21593765_CT
Perturb. sim.: Wels_21593765_ZEB1_H_KD

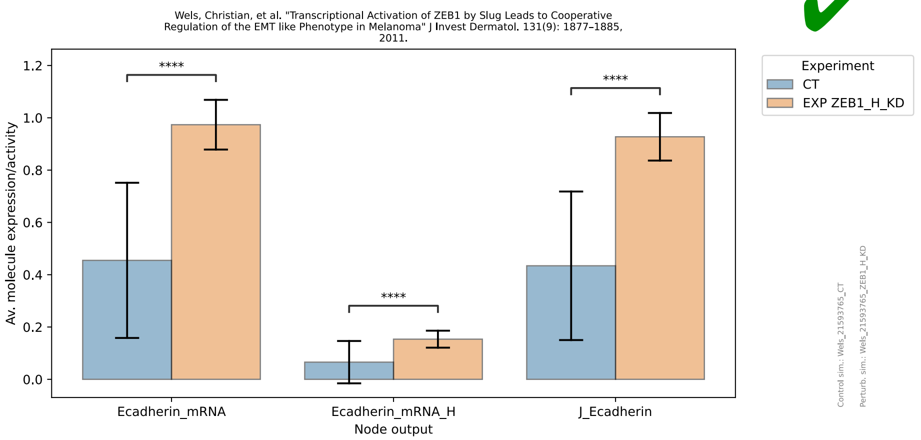
Wels, Christian, et al. "Transcriptional Activation of ZEB1 by Slug Leads to Cooperative Regulation of the EMT like Phenotype in Melanoma" J Invest Dermatol. 131(9): 1877–1885, 2011.
Experiment
CT
EXP ZEB1_H_KD
Av. molecule expression/activity
****
Ecadherin_mRNA
Ecadherin_mRNA_H
J_Ecadherin
Node output
Control sim.: Wels_21593765_CT
Perturb. sim.: Wels_21593765_ZEB1_H_KD

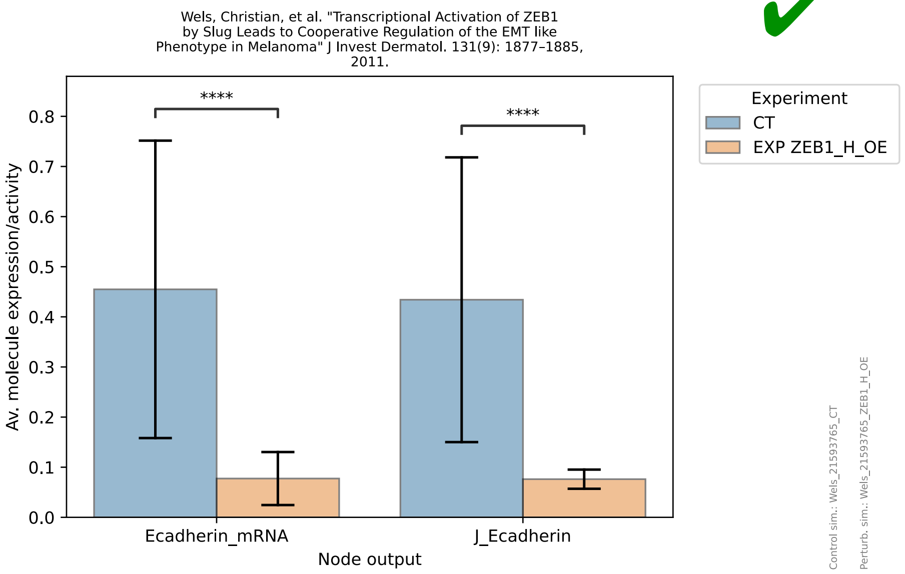
Wels, Christian, et al. "Transcriptional Activation of ZEB1 by Slug Leads to Cooperative Regulation of the EMT like Phenotype in Melanoma" J Invest Dermatol. 131(9): 1877–1885, 2011.
Experiment
CT
EXP ZEB1_H_OE
Av. molecule expression/activity
****
Ecadherin_mRNA
J_Ecadherin
Node output
Control sim.: Wels_21593765_CT
Perturb. sim.: Wels_21593765_ZEB1_H_OE

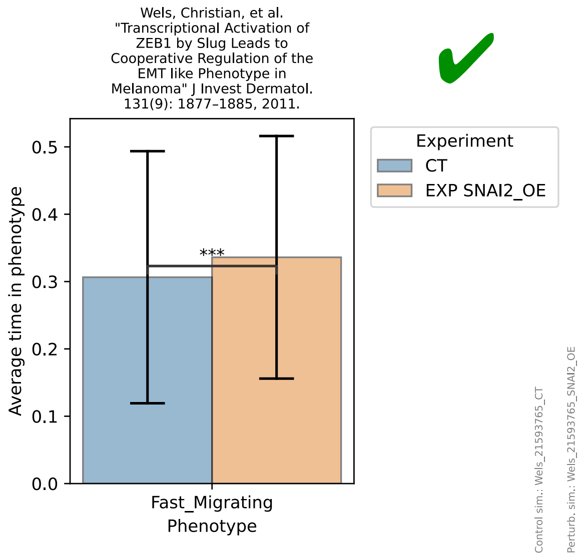
Wels, Christian, et al. "Transcriptional Activation of ZEB1 by Slug Leads to Cooperative Regulation of the EMT like Phenotype in Melanoma" J Invest Dermatol. 131(9): 1877–1885, 2011.
Experiment
CT
EXP SNAI2_OE
Average time in phenotype
***
Fast_Migrating
Phenotype
Control sim.: Wels_21593765_CT
Perturb. sim.: Wels_21593765_SNAI2_OE

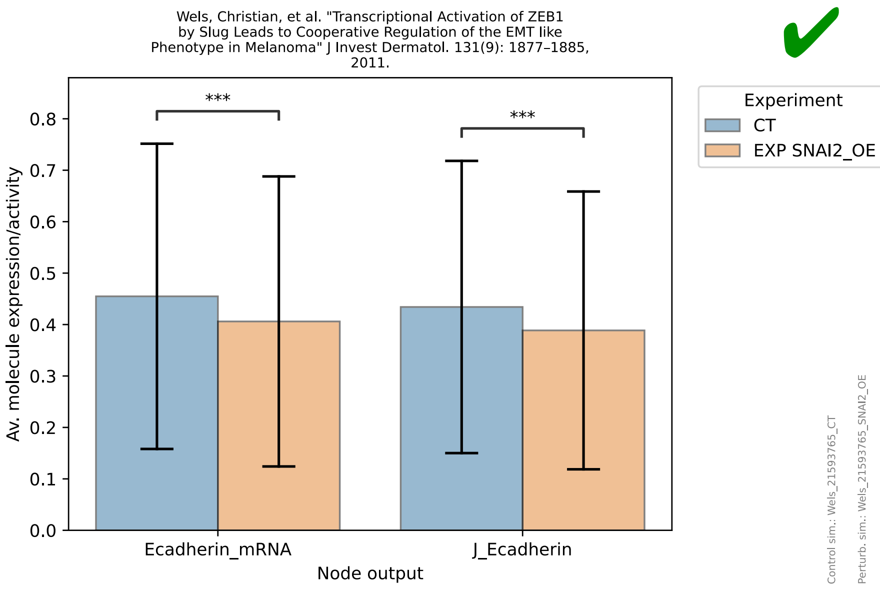
Wels, Christian, et al. "Transcriptional Activation of ZEB1 by Slug Leads to Cooperative Regulation of the EMT like Phenotype in Melanoma" J Invest Dermatol. 131(9): 1877–1885, 2011.
Av. molecule expression/activity
0.0
0.1
0.2
0.3
0.4
0.5
0.6
0.7
0.8
***
***
Ecadherin_mRNA
J_Ecadherin
Node output
Experiment
CT
EXP SNAI2_OE
Control sim.: Wels_21593765_CT
Perturb. sim.: Wels_21593765_SNAI2_OE

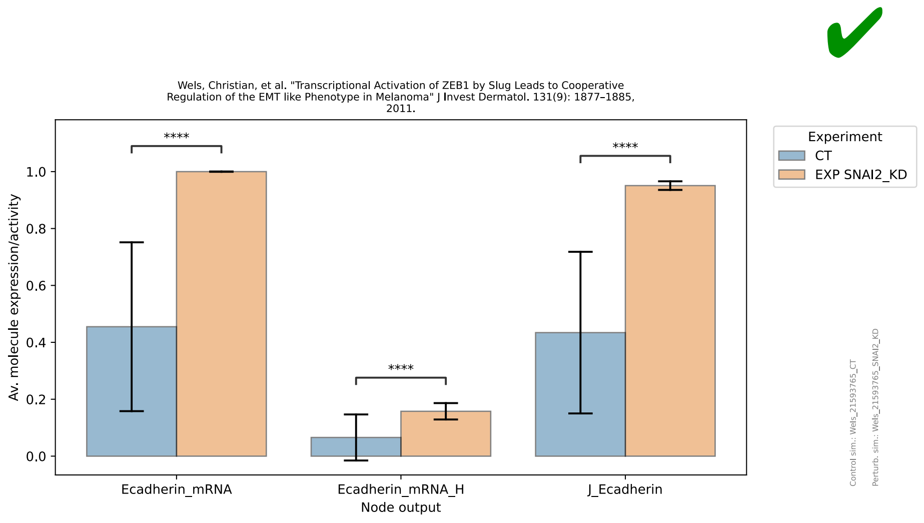
Wels, Christian, et al. "Transcriptional Activation of ZEB1 by Slug Leads to Cooperative Regulation of the EMT like Phenotype in Melanoma" J Invest Dermatol. 131(9): 1877–1885, 2011.
Av. molecule expression/activity
0.0
0.2
0.4
0.6
0.8
1.0
****
****
****
Ecadherin_mRNA
Ecadherin_mRNA_H
J_Ecadherin
Node output
Experiment
CT
EXP SNAI2_KD
Control sim.: Wels_21593765_CT
Perturb. sim.: Wels_21593765_SNAI2_KD

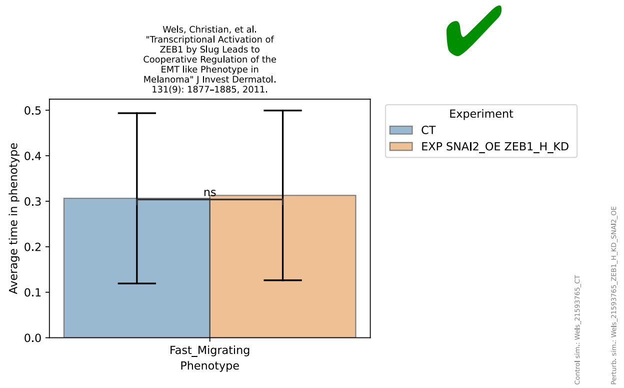
Wels, Christian, et al. "Transcriptional Activation of ZEB1 by Slug Leads to Cooperative Regulation of the EMT like Phenotype in Melanoma" J Invest Dermatol. 131(9): 1877–1885, 2011.
Average time in phenotype
0.0
0.1
0.2
0.3
0.4
0.5
ns
Fast_Migrating
Phenotype
Experiment
CT
EXP SNAI2_OE ZEB1_H_KD
Control sim.: Wels_21593765_CT
Perturb. sim.: Wels_21593765_ZEB1_H_KD_SNAI2_OE

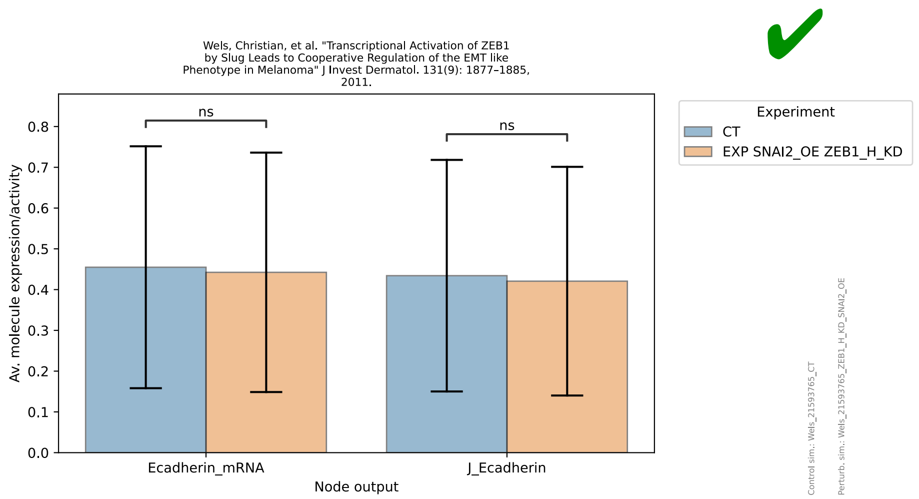
Wels, Christian, et al. "Transcriptional Activation of ZEB1 by Slug Leads to Cooperative Regulation of the EMT like Phenotype in Melanoma" J Invest Dermatol. 131(9): 1877–1885, 2011.
Av. molecule expression/activity
0.0
0.1
0.2
0.3
0.4
0.5
0.6
0.7
0.8
ns
ns
Ecadherin_mRNA
J_Ecadherin
Node output
Experiment
CT
EXP SNAI2_OE ZEB1_H_KD
Control sim.: Wels_21593765_CT
Perturb. sim.: Wels_21593765_ZEB1_H_KD_SNAI2_OE

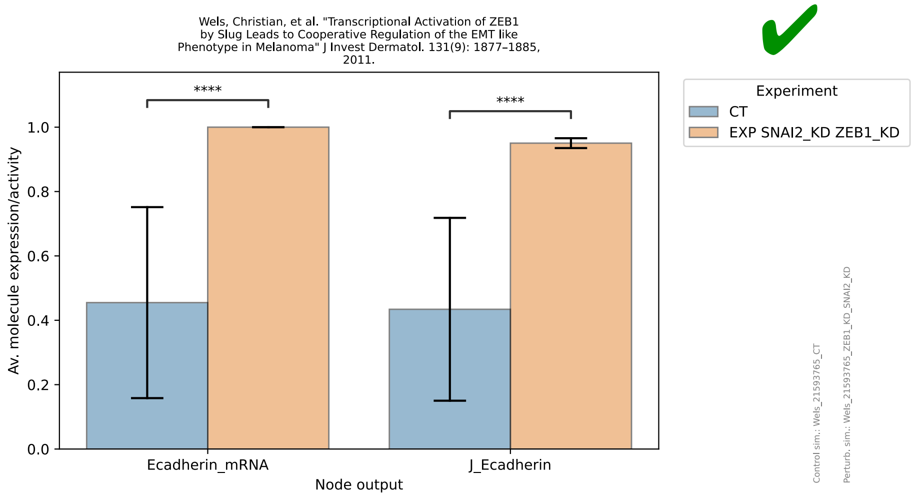
Wels, Christian, et al. "Transcriptional Activation of ZEB1 by Slug Leads to Cooperative Regulation of the EMT like Phenotype in Melanoma" J Invest Dermatol. 131(9): 1877–1885, 2011.
Av. molecule expression/activity
Ecadherin_mRNA
J_Ecadherin
Node output
****
****
Experiment
CT
EXP SNAI2_KD ZEB1_KD
Control sim.: Wels_21593765_CT
Perturb. sim.: Wels_21593765_ZEB1_KD_SNAI2_KD

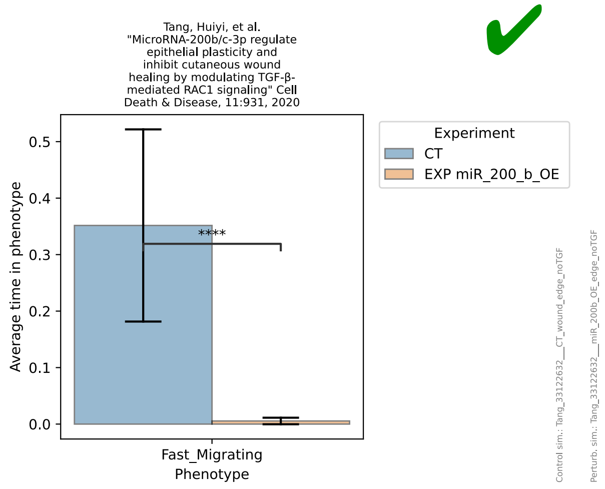
Tang, Huiyi, et al. "MicroRNA-200b/c-3p regulate epithelial plasticity and inhibit cutaneous wound healing by modulating TGF-β-mediated RAC1 signaling" Cell Death & Disease, 11:931, 2020
Average time in phenotype
Fast_Migrating
Phenotype
****
Experiment
CT
EXP miR_200_b_OE
Control sim.: Tang_33122632___CT_wound_edge_noTGF
Perturb. sim.: Tang_33122632___miR_200b_OE_edge_noTGF

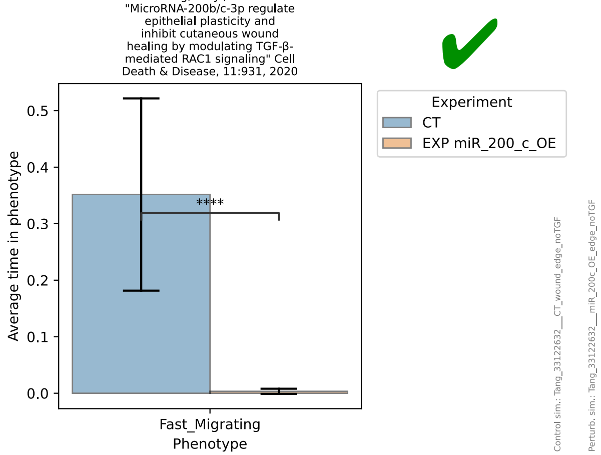
Tang, Huiyi, et al. "MicroRNA-200b/c-3p regulate epithelial plasticity and inhibit cutaneous wound healing by modulating TGF-β-mediated RAC1 signaling" Cell Death & Disease, 11:931, 2020
Average time in phenotype
Fast_Migrating
Phenotype
****
Experiment
CT
EXP miR_200_c_OE
Control sim.: Tang_33122632___CT_wound_edge_noTGF
Perturb. sim.: Tang_33122632___miR_200c_OE_edge_noTGF

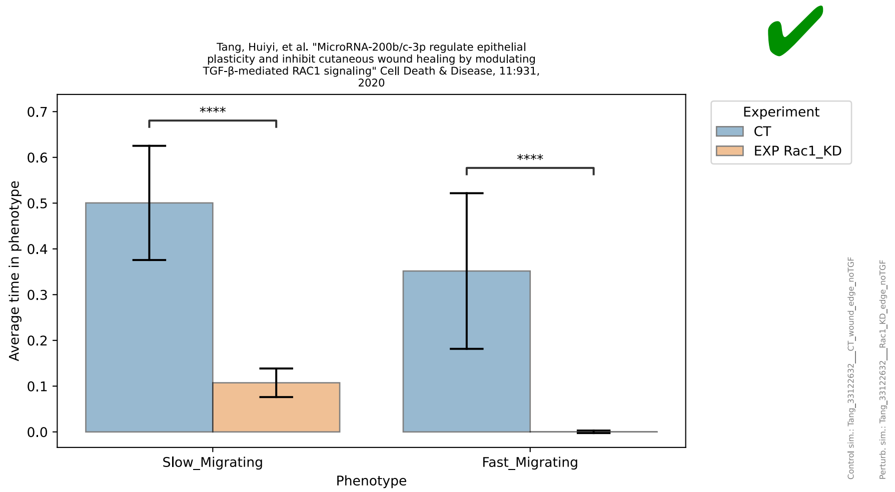
Tang, Huiyi, et al. "MicroRNA-200b/c-3p regulate epithelial plasticity and inhibit cutaneous wound healing by modulating TGF-β-mediated RAC1 signaling" Cell Death & Disease, 11:931, 2020
Average time in phenotype
Slow_Migrating
Fast_Migrating
Phenotype
****
****
Experiment
CT
EXP Rac1_KD
Control sim.: Tang_33122632___CT_wound_edge_noTGF
Perturb. sim.: Tang_33122632___Rac1_KD_edge_noTGF

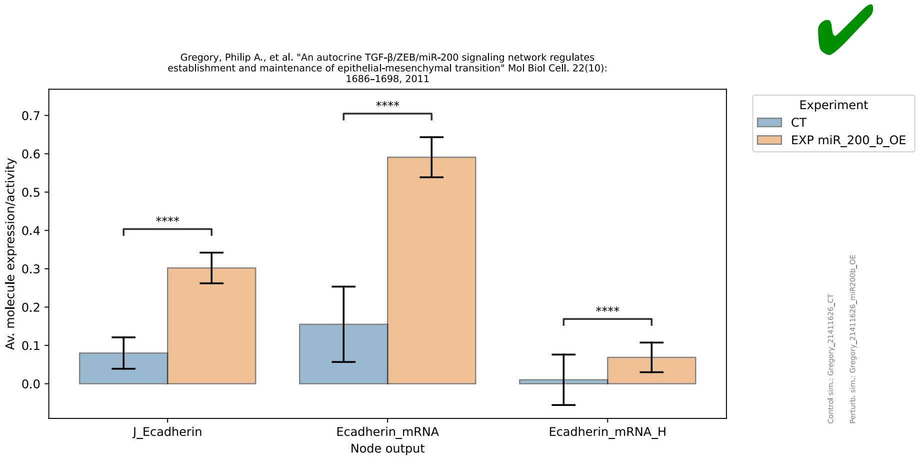
Gregory, Philip A., et al. "An autocrine TGF-β/ZEB/miR-200 signaling network regulates establishment and maintenance of epithelial-mesenchymal transition" Mol Biol Cell. 22(10): 1686–1698, 2011
Av. molecule expression/activity
Node output
J_Ecadherin
Ecadherin_mRNA
Ecadherin_mRNA_H
****
Experiment
CT
EXP miR_200_b_OE
Control sim.: Gregory_21411626_CT
Perturb. sim.: Gregory_21411626_miR200b_OE

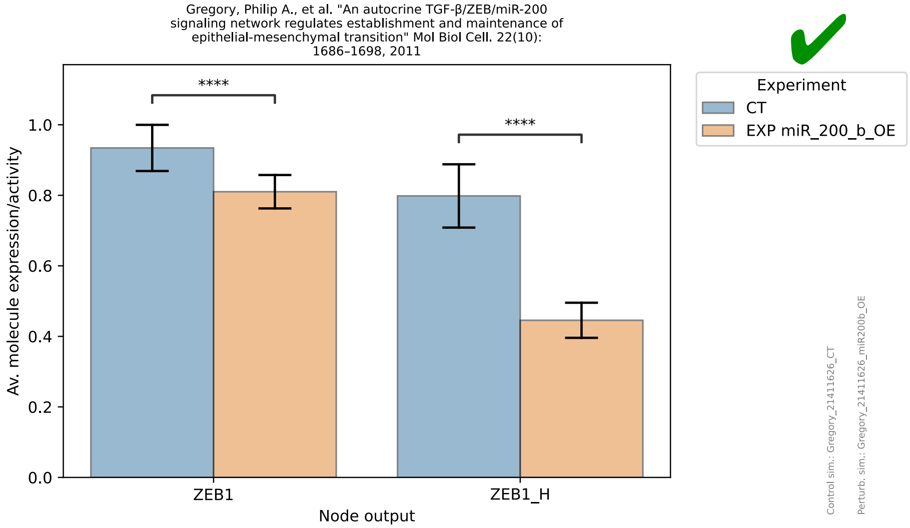
Gregory, Philip A., et al. "An autocrine TGF-β/ZEB/miR-200 signaling network regulates establishment and maintenance of epithelial-mesenchymal transition" Mol Biol Cell. 22(10): 1686–1698, 2011
Av. molecule expression/activity
Node output
ZEB1
ZEB1_H
****
Experiment
CT
EXP miR_200_b_OE
Control sim.: Gregory_21411626_CT
Perturb. sim.: Gregory_21411626_miR200b_OE

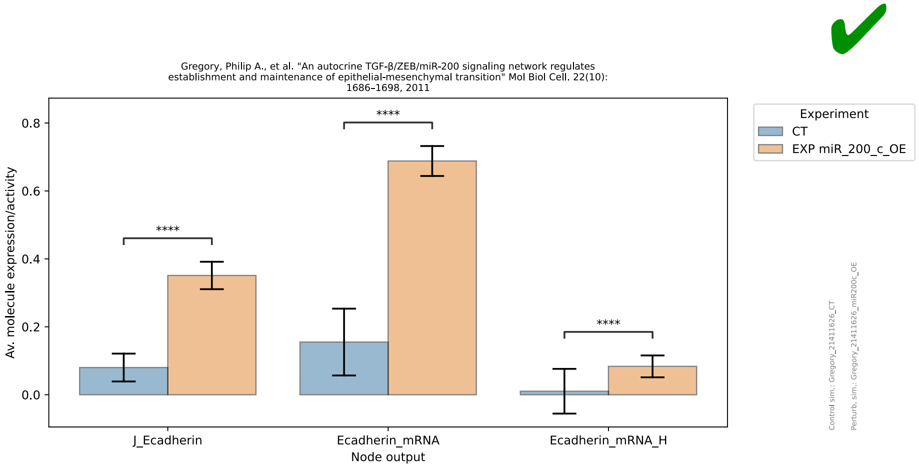
Gregory, Philip A., et al. "An autocrine TGF-β/ZEB/miR-200 signaling network regulates establishment and maintenance of epithelial-mesenchymal transition" Mol Biol Cell. 22(10): 1686–1698, 2011
Av. molecule expression/activity
Node output
J_Ecadherin
Ecadherin_mRNA
Ecadherin_mRNA_H
****
Experiment
CT
EXP miR_200_c_OE
Control sim.: Gregory_21411626_CT
Perturb. sim.: Gregory_21411626_miR200c_OE

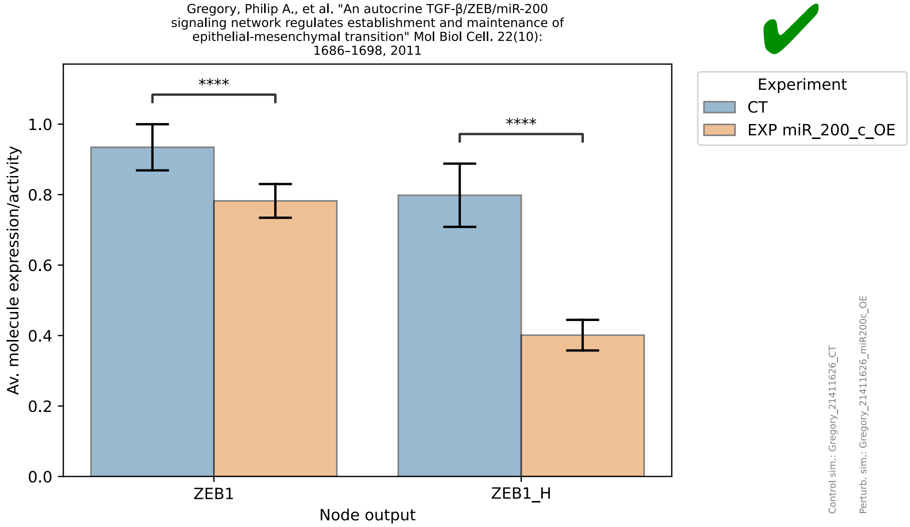
Gregory, Philip A., et al. "An autocrine TGF-β/ZEB/miR-200 signaling network regulates establishment and maintenance of epithelial-mesenchymal transition" Mol Biol Cell. 22(10): 1686–1698, 2011
Av. molecule expression/activity
Node output
ZEB1
ZEB1_H
****
Experiment
CT
EXP miR_200_c_OE
Control sim.: Gregory_21411626_CT
Perturb. sim.: Gregory_21411626_miR200c_OE

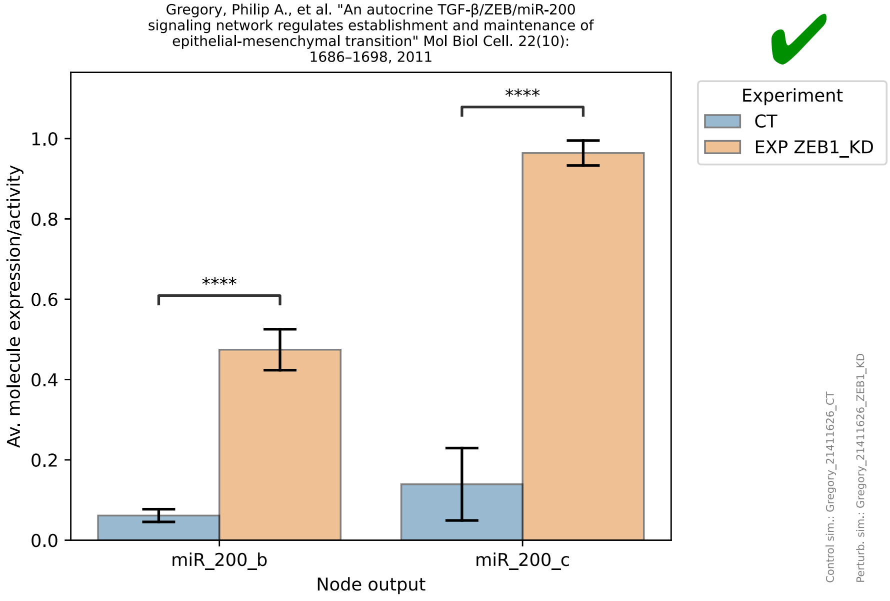
Gregory, Philip A., et al. "An autocrine TGF-β/ZEB/miR-200 signaling network regulates establishment and maintenance of epithelial-mesenchymal transition" Mol Biol Cell. 22(10): 1686–1698, 2011
Av. molecule expression/activity
Node output
miR_200_b
miR_200_c
****
****
Experiment
CT
EXP ZEB1_KD
Control sim.: Gregory_21411626_CT
Perturb. sim.: Gregory_21411626_ZEB1_KD

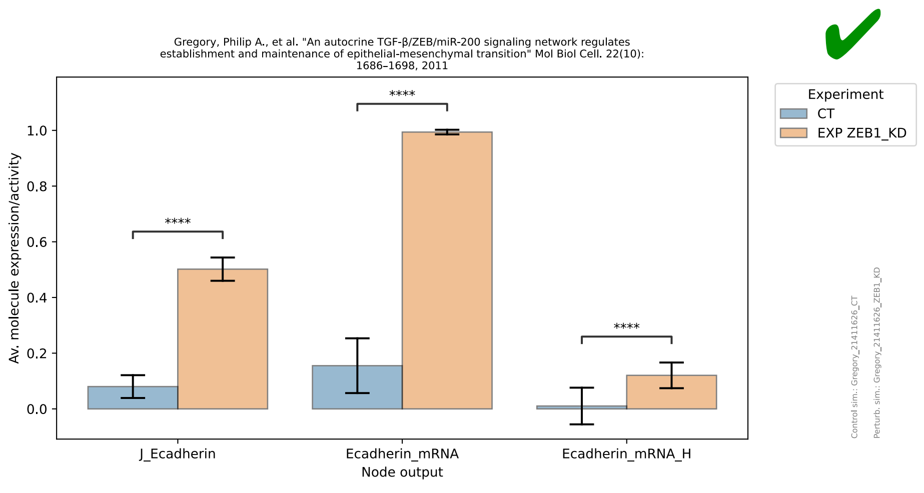
Gregory, Philip A., et al. "An autocrine TGF-β/ZEB/miR-200 signaling network regulates establishment and maintenance of epithelial-mesenchymal transition" Mol Biol Cell. 22(10): 1686–1698, 2011
Av. molecule expression/activity
Node output
J_Ecadherin
Ecadherin_mRNA
Ecadherin_mRNA_H
****
****
****
Experiment
CT
EXP ZEB1_KD
Control sim.: Gregory_21411626_CT
Perturb. sim.: Gregory_21411626_ZEB1_KD

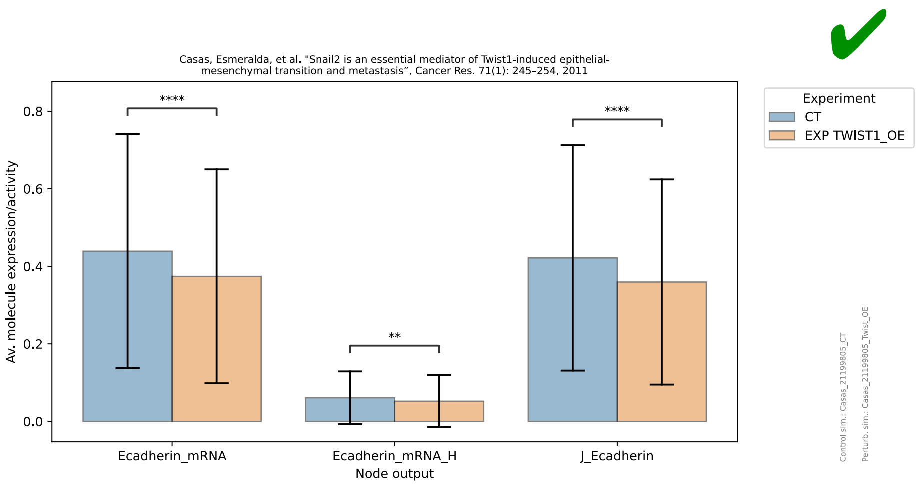
Casas, Esmeralda, et al. "Snail2 is an essential mediator of Twist1-induced epithelial-mesenchymal transition and metastasis", Cancer Res. 71(1): 245–254, 2011
Av. molecule expression/activity
Node output
Ecadherin_mRNA
Ecadherin_mRNA_H
J_Ecadherin
****
**
****
Experiment
CT
EXP TWIST1_OE
Control sim.: Casas_21199805_CT
Perturb. sim.: Casas_21199805_Twist_OE

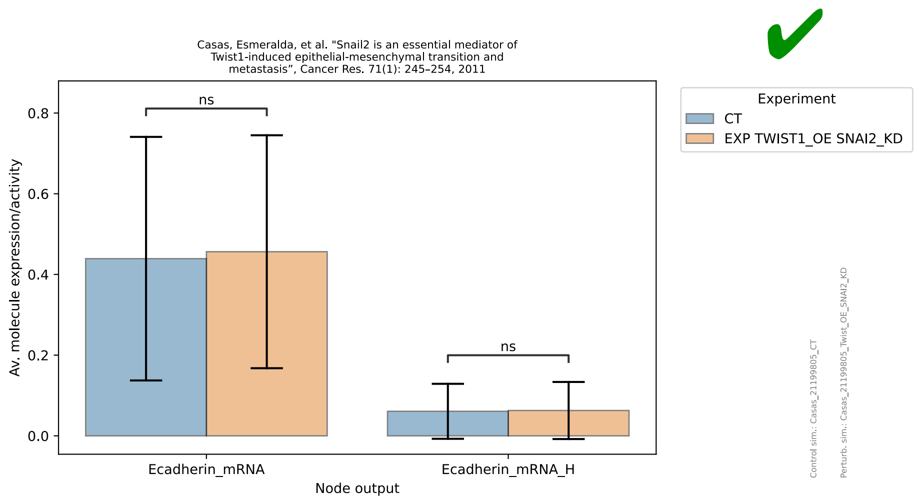
Casas, Esmeralda, et al. "Snail2 is an essential mediator of Twist1-induced epithelial-mesenchymal transition and metastasis", Cancer Res. 71(1): 245–254, 2011
Av. molecule expression/activity
Node output
Ecadherin_mRNA
Ecadherin_mRNA_H
ns
ns
Experiment
CT
EXP TWIST1_OE SNAI2_KD
Control sim.: Casas_21199805_CT
Perturb. sim.: Casas_21199805_Twist_OE_SNAI2_KD

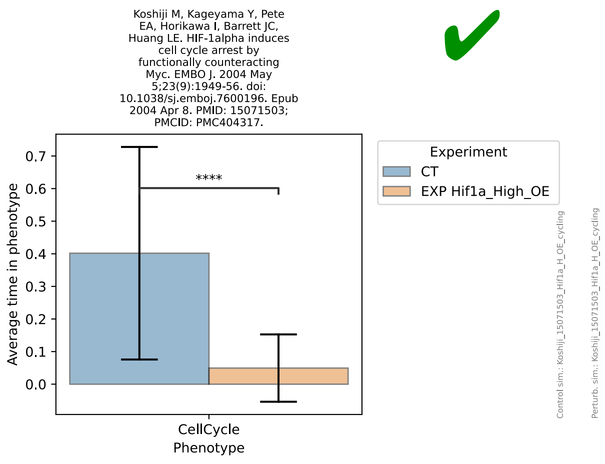
Koshiji M, Kageyama Y, Pete EA, Horikawa I, Barrett JC, Huang LE. HIF-1alpha induces cell cycle arrest by functionally counteracting Myc. EMBO J. 2004 May 5;23(9):1949-56. doi: 10.1038/sj.emboj.7600196. Epub 2004 Apr 8. PMID: 15071503; PMCID: PMC404317.
✔
Average time in phenotype
0.0
0.1
0.2
0.3
0.4
0.5
0.6
0.7
****
CellCycle
Phenotype
Experiment
CT
EXP Hif1a_High_OE
Control sim.: Koshiji_15071503_Hif1a_H_OE_cycling
Perturb. sim.: Koshiji_15071503_Hif1a_H_OE_cycling

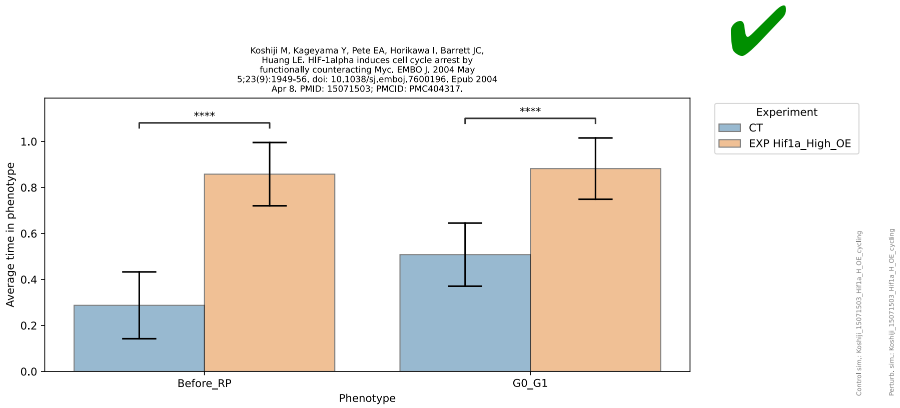
Koshiji M, Kageyama Y, Pete EA, Horikawa I, Barrett JC, Huang LE. HIF-1alpha induces cell cycle arrest by functionally counteracting Myc. EMBO J. 2004 May 5;23(9):1949-56. doi: 10.1038/sj.emboj.7600196. Epub 2004 Apr 8. PMID: 15071503; PMCID: PMC404317.
✔
Average time in phenotype
0.0
0.2
0.4
0.6
0.8
1.0
****
****
Before_RP
G0_G1
Phenotype
Experiment
CT
EXP Hif1a_High_OE
Control sim.: Koshiji_15071503_Hif1a_H_OE_cycling
Perturb. sim.: Koshiji_15071503_Hif1a_H_OE_cycling

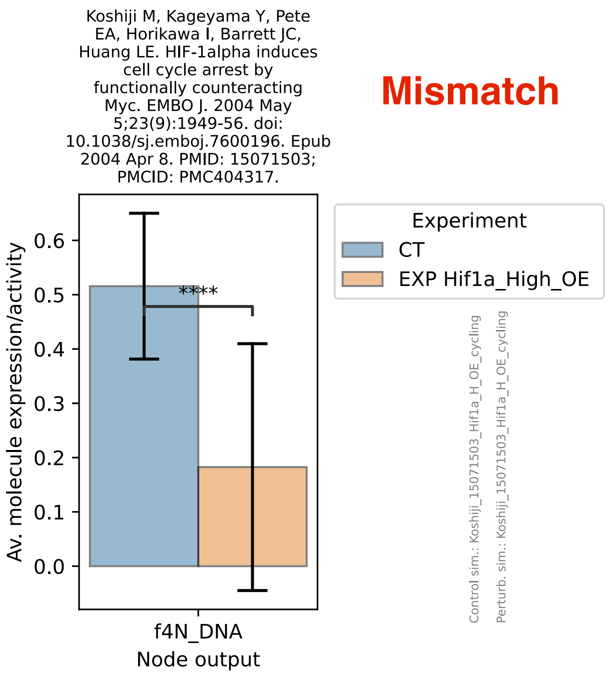
Koshiji M, Kageyama Y, Pete EA, Horikawa I, Barrett JC, Huang LE. HIF-1alpha induces cell cycle arrest by functionally counteracting Myc. EMBO J. 2004 May 5;23(9):1949-56. doi: 10.1038/sj.emboj.7600196. Epub 2004 Apr 8. PMID: 15071503; PMCID: PMC404317.
Mismatch
Av. molecule expression/activity
0.0
0.1
0.2
0.3
0.4
0.5
0.6
****
f4N_DNA
Node output
Experiment
CT
EXP Hif1a_High_OE
Control sim.: Koshiji_15071503_Hif1a_H_OE_cycling
Perturb. sim.: Koshiji_15071503_Hif1a_H_OE_cycling

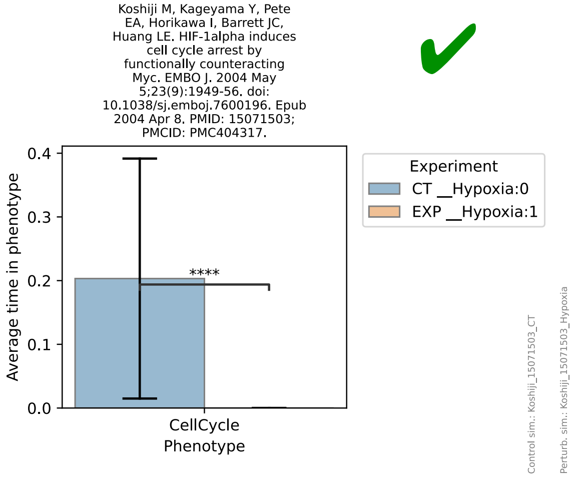
Koshiji M, Kageyama Y, Pete EA, Horikawa I, Barrett JC, Huang LE. HIF-1alpha induces cell cycle arrest by functionally counteracting Myc. EMBO J. 2004 May 5;23(9):1949-56. doi: 10.1038/sj.emboj.7600196. Epub 2004 Apr 8. PMID: 15071503; PMCID: PMC404317.
✔
Average time in phenotype
0.0
0.1
0.2
0.3
0.4
****
CellCycle
Phenotype
Experiment
CT __Hypoxia:0
EXP __Hypoxia:1
Control sim.: Koshiji_15071503_CT
Perturb. sim.: Koshiji_15071503_Hypoxia

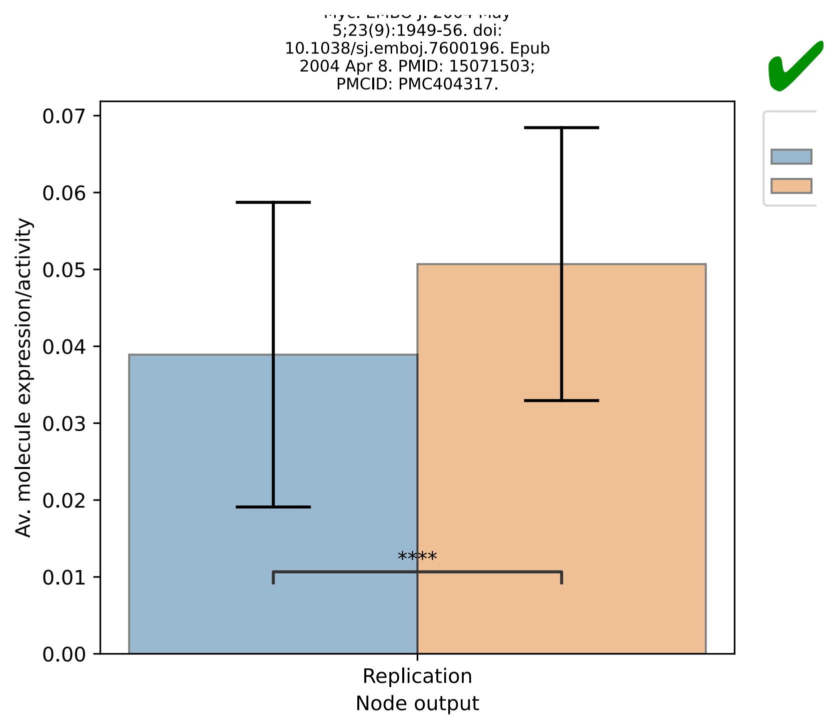
5;23(9):1949-56. doi:
10.1038/sj.emboj.7600196. Epub
2004 Apr 8. PMID: 15071503;
PMCID: PMC404317.
Av. molecule expression/activity
0.07
0.06
0.05
0.04
0.03
0.02
0.01
0.00
****
Replication
Node output

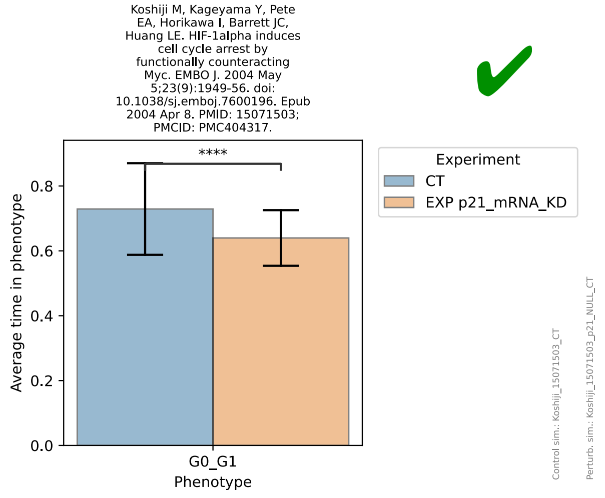
Koshiji M, Kageyama Y, Pete
EA, Horikawa I, Barrett JC,
Huang LE. HIF-1alpha induces
cell cycle arrest by
functionally counteracting
Myc. EMBO J. 2004 May
5;23(9):1949-56. doi:
10.1038/sj.emboj.7600196. Epub
2004 Apr 8. PMID: 15071503;
PMCID: PMC404317.
Experiment
CT
EXP p21_mRNA_KD
Average time in phenotype
0.8
0.6
0.4
0.2
0.0
****
G0_G1
Phenotype
Control sim.: Koshiji_15071503_CT
Perturb. sim.: Koshiji_15071503_p21_NULL_CT

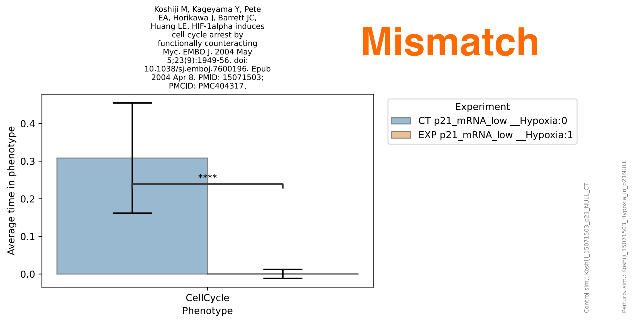
Koshiji M, Kageyama Y, Pete
EA, Horikawa I, Barrett JC,
Huang LE. HIF-1alpha induces
cell cycle arrest by
functionally counteracting
Myc. EMBO J. 2004 May
5;23(9):1949-56. doi:
10.1038/sj.emboj.7600196. Epub
2004 Apr 8. PMID: 15071503;
PMCID: PMC404317.
Mismatch
Experiment
CT p21_mRNA_low __Hypoxia:0
EXP p21_mRNA_low __Hypoxia:1
Average time in phenotype
0.4
0.3
0.2
0.1
0.0
****
CellCycle
Phenotype
Control sim.: Koshiji_15071503_p21_NULL_CT
Perturb. sim.: Koshiji_15071503_Hypoxia_in_p21NULL

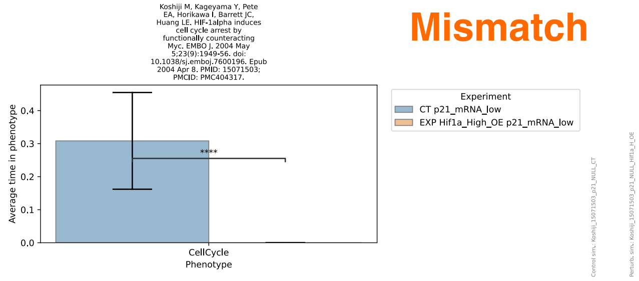
Koshiji M, Kageyama Y, Pete
EA, Horikawa I, Barrett JC,
Huang LE. HIF-1alpha induces
cell cycle arrest by
functionally counteracting
Myc. EMBO J. 2004 May
5;23(9):1949-56. doi:
10.1038/sj.emboj.7600196. Epub
2004 Apr 8. PMID: 15071503;
PMCID: PMC404317.
Mismatch
Experiment
CT p21_mRNA_low
EXP Hif1a_High_OE p21_mRNA_low
Average time in phenotype
0.4
0.3
0.2
0.1
0.0
****
CellCycle
Phenotype
Control sim.: Koshiji_15071503_p21_NULL_CT
Perturb. sim.: Koshiji_15071503_p21_NULL_Hif1a_H_OE

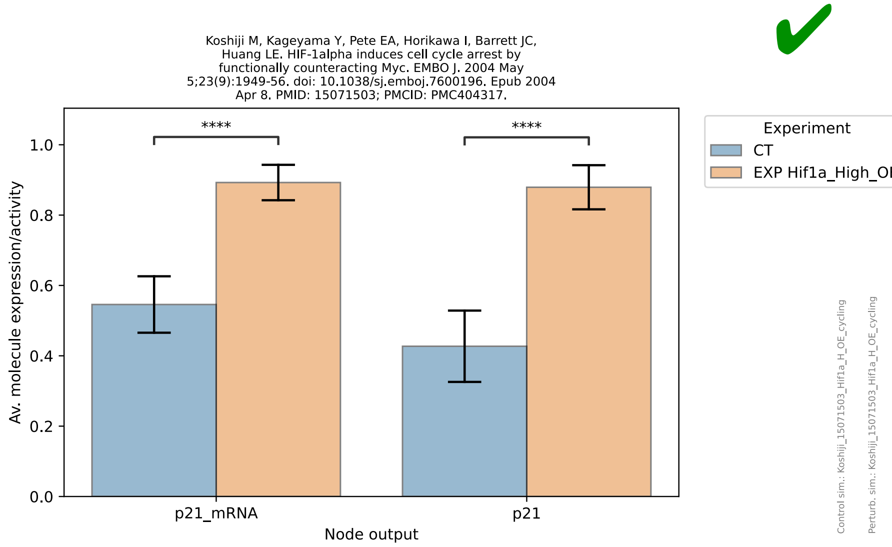
Koshiji M, Kageyama Y, Pete EA, Horikawa I, Barrett JC, Huang LE. HIF-1alpha induces cell cycle arrest by functionally counteracting Myc. EMBO J. 2004 May 5;23(9):1949-56. doi: 10.1038/sj.emboj.7600196. Epub 2004 Apr 8. PMID: 15071503; PMCID: PMC404317.
Av. molecule expression/activity
Node output
p21_mRNA
p21
****
****
Experiment
CT
EXP Hif1a_High_OE
Control sim.: Koshiji_15071503_Hif1a_H_OE_cycling
Perturb. sim.: Koshiji_15071503_Hif1a_H_OE_cycling

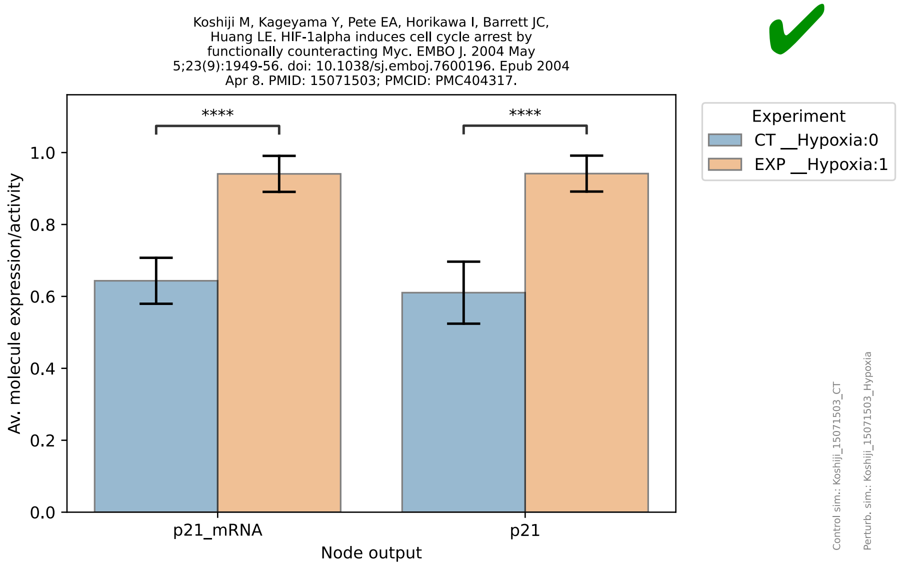
Koshiji M, Kageyama Y, Pete EA, Horikawa I, Barrett JC, Huang LE. HIF-1alpha induces cell cycle arrest by functionally counteracting Myc. EMBO J. 2004 May 5;23(9):1949-56. doi: 10.1038/sj.emboj.7600196. Epub 2004 Apr 8. PMID: 15071503; PMCID: PMC404317.
Av. molecule expression/activity
Node output
p21_mRNA
p21
****
****
Experiment
CT __Hypoxia:0
EXP __Hypoxia:1
Control sim.: Koshiji_15071503_CT
Perturb. sim.: Koshiji_15071503_Hypoxia

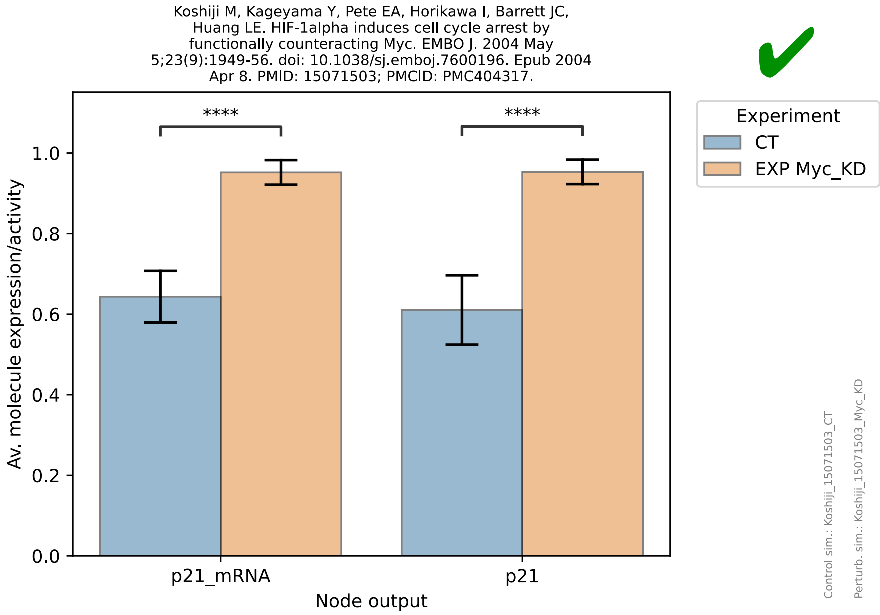
Koshiji M, Kageyama Y, Pete EA, Horikawa I, Barrett JC, Huang LE. HIF-1alpha induces cell cycle arrest by functionally counteracting Myc. EMBO J. 2004 May 5;23(9):1949-56. doi: 10.1038/sj.emboj.7600196. Epub 2004 Apr 8. PMID: 15071503; PMCID: PMC404317.
Av. molecule expression/activity
Node output
p21_mRNA
p21
****
****
Experiment
CT
EXP Myc_KD
Control sim.: Koshiji_15071503_CT
Perturb. sim.: Koshiji_15071503_Myc_KD

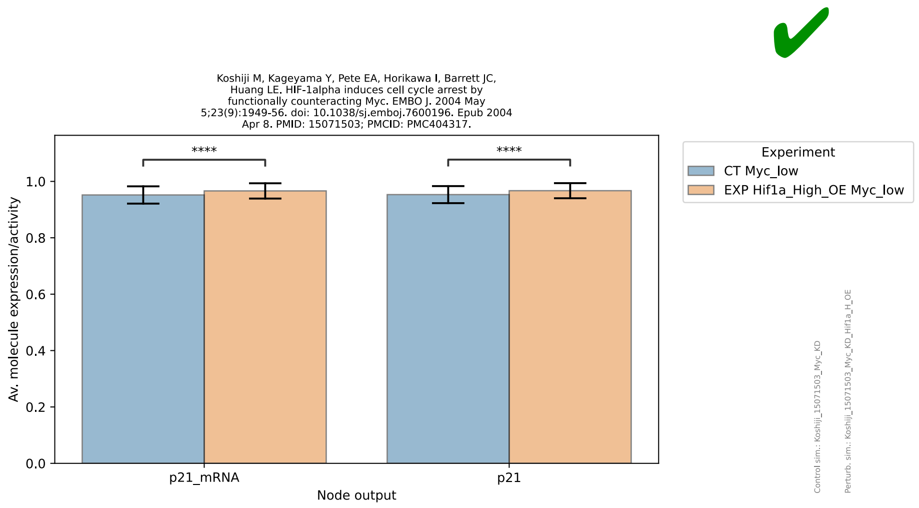
Koshiji M, Kageyama Y, Pete EA, Horikawa I, Barrett JC, Huang LE. HIF-1alpha induces cell cycle arrest by functionally counteracting Myc. EMBO J. 2004 May 5;23(9):1949-56. doi: 10.1038/sj.emboj.7600196. Epub 2004 Apr 8. PMID: 15071503; PMCID: PMC404317.
Av. molecule expression/activity
Node output
p21_mRNA
p21
****
****
Experiment
CT Myc_low
EXP Hif1a_High_OE Myc_low

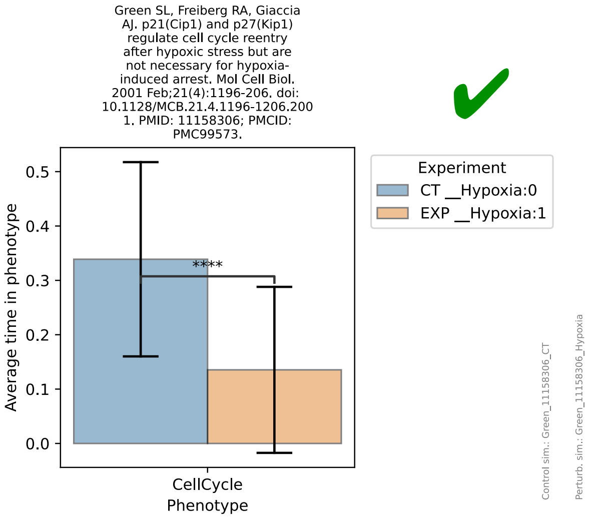
Green SL, Freiberg RA, Giaccia AJ. p21(Cip1) and p27(Kip1) regulate cell cycle reentry after hypoxic stress but are not necessary for hypoxia-induced arrest. Mol Cell Biol. 2001 Feb;21(4):1196-206. doi: 10.1128/MCB.21.4.1196-1206.2001. PMID: 11158306; PMCID: PMC99573.
✔
Experiment
CT __Hypoxia:0
EXP __Hypoxia:1
Average time in phenotype
****
CellCycle
Phenotype
Control sim.: Green_11158306_CT
Perturb. sim.: Green_11158306_Hypoxia

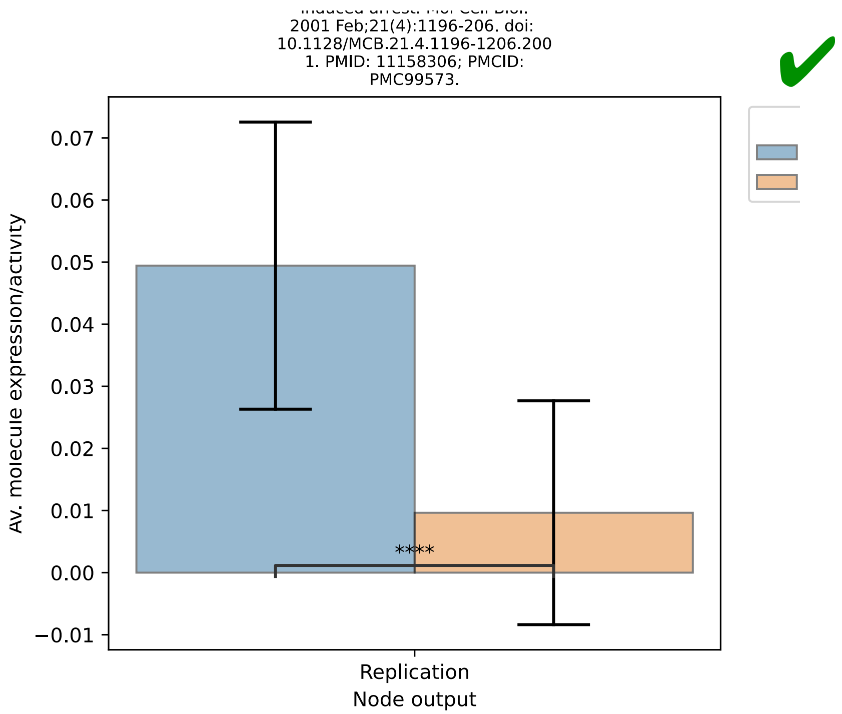
2001 Feb;21(4):1196-206. doi: 10.1128/MCB.21.4.1196-1206.2001. PMID: 11158306; PMCID: PMC99573.
✔
Av. molecule expression/activity
****
Replication
Node output

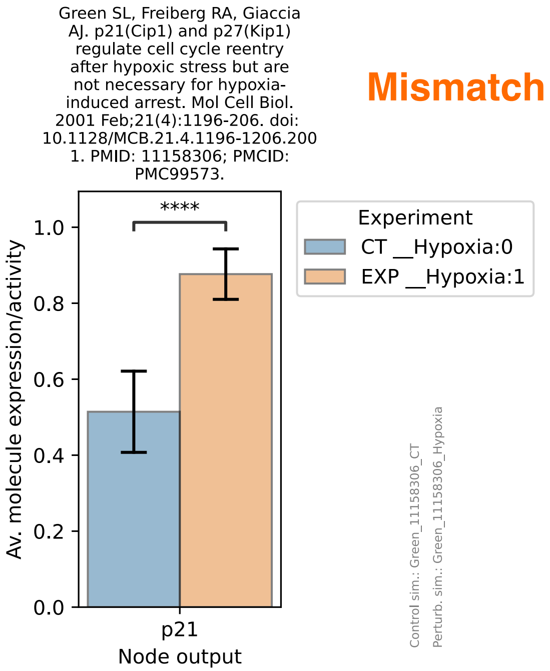
Green SL, Freiberg RA, Giaccia AJ. p21(Cip1) and p27(Kip1) regulate cell cycle reentry after hypoxic stress but are not necessary for hypoxia-induced arrest. Mol Cell Biol. 2001 Feb;21(4):1196-206. doi: 10.1128/MCB.21.4.1196-1206.2001. PMID: 11158306; PMCID: PMC99573.
Mismatch
Experiment
CT __Hypoxia:0
EXP __Hypoxia:1
Av. molecule expression/activity
****
p21
Node output
Control sim.: Green_11158306_CT
Perturb. sim.: Green_11158306_Hypoxia

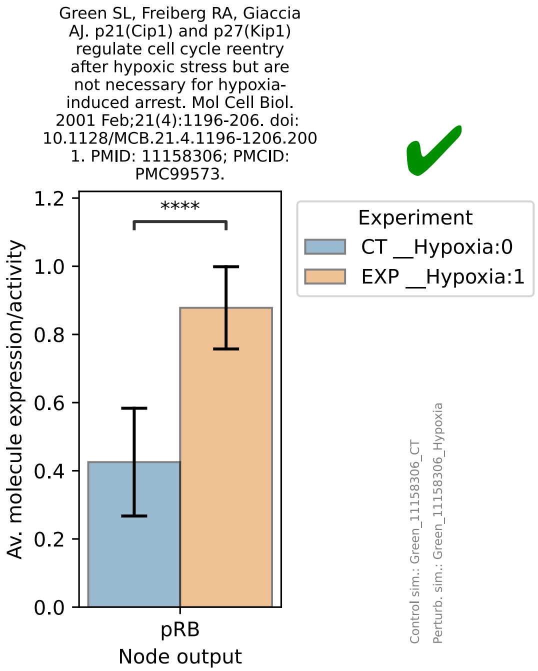
Green SL, Freiberg RA, Giaccia AJ. p21(Cip1) and p27(Kip1) regulate cell cycle reentry after hypoxic stress but are not necessary for hypoxia-induced arrest. Mol Cell Biol. 2001 Feb;21(4):1196-206. doi: 10.1128/MCB.21.4.1196-1206.2001. PMID: 11158306; PMCID: PMC99573.
✔
Experiment
CT __Hypoxia:0
EXP __Hypoxia:1
Av. molecule expression/activity
****
pRB
Node output
Control sim.: Green_11158306_CT
Perturb. sim.: Green_11158306_Hypoxia

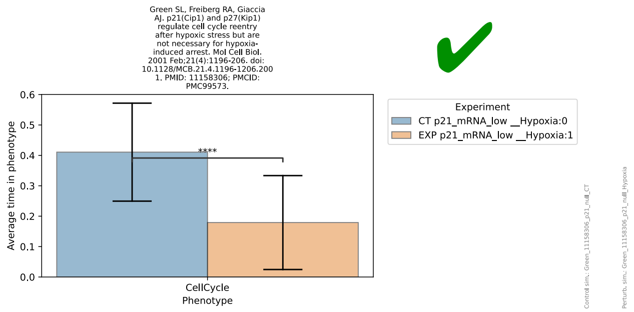
Green SL, Freiberg RA, Giaccia AJ. p21(Cip1) and p27(Kip1) regulate cell cycle reentry after hypoxic stress but are not necessary for hypoxia-induced arrest. Mol Cell Biol. 2001 Feb;21(4):1196-206. doi: 10.1128/MCB.21.4.1196-1206.2001. PMID: 11158306; PMCID: PMC99573.
Average time in phenotype
****
CellCycle
Phenotype
Experiment
CT p21_mRNA_low __Hypoxia:0
EXP p21_mRNA_low __Hypoxia:1
Control sim.: Green_11158306_p21_null_CT
Perturb. sim.: Green_11158306_p21_null_Hypoxia

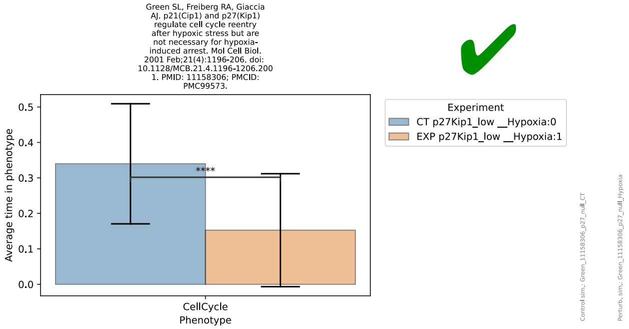
Green SL, Freiberg RA, Giaccia AJ. p21(Cip1) and p27(Kip1) regulate cell cycle reentry after hypoxic stress but are not necessary for hypoxia-induced arrest. Mol Cell Biol. 2001 Feb;21(4):1196-206. doi: 10.1128/MCB.21.4.1196-1206.2001. PMID: 11158306; PMCID: PMC99573.
Average time in phenotype
****
CellCycle
Phenotype
Experiment
CT p27Kip1_low __Hypoxia:0
EXP p27Kip1_low __Hypoxia:1
Control sim.: Green_11158306_p27_null_CT
Perturb. sim.: Green_11158306_p27_null_Hypoxia

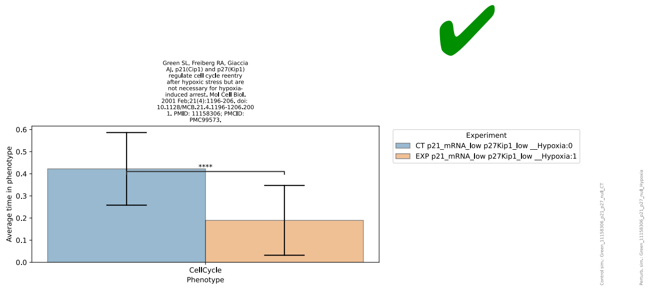
Green SL, Freiberg RA, Giaccia AJ. p21(Cip1) and p27(Kip1) regulate cell cycle reentry after hypoxic stress but are not necessary for hypoxia-induced arrest. Mol Cell Biol. 2001 Feb;21(4):1196-206. doi: 10.1128/MCB.21.4.1196-1206.2001. PMID: 11158306; PMCID: PMC99573.
Average time in phenotype
****
CellCycle
Phenotype
Experiment
CT p21_mRNA_low p27Kip1_low __Hypoxia:0
EXP p21_mRNA_low p27Kip1_low __Hypoxia:1
Control sim.: Green_11158306_p21_p27_null_CT
Perturb. sim.: Green_11158306_p21_p27_null_Hypoxia

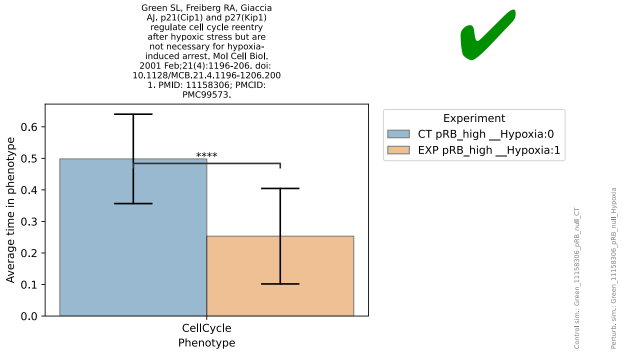
Green SL, Freiberg RA, Giaccia AJ. p21(Cip1) and p27(Kip1) regulate cell cycle reentry after hypoxic stress but are not necessary for hypoxia-induced arrest. Mol Cell Biol. 2001 Feb;21(4):1196-206. doi: 10.1128/MCB.21.4.1196-1206.2001. PMID: 11158306; PMCID: PMC99573.
Average time in phenotype
****
CellCycle
Phenotype
Experiment
CT pRB_high __Hypoxia:0
EXP pRB_high __Hypoxia:1
Control sim.: Green_11158306_pRB_null_CT
Perturb. sim.: Green_11158306_pRB_null_Hypoxia

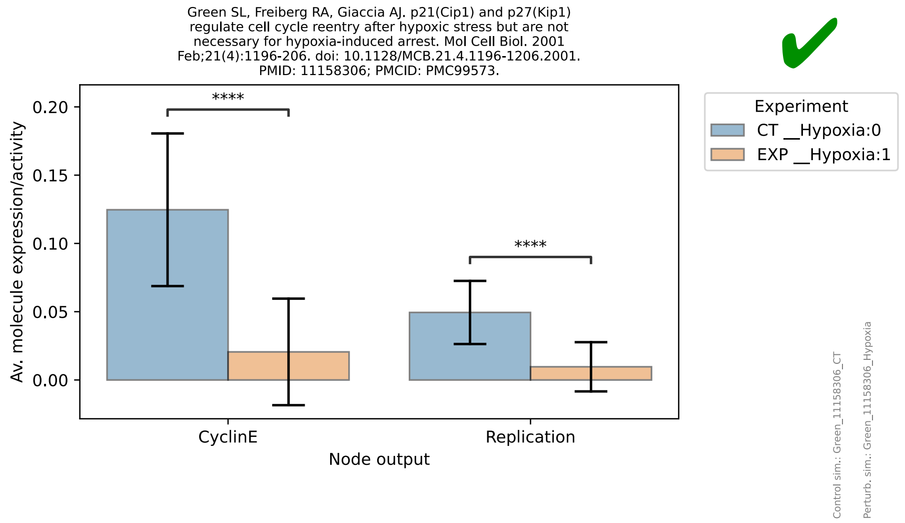

Green SL, Freiberg RA, Giaccia AJ. p21(Cip1) and p27(Kip1) regulate cell cycle reentry after hypoxic stress but are not necessary for hypoxia-induced arrest. Mol Cell Biol. 2001 Feb;21(4):1196-206. doi: 10.1128/MCB.21.4.1196-1206.2001. PMID: 11158306; PMCID: PMC99573.
Av. molecule expression/activity
0.20
0.15
0.10
0.05
0.00
****
****
CyclinE
Replication
Node output
Experiment
CT __Hypoxia:0
EXP __Hypoxia:1
Control sim.: Green_11158306_CT
Perturb. sim.: Green_11158306_Hypoxia


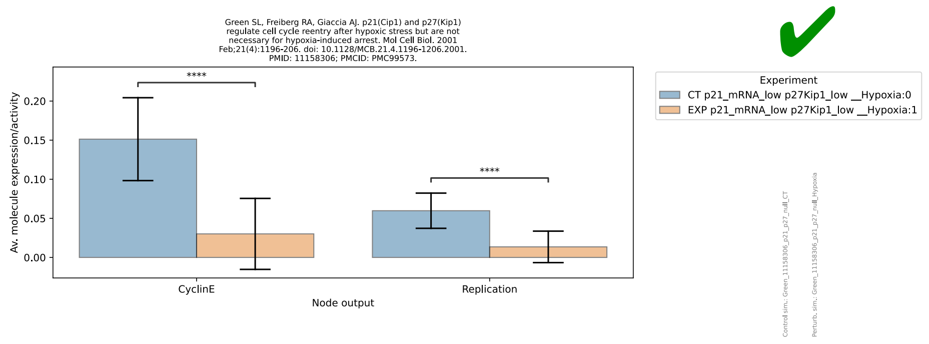

Green SL, Freiberg RA, Giaccia AJ. p21(Cip1) and p27(Kip1) regulate cell cycle reentry after hypoxic stress but are not necessary for hypoxia-induced arrest. Mol Cell Biol. 2001 Feb;21(4):1196-206. doi: 10.1128/MCB.21.4.1196-1206.2001. PMID: 11158306; PMCID: PMC99573.
Av. molecule expression/activity
0.20
0.15
0.10
0.05
0.00
****
****
CyclinE
Replication
Node output
Experiment
CT p21_mRNA_low p27Kip1_low __Hypoxia:0
EXP p21_mRNA_low p27Kip1_low __Hypoxia:1


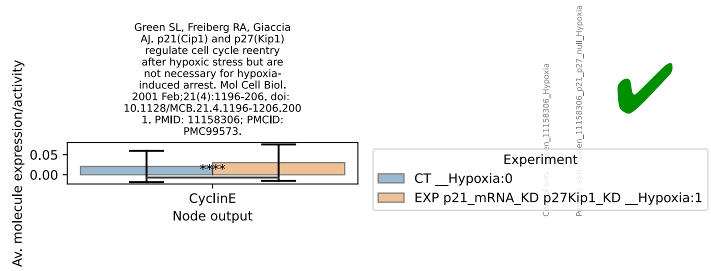

Green SL, Freiberg RA, Giaccia AJ. p21(Cip1) and p27(Kip1) regulate cell cycle reentry after hypoxic stress but are not necessary for hypoxia-induced arrest. Mol Cell Biol. 2001 Feb;21(4):1196-206. doi: 10.1128/MCB.21.4.1196-1206.2001. PMID: 11158306; PMCID: PMC99573.
Av. molecule expression/activity
0.05
0.00
****
CyclinE
Node output
Experiment
CT __Hypoxia:0
EXP p21_mRNA_KD p27Kip1_KD __Hypoxia:1


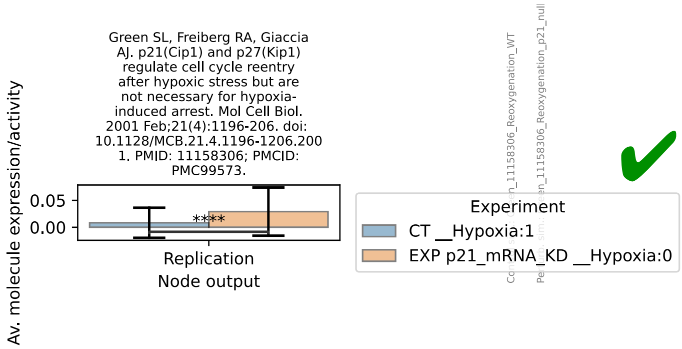

Green SL, Freiberg RA, Giaccia AJ. p21(Cip1) and p27(Kip1) regulate cell cycle reentry after hypoxic stress but are not necessary for hypoxia-induced arrest. Mol Cell Biol. 2001 Feb;21(4):1196-206. doi: 10.1128/MCB.21.4.1196-1206.2001. PMID: 11158306; PMCID: PMC99573.
Av. molecule expression/activity
0.05
0.00
****
Replication
Node output
Experiment
CT __Hypoxia:1
EXP p21_mRNA_KD __Hypoxia:0

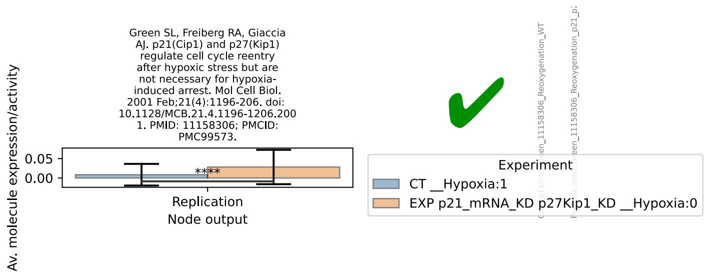
Green SL, Freiberg RA, Giaccia AJ. p21(Cip1) and p27(Kip1) regulate cell cycle reentry after hypoxic stress but are not necessary for hypoxia-induced arrest. Mol Cell Biol. 2001 Feb;21(4):1196-206. doi: 10.1128/MCB.21.4.1196-1206.2001. PMID: 11158306; PMCID: PMC99573.
Av. molecule expression/activity
0.05
0.00
****
Replication
Node output
Experiment
CT __Hypoxia:1
EXP p21_mRNA_KD p27Kip1_KD __Hypoxia:0
Control sim.: Green_11158306_Reoxygenation_WT
Perturb. sim.: Green_11158306_Reoxygenation_p21_p

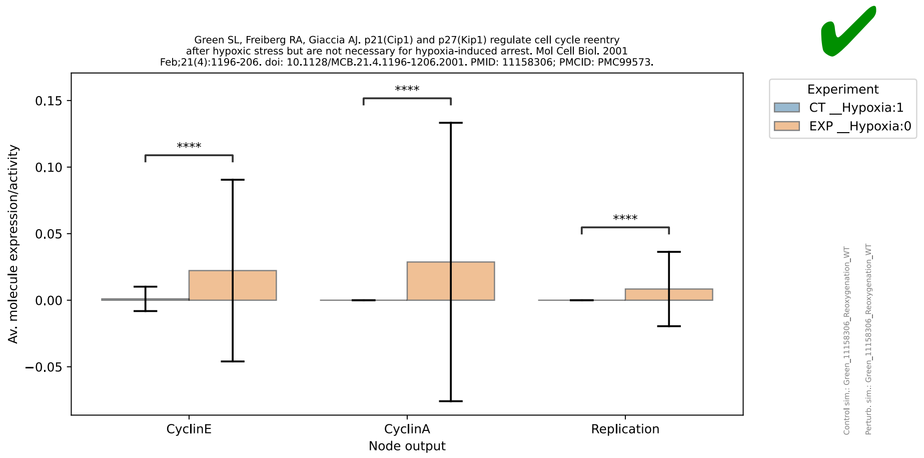
Green SL, Freiberg RA, Giaccia AJ. p21(Cip1) and p27(Kip1) regulate cell cycle reentry after hypoxic stress but are not necessary for hypoxia-induced arrest. Mol Cell Biol. 2001 Feb;21(4):1196-206. doi: 10.1128/MCB.21.4.1196-1206.2001. PMID: 11158306; PMCID: PMC99573.
Av. molecule expression/activity
0.15
0.10
0.05
0.00
−0.05
****
****
****
CyclinE
CyclinA
Replication
Node output
Experiment
CT __Hypoxia:1
EXP __Hypoxia:0
Control sim.: Green_11158306_Reoxygenation_WT
Perturb. sim.: Green_11158306_Reoxygenation_WT

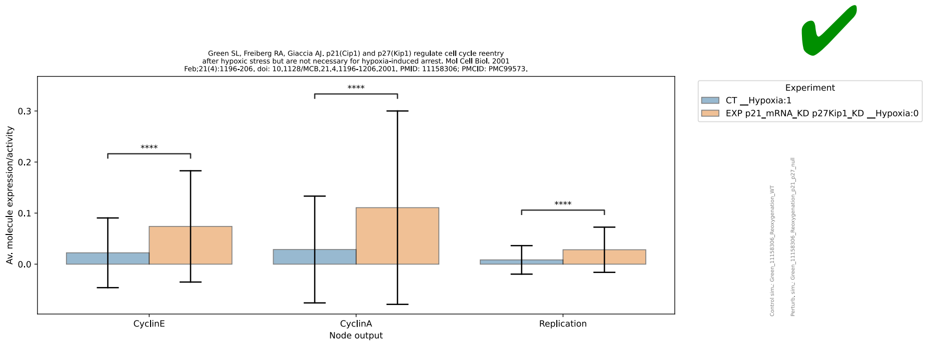
Green SL, Freiberg RA, Giaccia AJ. p21(Cip1) and p27(Kip1) regulate cell cycle reentry after hypoxic stress but are not necessary for hypoxia-induced arrest. Mol Cell Biol. 2001 Feb;21(4):1196-206. doi: 10.1128/MCB.21.4.1196-1206.2001. PMID: 11158306; PMCID: PMC99573.
Av. molecule expression/activity
0.3
0.2
0.1
0.0
****
****
****
CyclinE
CyclinA
Replication
Node output
Experiment
CT __Hypoxia:1
EXP p21_mRNA_KD p27Kip1_KD __Hypoxia:0
Control sim.: Green_11158306_Reoxygenation_WT
Perturb. sim.: Green_11158306_Reoxygenation_p21_p27_null

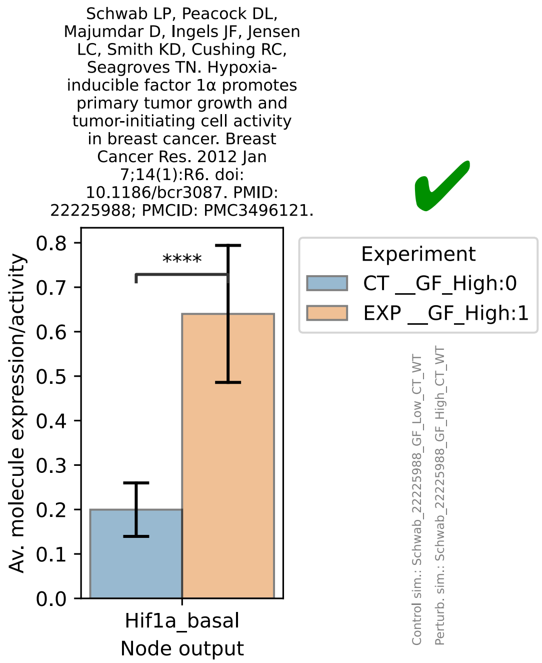
Schwab LP, Peacock DL, Majumdar D, Ingels JF, Jensen LC, Smith KD, Cushing RC, Seagroves TN. Hypoxia-inducible factor 1α promotes primary tumor growth and tumor-initiating cell activity in breast cancer. Breast Cancer Res. 2012 Jan 7;14(1):R6. doi: 10.1186/bcr3087. PMID: 22225988; PMCID: PMC3496121.
Av. molecule expression/activity
0.8
0.7
0.6
0.5
0.4
0.3
0.2
0.1
0.0
****
Hif1a_basal
Node output
Experiment
CT __GF_High:0
EXP __GF_High:1
Control sim.: Schwab_22225988_GF_Low_CT_WT
Perturb. sim.: Schwab_22225988_GF_High_CT_WT

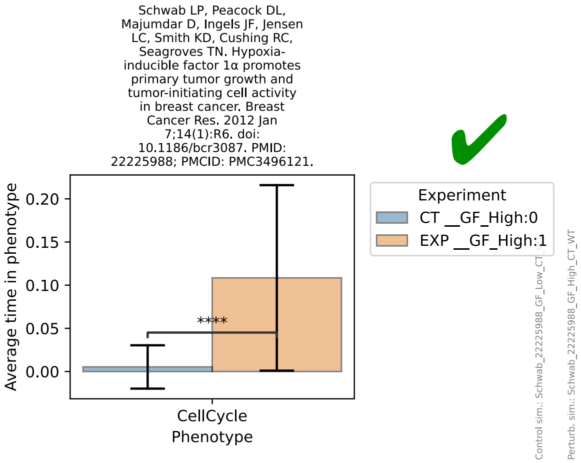

Schwab LP, Peacock DL, Majumdar D, Ingels JF, Jensen LC, Smith KD, Cushing RC, Seagroves TN. Hypoxia-inducible factor 1α promotes primary tumor growth and tumor-initiating cell activity in breast cancer. Breast Cancer Res. 2012 Jan 7;14(1):R6. doi: 10.1186/bcr3087. PMID: 22225988; PMCID: PMC3496121.
✔
Average time in phenotype
0.20
0.15
0.10
0.05
0.00
****
CellCycle
Phenotype
Experiment
CT __GF_High:0
EXP __GF_High:1
Control sim.: Schwab_22225988_GF_Low_CT
Perturb. sim.: Schwab_22225988_GF_High_CT_WT


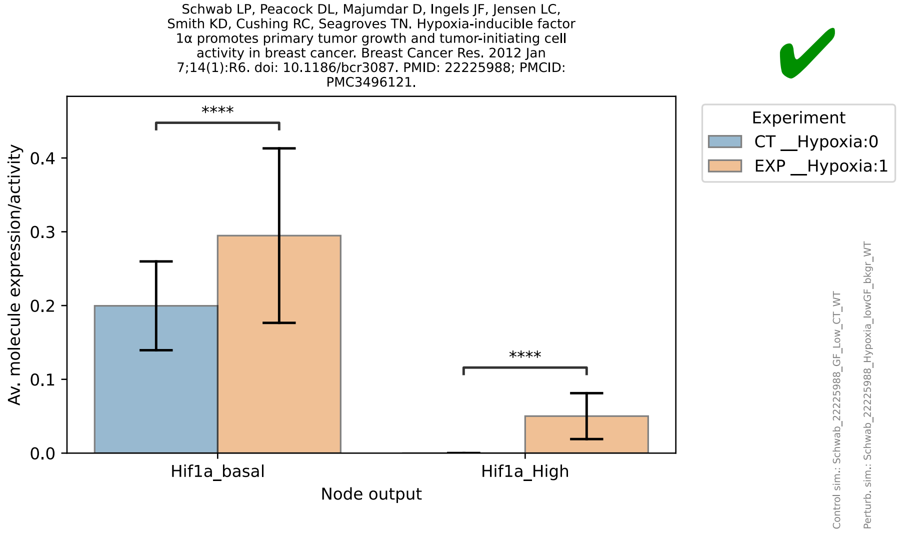

Schwab LP, Peacock DL, Majumdar D, Ingels JF, Jensen LC, Smith KD, Cushing RC, Seagroves TN. Hypoxia-inducible factor 1α promotes primary tumor growth and tumor-initiating cell activity in breast cancer. Breast Cancer Res. 2012 Jan 7;14(1):R6. doi: 10.1186/bcr3087. PMID: 22225988; PMCID: PMC3496121.
✔
Av. molecule expression/activity
0.4
0.3
0.2
0.1
0.0
****
****
Hif1a_basal
Hif1a_High
Node output
Experiment
CT __Hypoxia:0
EXP __Hypoxia:1
Control sim.: Schwab_22225988_GF_Low_CT_WT
Perturb. sim.: Schwab_22225988_Hypoxia_lowGF_bkgr_WT


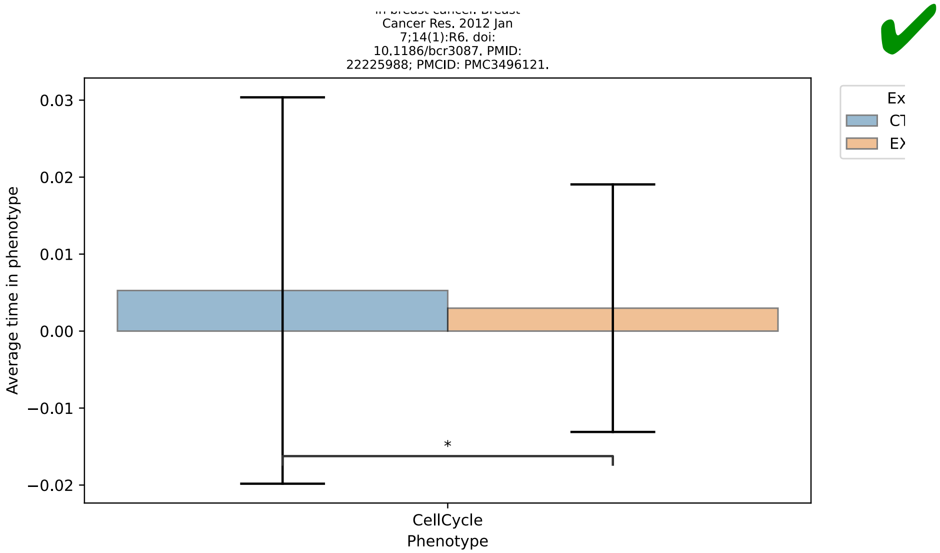

Cancer Res. 2012 Jan 7;14(1):R6. doi: 10.1186/bcr3087. PMID: 22225988; PMCID: PMC3496121.
✔
Average time in phenotype
0.03
0.02
0.01
0.00
−0.01
−0.02
*
CellCycle
Phenotype


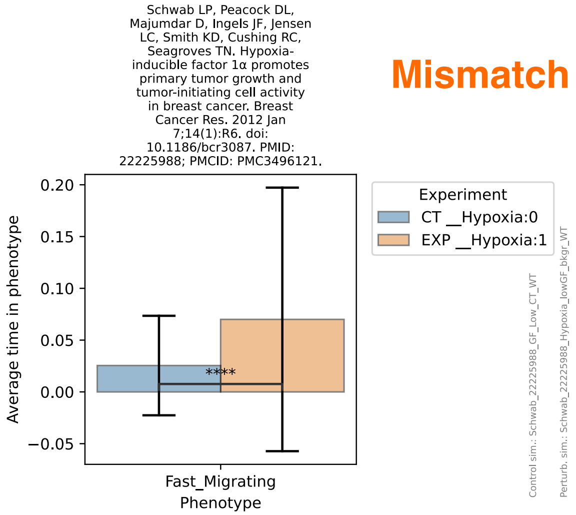

Schwab LP, Peacock DL, Majumdar D, Ingels JF, Jensen LC, Smith KD, Cushing RC, Seagroves TN. Hypoxia-inducible factor 1α promotes primary tumor growth and tumor-initiating cell activity in breast cancer. Breast Cancer Res. 2012 Jan 7;14(1):R6. doi: 10.1186/bcr3087. PMID: 22225988; PMCID: PMC3496121.
Mismatch
Average time in phenotype
0.20
0.15
0.10
0.05
0.00
−0.05
****
Fast_Migrating
Phenotype
Experiment
CT __Hypoxia:0
EXP __Hypoxia:1
Control sim.: Schwab_22225988_GF_Low_CT_WT
Perturb. sim.: Schwab_22225988_Hypoxia_lowGF_bkgr_WT

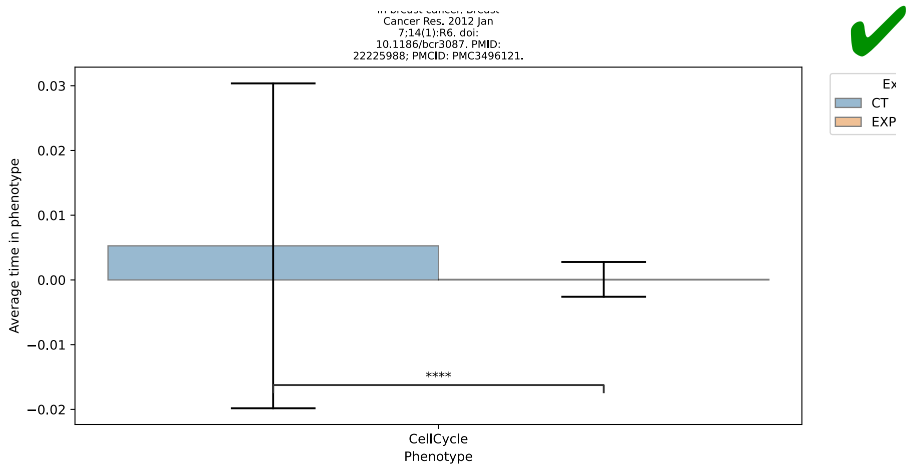
Cancer Res. 2012 Jan
7;14(1):R6. doi:
10.1186/bcr3087. PMID:
22225988; PMCID: PMC3496121.
Average time in phenotype
0.03
0.02
0.01
0.00
−0.01
−0.02
****
CellCycle
Phenotype
CT
EXP

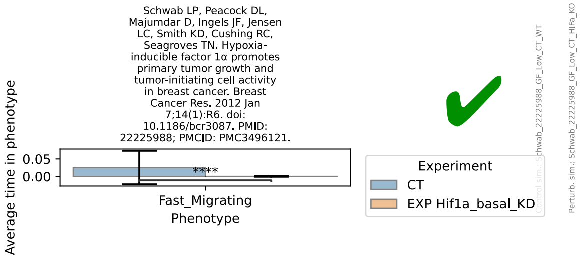
Schwab LP, Peacock DL,
Majumdar D, Ingels JF, Jensen
LC, Smith KD, Cushing RC,
Seagroves TN. Hypoxia-
inducible factor 1α promotes
primary tumor growth and
tumor-initiating cell activity
in breast cancer. Breast
Cancer Res. 2012 Jan
7;14(1):R6. doi:
10.1186/bcr3087. PMID:
22225988; PMCID: PMC3496121.
Average time in phenotype
0.05
0.00
****
Fast_Migrating
Phenotype
Experiment
CT
EXP Hif1a_basal_KD

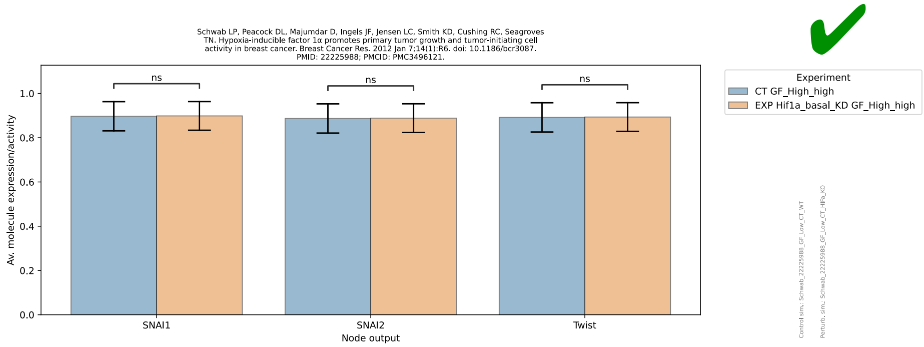
Schwab LP, Peacock DL, Majumdar D, Ingels JF, Jensen LC, Smith KD, Cushing RC, Seagroves
TN. Hypoxia-inducible factor 1α promotes primary tumor growth and tumor-initiating cell
activity in breast cancer. Breast Cancer Res. 2012 Jan 7;14(1):R6. doi: 10.1186/bcr3087.
PMID: 22225988; PMCID: PMC3496121.
Av. molecule expression/activity
1.0
0.8
0.6
0.4
0.2
0.0
ns
ns
ns
SNAI1
SNAI2
Twist
Node output
Experiment
CT GF_High_high
EXP Hif1a_basal_KD GF_High_high

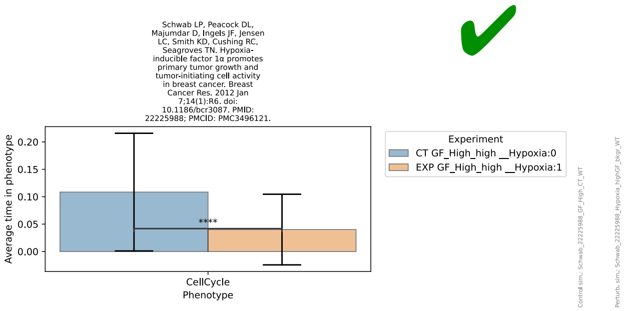
Schwab LP, Peacock DL,
Majumdar D, Ingels JF, Jensen
LC, Smith KD, Cushing RC,
Seagroves TN. Hypoxia-
inducible factor 1α promotes
primary tumor growth and
tumor-initiating cell activity
in breast cancer. Breast
Cancer Res. 2012 Jan
7;14(1):R6. doi:
10.1186/bcr3087. PMID:
22225988; PMCID: PMC3496121.
Average time in phenotype
0.20
0.15
0.10
0.05
0.00
****
CellCycle
Phenotype
Experiment
CT GF_High_high __Hypoxia:0
EXP GF_High_high __Hypoxia:1

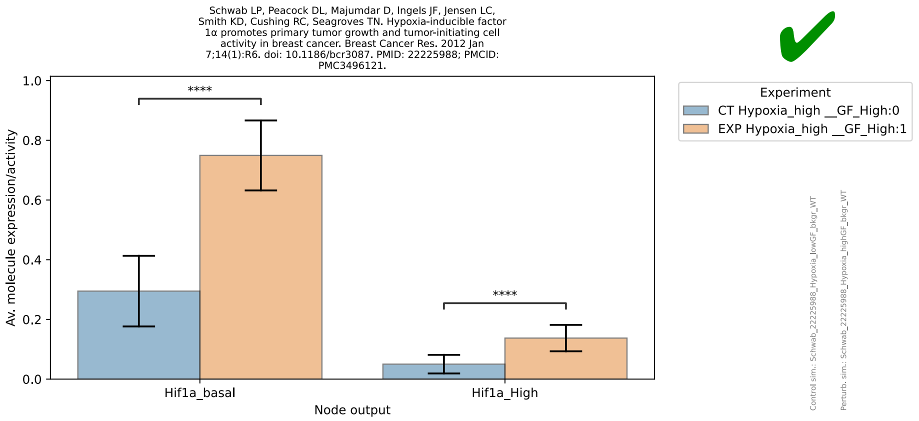
Schwab LP, Peacock DL, Majumdar D, Ingels JF, Jensen LC, Smith KD, Cushing RC, Seagroves TN. Hypoxia-inducible factor 1α promotes primary tumor growth and tumor-initiating cell activity in breast cancer. Breast Cancer Res. 2012 Jan 7;14(1):R6. doi: 10.1186/bcr3087. PMID: 22225988; PMCID: PMC3496121.
Av. molecule expression/activity
1.0
0.8
0.6
0.4
0.2
0.0
****
****
Hif1a_basal
Hif1a_High
Node output
Experiment
CT Hypoxia_high __GF_High:0
EXP Hypoxia_high __GF_High:1
Control sim.: Schwab_22225988_Hypoxia_lowGF_bkgr_WT
Perturb. sim.: Schwab_22225988_Hypoxia_highGF_bkgr_WT

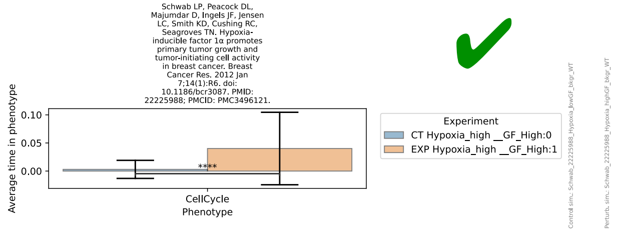
Schwab LP, Peacock DL, Majumdar D, Ingels JF, Jensen LC, Smith KD, Cushing RC, Seagroves TN. Hypoxia-inducible factor 1α promotes primary tumor growth and tumor-initiating cell activity in breast cancer. Breast Cancer Res. 2012 Jan 7;14(1):R6. doi: 10.1186/bcr3087. PMID: 22225988; PMCID: PMC3496121.
Average time in phenotype
0.10
0.05
0.00
****
CellCycle
Phenotype
Experiment
CT Hypoxia_high __GF_High:0
EXP Hypoxia_high __GF_High:1

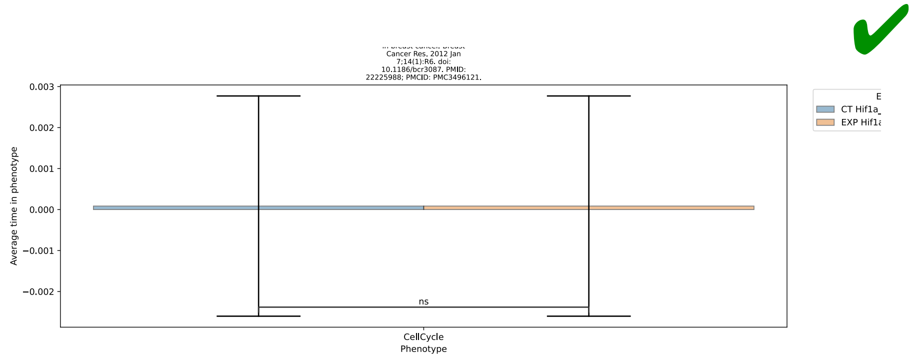
Cancer Res. 2012 Jan 7;14(1):R6. doi: 10.1186/bcr3087. PMID: 22225988; PMCID: PMC3496121.
Average time in phenotype
0.003
0.002
0.001
0.000
−0.001
−0.002
ns
CellCycle
Phenotype
CT Hif1a_
EXP Hif1a

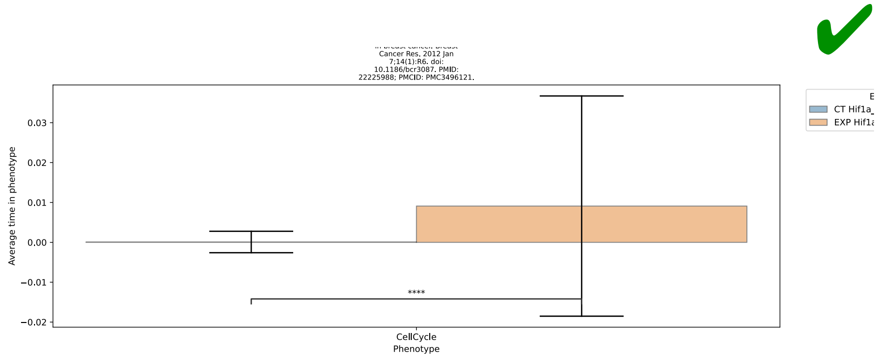
Cancer Res. 2012 Jan 7;14(1):R6. doi: 10.1186/bcr3087. PMID: 22225988; PMCID: PMC3496121.
Average time in phenotype
0.03
0.02
0.01
0.00
−0.01
−0.02
****
CellCycle
Phenotype
CT Hif1a_
EXP Hif1a

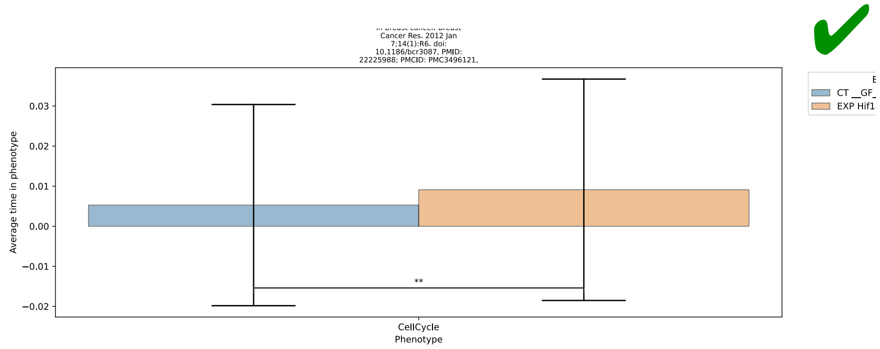
Cancer Res. 2012 Jan
7;14(1):R6. doi:
10.1186/bcr3087. PMID:
22225988; PMCID: PMC3496121.
Average time in phenotype
**
CellCycle
Phenotype
CT __GF_
EXP Hif1

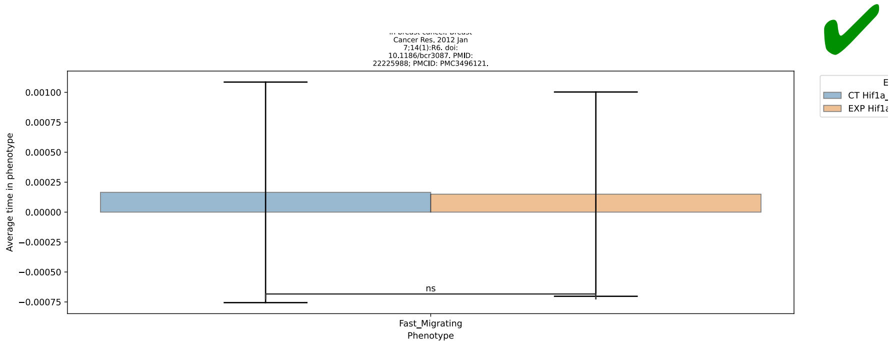
Cancer Res. 2012 Jan
7;14(1):R6. doi:
10.1186/bcr3087. PMID:
22225988; PMCID: PMC3496121.
Average time in phenotype
ns
Fast_Migrating
Phenotype
CT Hif1a_
EXP Hif1a

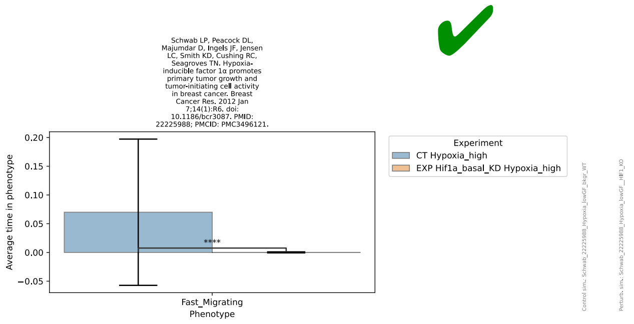
Schwab LP, Peacock DL,
Majumdar D, Ingels JF, Jensen
LC, Smith KD, Cushing RC,
Seagroves TN. Hypoxia-
inducible factor 1α promotes
primary tumor growth and
tumor-initiating cell activity
in breast cancer. Breast
Cancer Res. 2012 Jan
7;14(1):R6. doi:
10.1186/bcr3087. PMID:
22225988; PMCID: PMC3496121.
Average time in phenotype
****
Fast_Migrating
Phenotype
Experiment
CT Hypoxia_high
EXP Hif1a_basal_KD Hypoxia_high

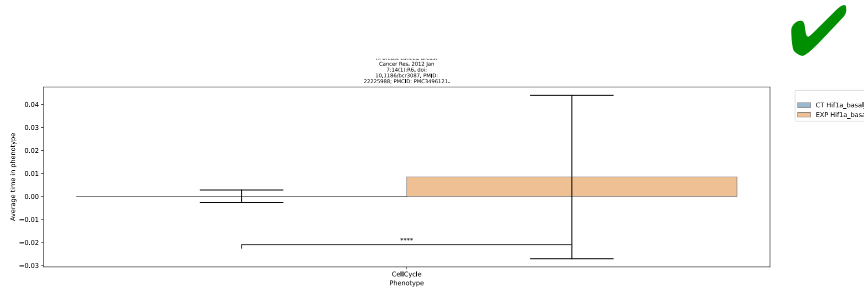
Cancer Res. 2012 Jan
7;14(1):R6. doi:
10.1186/bcr3087. PMID:
22225988; PMCID: PMC3496121.
Average time in phenotype
****
CellCycle
Phenotype
CT Hif1a_basal
EXP Hif1a_basa

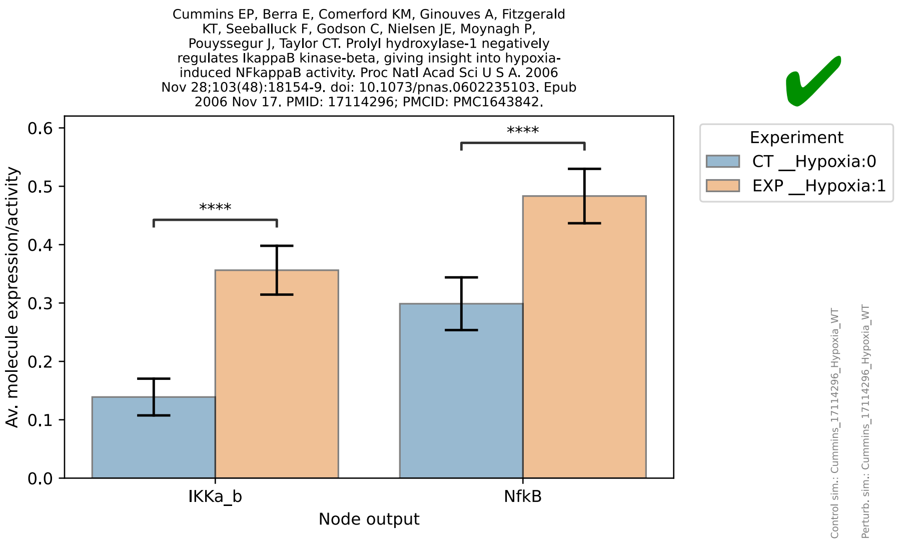
Cummins EP, Berra E, Comerford KM, Ginouves A, Fitzgerald KT, Seeballuck F, Godson C, Nielsen JE, Moynagh P, Pouyssegur J, Taylor CT. Prolyl hydroxylase-1 negatively regulates IkappaB kinase-beta, giving insight into hypoxia-induced NFkappaB activity. Proc Natl Acad Sci U S A. 2006 Nov 28;103(48):18154-9. doi: 10.1073/pnas.0602235103. Epub 2006 Nov 17. PMID: 17114296; PMCID: PMC1643842.
Av. molecule expression/activity
Node output
IKKa_b
NfkB
****
Experiment
CT __Hypoxia:0
EXP __Hypoxia:1
Control sim.: Cummins_17114296_Hypoxia_WT
Perturb. sim.: Cummins_17114296_Hypoxia_WT

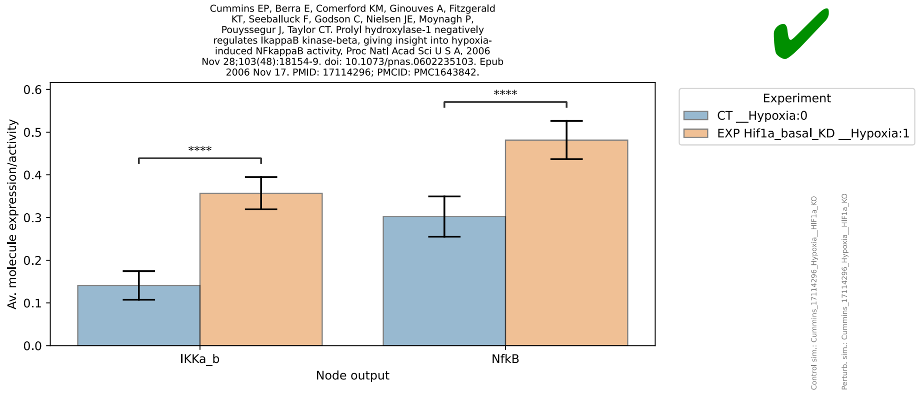
Cummins EP, Berra E, Comerford KM, Ginouves A, Fitzgerald KT, Seeballuck F, Godson C, Nielsen JE, Moynagh P, Pouyssegur J, Taylor CT. Prolyl hydroxylase-1 negatively regulates IkappaB kinase-beta, giving insight into hypoxia-induced NFkappaB activity. Proc Natl Acad Sci U S A. 2006 Nov 28;103(48):18154-9. doi: 10.1073/pnas.0602235103. Epub 2006 Nov 17. PMID: 17114296; PMCID: PMC1643842.
Av. molecule expression/activity
Node output
IKKa_b
NfkB
****
Experiment
CT __Hypoxia:0
EXP Hif1a_basal_KD __Hypoxia:1

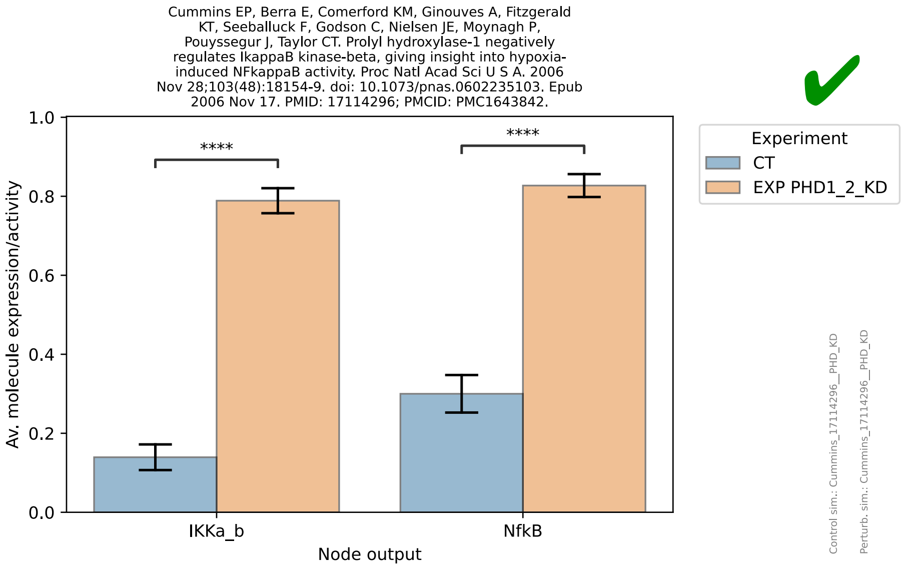
Cummins EP, Berra E, Comerford KM, Ginouves A, Fitzgerald KT, Seeballuck F, Godson C, Nielsen JE, Moynagh P, Pouyssegur J, Taylor CT. Prolyl hydroxylase-1 negatively regulates IkappaB kinase-beta, giving insight into hypoxia-induced NFkappaB activity. Proc Natl Acad Sci U S A. 2006 Nov 28;103(48):18154-9. doi: 10.1073/pnas.0602235103. Epub 2006 Nov 17. PMID: 17114296; PMCID: PMC1643842.
Av. molecule expression/activity
Node output
IKKa_b
NfkB
****
Experiment
CT
EXP PHD1_2_KD
Control sim.: Cummins_17114296__PHD_KD
Perturb. sim.: Cummins_17114296__PHD_KD

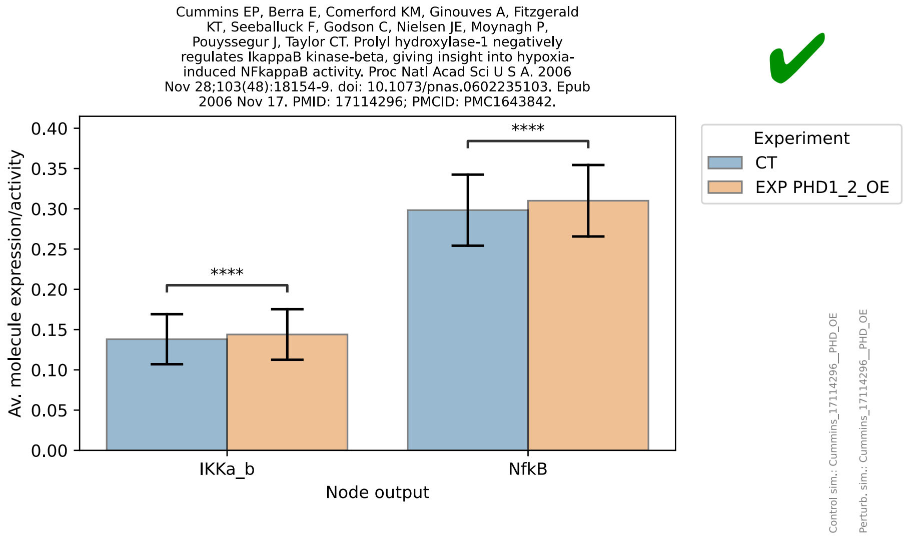
Cummins EP, Berra E, Comerford KM, Ginouves A, Fitzgerald KT, Seeballuck F, Godson C, Nielsen JE, Moynagh P, Pouyssegur J, Taylor CT. Prolyl hydroxylase-1 negatively regulates IkappaB kinase-beta, giving insight into hypoxia-induced NFkappaB activity. Proc Natl Acad Sci U S A. 2006 Nov 28;103(48):18154-9. doi: 10.1073/pnas.0602235103. Epub 2006 Nov 17. PMID: 17114296; PMCID: PMC1643842.
Av. molecule expression/activity
Node output
IKKa_b
NfkB
****
Experiment
CT
EXP PHD1_2_OE
Control sim.: Cummins_17114296__PHD_OE
Perturb. sim.: Cummins_17114296__PHD_OE

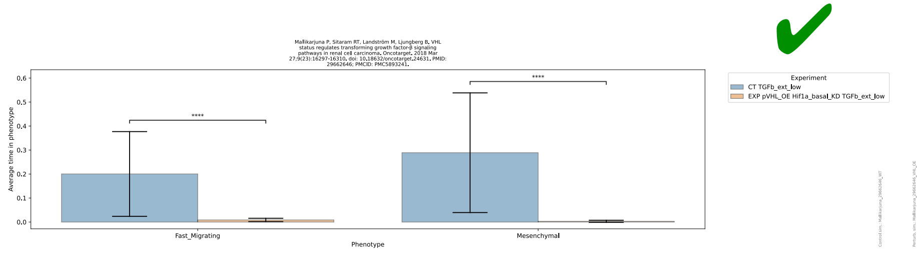

Mallikarjuna P, Sitaram RT, Landström M, Ljungberg B. VHL status regulates transforming growth factor-β signaling pathways in renal cell carcinoma. Oncotarget. 2018 Mar 27;9(23):16297-16310. doi: 10.18632/oncotarget.24631. PMID: 29662646; PMCID: PMC5893241.
Average time in phenotype
Phenotype
Fast_Migrating
Mesenchymal
****
Experiment
CT TGFb_ext_low
EXP pVHL_OE Hif1a_basal_KD TGFb_ext_low
Control sim.: Mallikarjuna_29662646_WT
Perturb. sim.: Mallikarjuna_29662646_VHL_OE


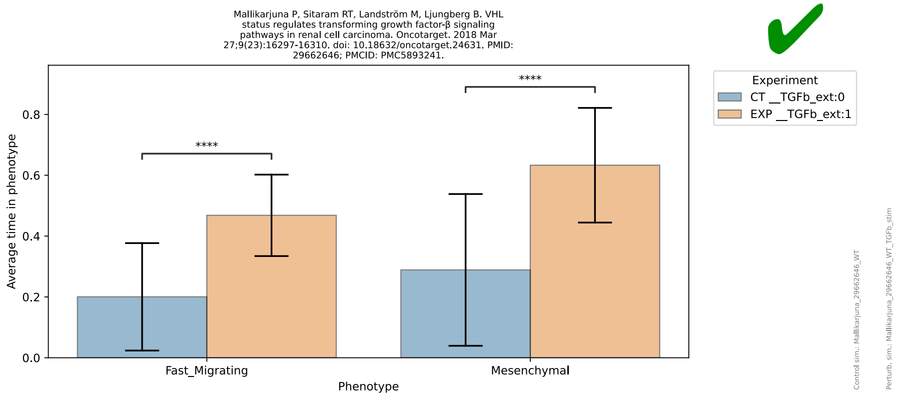

Mallikarjuna P, Sitaram RT, Landström M, Ljungberg B. VHL status regulates transforming growth factor-β signaling pathways in renal cell carcinoma. Oncotarget. 2018 Mar 27;9(23):16297-16310. doi: 10.18632/oncotarget.24631. PMID: 29662646; PMCID: PMC5893241.
Average time in phenotype
Phenotype
Fast_Migrating
Mesenchymal
****
Experiment
CT __TGFb_ext:0
EXP __TGFb_ext:1
Control sim.: Mallikarjuna_29662646_WT
Perturb. sim.: Mallikarjuna_29662646_WT_TGFb_stim


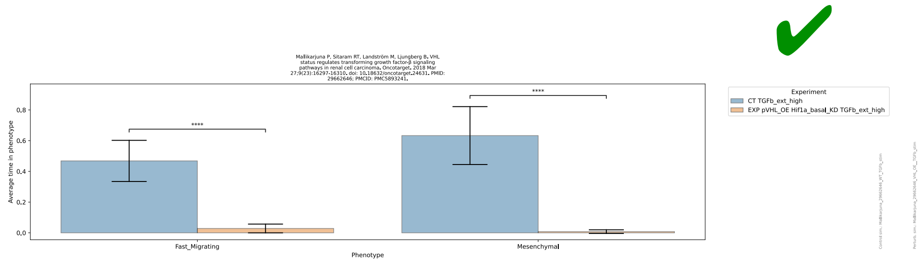

Mallikarjuna P, Sitaram RT, Landström M, Ljungberg B. VHL status regulates transforming growth factor-β signaling pathways in renal cell carcinoma. Oncotarget. 2018 Mar 27;9(23):16297-16310. doi: 10.18632/oncotarget.24631. PMID: 29662646; PMCID: PMC5893241.
Average time in phenotype
Phenotype
Fast_Migrating
Mesenchymal
****
Experiment
CT TGFb_ext_high
EXP pVHL_OE Hif1a_basal_KD TGFb_ext_high
Control sim.: Mallikarjuna_29662646_WT_TGFb_stim
Perturb. sim.: Mallikarjuna_29662646_VHL_OE__TGFb_stim


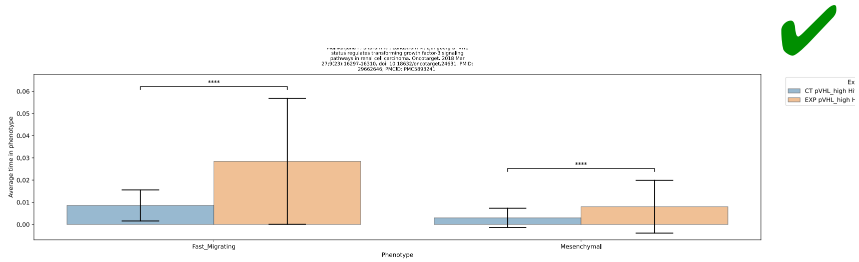

status regulates transforming growth factor-β signaling pathways in renal cell carcinoma. Oncotarget. 2018 Mar 27;9(23):16297-16310. doi: 10.18632/oncotarget.24631. PMID: 29662646; PMCID: PMC5893241.
Average time in phenotype
Phenotype
Fast_Migrating
Mesenchymal
****

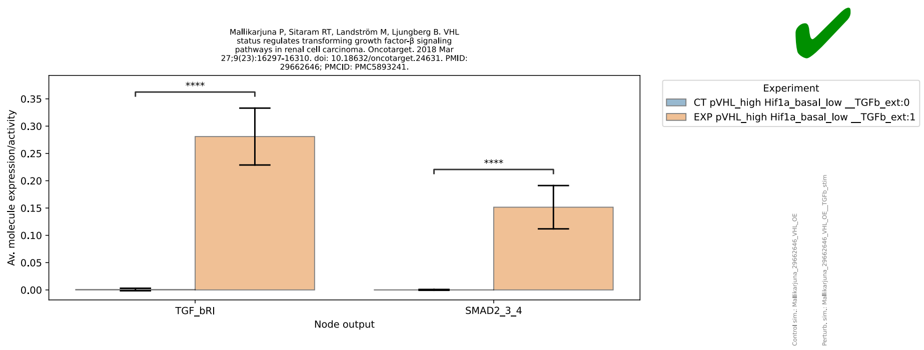
Mallikarjuna P, Sitaram RT, Landström M, Ljungberg B. VHL status regulates transforming growth factor-β signaling pathways in renal cell carcinoma. Oncotarget. 2018 Mar 27;9(23):16297-16310. doi: 10.18632/oncotarget.24631. PMID: 29662646; PMCID: PMC5893241.
Av. molecule expression/activity
0.35
0.30
0.25
0.20
0.15
0.10
0.05
0.00
****
****
TGF_bRI
SMAD2_3_4
Node output
Experiment
CT pVHL_high Hif1a_basal_low __TGFb_ext:0
EXP pVHL_high Hif1a_basal_low __TGFb_ext:1
Control sim.: Mallikarjuna_29662646_VHL_OE
Perturb. sim.: Mallikarjuna_29662646_VHL_OE__TGFb_stim

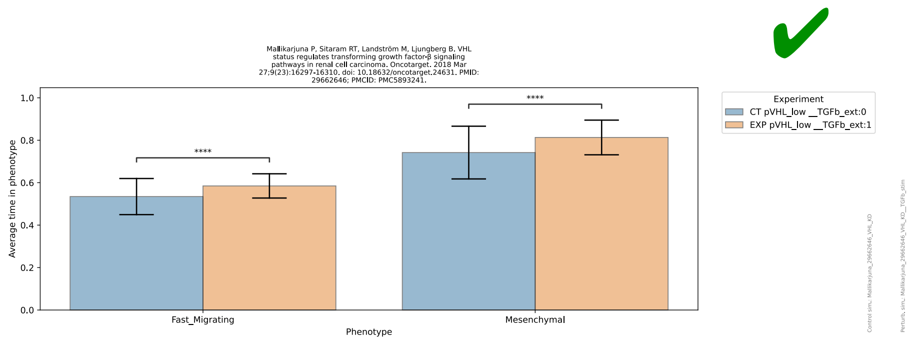
Mallikarjuna P, Sitaram RT, Landström M, Ljungberg B. VHL status regulates transforming growth factor-β signaling pathways in renal cell carcinoma. Oncotarget. 2018 Mar 27;9(23):16297-16310. doi: 10.18632/oncotarget.24631. PMID: 29662646; PMCID: PMC5893241.
Average time in phenotype
1.0
0.8
0.6
0.4
0.2
0.0
****
****
Fast_Migrating
Mesenchymal
Phenotype
Experiment
CT pVHL_low __TGFb_ext:0
EXP pVHL_low __TGFb_ext:1
Control sim.: Mallikarjuna_29662646_VHL_KD
Perturb. sim.: Mallikarjuna_29662646_VHL_KD__TGFb_stim

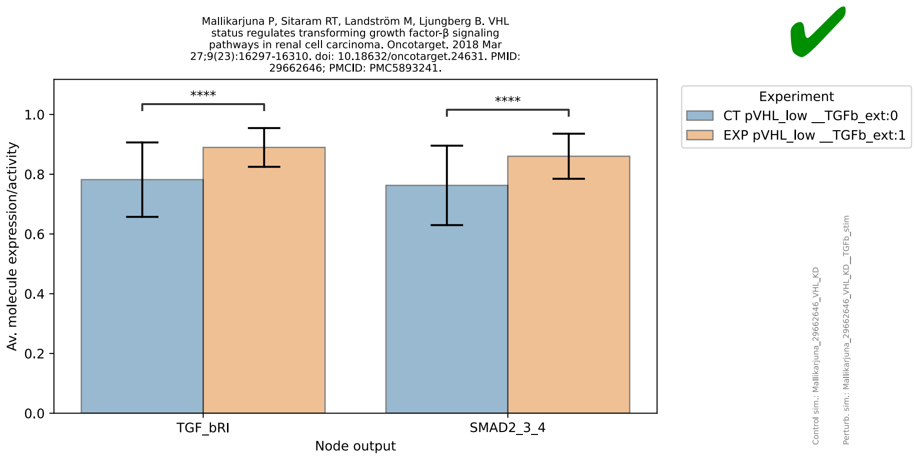
Mallikarjuna P, Sitaram RT, Landström M, Ljungberg B. VHL status regulates transforming growth factor-β signaling pathways in renal cell carcinoma. Oncotarget. 2018 Mar 27;9(23):16297-16310. doi: 10.18632/oncotarget.24631. PMID: 29662646; PMCID: PMC5893241.
Av. molecule expression/activity
1.0
0.8
0.6
0.4
0.2
0.0
****
****
TGF_bRI
SMAD2_3_4
Node output
Experiment
CT pVHL_low __TGFb_ext:0
EXP pVHL_low __TGFb_ext:1
Control sim.: Mallikarjuna_29662646_VHL_KD
Perturb. sim.: Mallikarjuna_29662646_VHL_KD__TGFb_stim

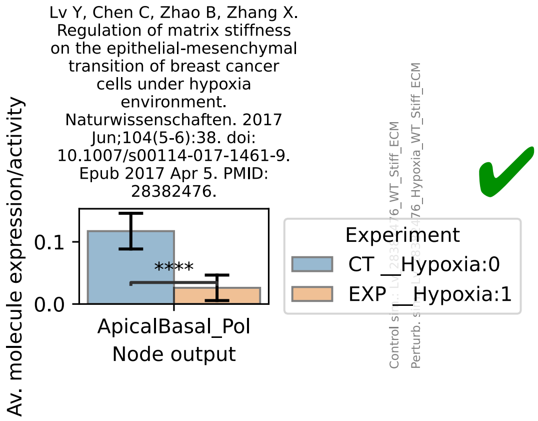
Lv Y, Chen C, Zhao B, Zhang X. Regulation of matrix stiffness on the epithelial-mesenchymal transition of breast cancer cells under hypoxia environment. Naturwissenschaften. 2017 Jun;104(5-6):38. doi: 10.1007/s00114-017-1461-9. Epub 2017 Apr 5. PMID: 28382476.
Av. molecule expression/activity
0.1
0.0
****
ApicalBasal_Pol
Node output
Experiment
CT __Hypoxia:0
EXP __Hypoxia:1
Control sim.: Lv_28382476_WT_Stiff_ECM
Perturb. sim.: Lv_28382476_Hypoxia_WT_Stiff_ECM

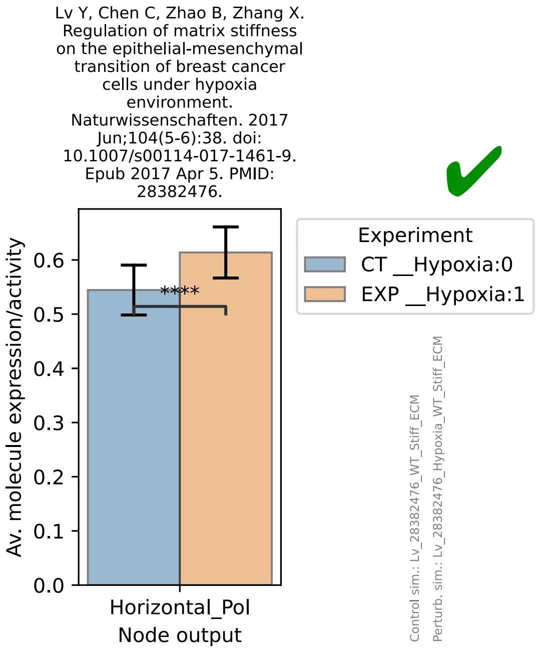
Lv Y, Chen C, Zhao B, Zhang X. Regulation of matrix stiffness on the epithelial-mesenchymal transition of breast cancer cells under hypoxia environment. Naturwissenschaften. 2017 Jun;104(5-6):38. doi: 10.1007/s00114-017-1461-9. Epub 2017 Apr 5. PMID: 28382476.
Av. molecule expression/activity
0.0
0.1
0.2
0.3
0.4
0.5
0.6
****
Horizontal_Pol
Node output
Experiment
CT __Hypoxia:0
EXP __Hypoxia:1
Control sim.: Lv_28382476_WT_Stiff_ECM
Perturb. sim.: Lv_28382476_Hypoxia_WT_Stiff_ECM

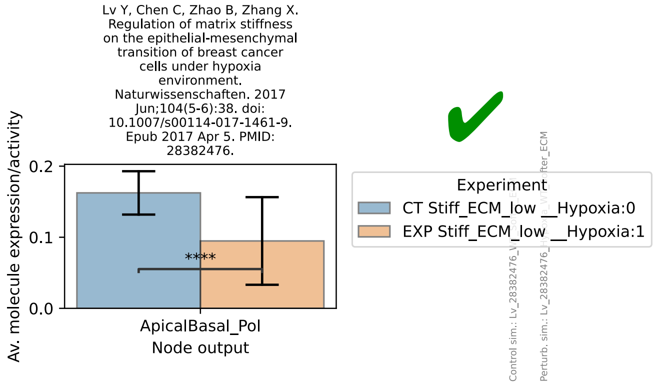
Lv Y, Chen C, Zhao B, Zhang X. Regulation of matrix stiffness on the epithelial-mesenchymal transition of breast cancer cells under hypoxia environment. Naturwissenschaften. 2017 Jun;104(5-6):38. doi: 10.1007/s00114-017-1461-9. Epub 2017 Apr 5. PMID: 28382476.
Av. molecule expression/activity
0.0
0.1
0.2
****
ApicalBasal_Pol
Node output
Experiment
CT Stiff_ECM_low __Hypoxia:0
EXP Stiff_ECM_low __Hypoxia:1

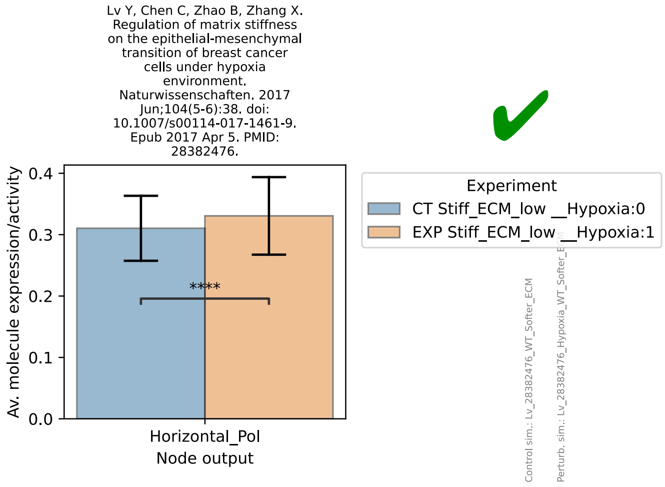
Lv Y, Chen C, Zhao B, Zhang X. Regulation of matrix stiffness on the epithelial-mesenchymal transition of breast cancer cells under hypoxia environment. Naturwissenschaften. 2017 Jun;104(5-6):38. doi: 10.1007/s00114-017-1461-9. Epub 2017 Apr 5. PMID: 28382476.
Av. molecule expression/activity
0.0
0.1
0.2
0.3
0.4
****
Horizontal_Pol
Node output
Experiment
CT Stiff_ECM_low __Hypoxia:0
EXP Stiff_ECM_low __Hypoxia:1
Control sim.: Lv_28382476_WT_Softer_ECM
Perturb. sim.: Lv_28382476_Hypoxia_WT_Softer_ECM

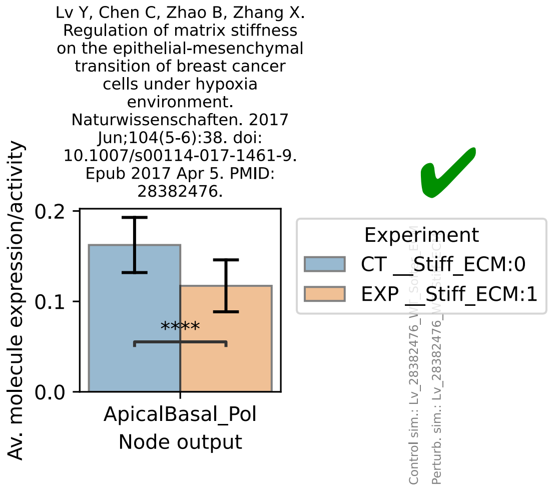
Lv Y, Chen C, Zhao B, Zhang X. Regulation of matrix stiffness on the epithelial-mesenchymal transition of breast cancer cells under hypoxia environment. Naturwissenschaften. 2017 Jun;104(5-6):38. doi: 10.1007/s00114-017-1461-9. Epub 2017 Apr 5. PMID: 28382476.
Av. molecule expression/activity
0.0
0.1
0.2
****
ApicalBasal_Pol
Node output
Experiment
CT __Stiff_ECM:0
EXP __Stiff_ECM:1

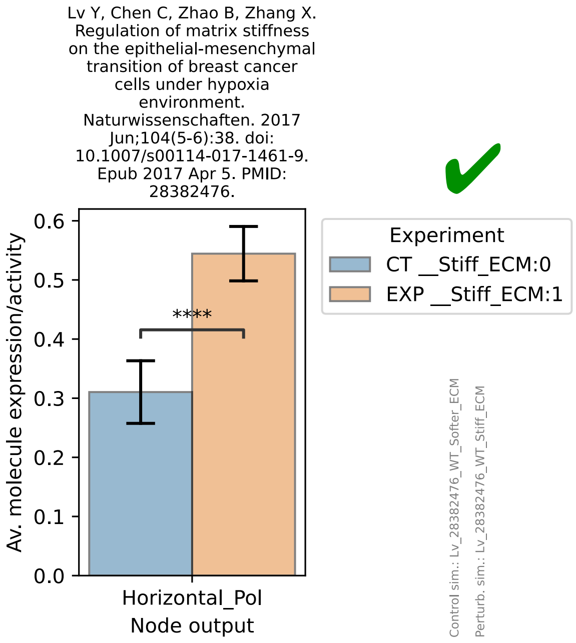
Lv Y, Chen C, Zhao B, Zhang X. Regulation of matrix stiffness on the epithelial-mesenchymal transition of breast cancer cells under hypoxia environment. Naturwissenschaften. 2017 Jun;104(5-6):38. doi: 10.1007/s00114-017-1461-9. Epub 2017 Apr 5. PMID: 28382476.
✔
Av. molecule expression/activity
0.6
0.5
0.4
0.3
0.2
0.1
0.0
****
Horizontal_Pol
Node output
Experiment
CT __Stiff_ECM:0
EXP __Stiff_ECM:1
Control sim.: Lv_28382476_WT_Softer_ECM
Perturb. sim.: Lv_28382476_WT_Stiff_ECM

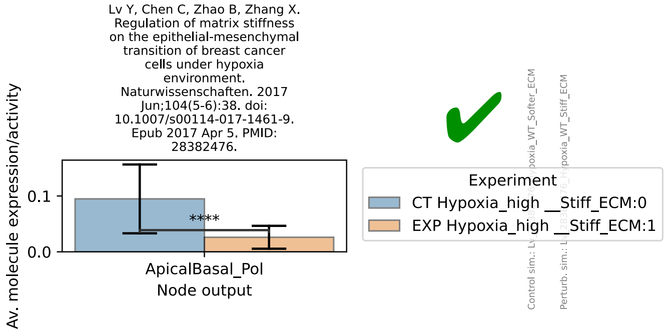
Lv Y, Chen C, Zhao B, Zhang X. Regulation of matrix stiffness on the epithelial-mesenchymal transition of breast cancer cells under hypoxia environment. Naturwissenschaften. 2017 Jun;104(5-6):38. doi: 10.1007/s00114-017-1461-9. Epub 2017 Apr 5. PMID: 28382476.
✔
Av. molecule expression/activity
0.1
0.0
****
ApicalBasal_Pol
Node output
Experiment
CT Hypoxia_high __Stiff_ECM:0
EXP Hypoxia_high __Stiff_ECM:1

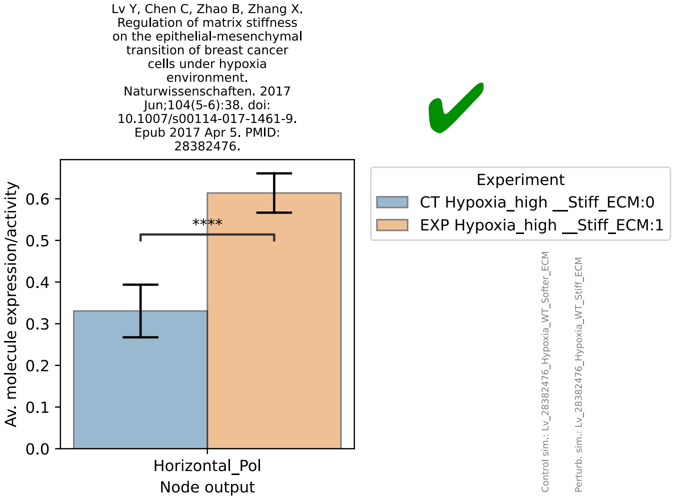
Lv Y, Chen C, Zhao B, Zhang X. Regulation of matrix stiffness on the epithelial-mesenchymal transition of breast cancer cells under hypoxia environment. Naturwissenschaften. 2017 Jun;104(5-6):38. doi: 10.1007/s00114-017-1461-9. Epub 2017 Apr 5. PMID: 28382476.
✔
Av. molecule expression/activity
0.6
0.5
0.4
0.3
0.2
0.1
0.0
****
Horizontal_Pol
Node output
Experiment
CT Hypoxia_high __Stiff_ECM:0
EXP Hypoxia_high __Stiff_ECM:1
Control sim.: Lv_28382476_Hypoxia_WT_Softer_ECM
Perturb. sim.: Lv_28382476_Hypoxia_WT_Stiff_ECM

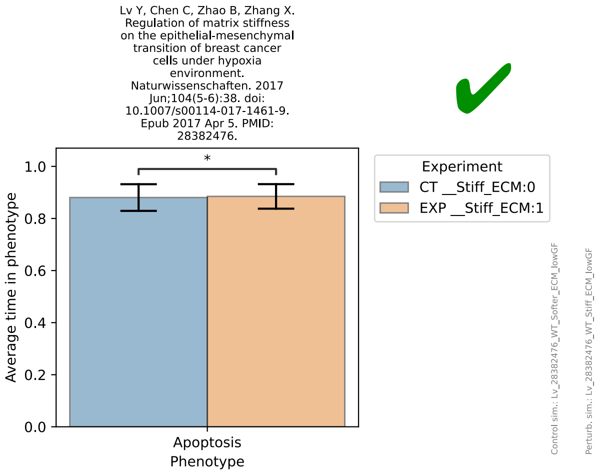
Lv Y, Chen C, Zhao B, Zhang X. Regulation of matrix stiffness on the epithelial-mesenchymal transition of breast cancer cells under hypoxia environment. Naturwissenschaften. 2017 Jun;104(5-6):38. doi: 10.1007/s00114-017-1461-9. Epub 2017 Apr 5. PMID: 28382476.
✔
Average time in phenotype
1.0
0.8
0.6
0.4
0.2
0.0
*
Apoptosis
Phenotype
Experiment
CT __Stiff_ECM:0
EXP __Stiff_ECM:1
Control sim.: Lv_28382476_WT_Softer_ECM_lowGF
Perturb. sim.: Lv_28382476_WT_Stiff_ECM_lowGF

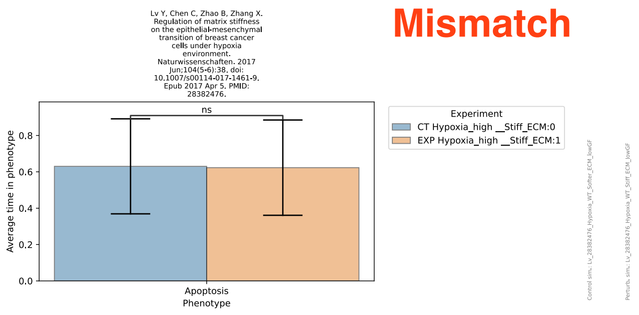
Lv Y, Chen C, Zhao B, Zhang X. Regulation of matrix stiffness on the epithelial-mesenchymal transition of breast cancer cells under hypoxia environment. Naturwissenschaften. 2017 Jun;104(5-6):38. doi: 10.1007/s00114-017-1461-9. Epub 2017 Apr 5. PMID: 28382476.
Mismatch
ns
Average time in phenotype
Apoptosis
Phenotype
Experiment
CT Hypoxia_high __Stiff_ECM:0
EXP Hypoxia_high __Stiff_ECM:1
Control sim.: Lv_28382476_Hypoxia_WT_Softer_ECM_lowGF
Perturb. sim.: Lv_28382476_Hypoxia_WT_Stiff_ECM_lowGF

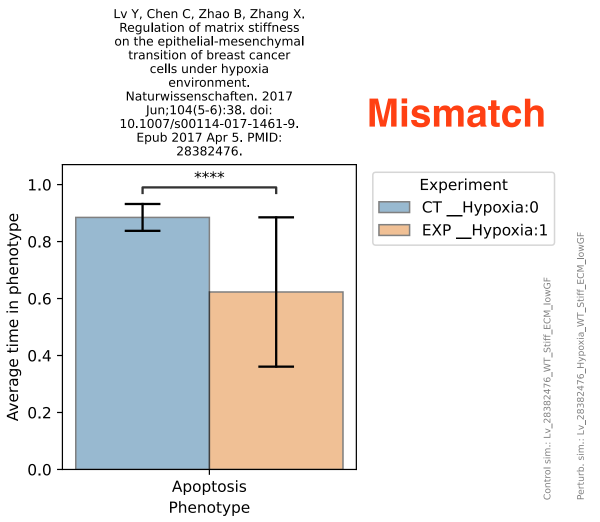
Lv Y, Chen C, Zhao B, Zhang X. Regulation of matrix stiffness on the epithelial-mesenchymal transition of breast cancer cells under hypoxia environment. Naturwissenschaften. 2017 Jun;104(5-6):38. doi: 10.1007/s00114-017-1461-9. Epub 2017 Apr 5. PMID: 28382476.
Mismatch
****
Average time in phenotype
Apoptosis
Phenotype
Experiment
CT __Hypoxia:0
EXP __Hypoxia:1
Control sim.: Lv_28382476_WT_Stiff_ECM_lowGF
Perturb. sim.: Lv_28382476_Hypoxia_WT_Stiff_ECM_lowGF

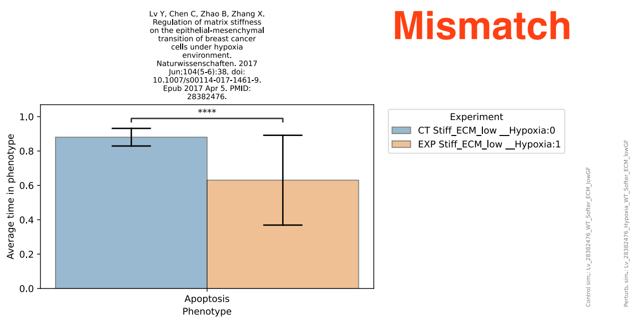
Lv Y, Chen C, Zhao B, Zhang X. Regulation of matrix stiffness on the epithelial-mesenchymal transition of breast cancer cells under hypoxia environment. Naturwissenschaften. 2017 Jun;104(5-6):38. doi: 10.1007/s00114-017-1461-9. Epub 2017 Apr 5. PMID: 28382476.
Mismatch
****
Average time in phenotype
Apoptosis
Phenotype
Experiment
CT Stiff_ECM_low __Hypoxia:0
EXP Stiff_ECM_low __Hypoxia:1
Control sim.: Lv_28382476_WT_Softer_ECM_lowGF
Perturb. sim.: Lv_28382476_Hypoxia_WT_Softer_ECM_lowGF

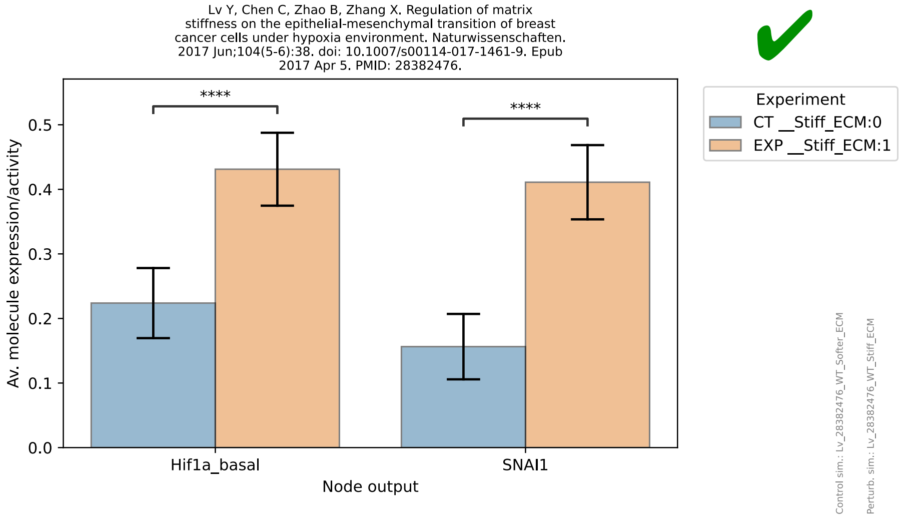
Lv Y, Chen C, Zhao B, Zhang X. Regulation of matrix stiffness on the epithelial-mesenchymal transition of breast cancer cells under hypoxia environment. Naturwissenschaften. 2017 Jun;104(5-6):38. doi: 10.1007/s00114-017-1461-9. Epub 2017 Apr 5. PMID: 28382476.
****
****
Av. molecule expression/activity
Hif1a_basal
SNAI1
Node output
Experiment
CT __Stiff_ECM:0
EXP __Stiff_ECM:1
Control sim.: Lv_28382476_WT_Softer_ECM
Perturb. sim.: Lv_28382476_WT_Stiff_ECM

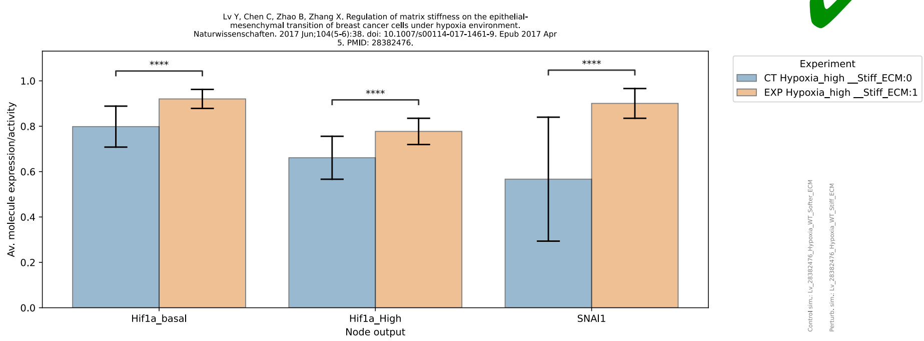
Lv Y, Chen C, Zhao B, Zhang X. Regulation of matrix stiffness on the epithelial-mesenchymal transition of breast cancer cells under hypoxia environment. Naturwissenschaften. 2017 Jun;104(5-6):38. doi: 10.1007/s00114-017-1461-9. Epub 2017 Apr 5. PMID: 28382476.
Av. molecule expression/activity
1.0
0.8
0.6
0.4
0.2
0.0
****
****
****
Hif1a_basal
Hif1a_High
SNAI1
Node output
Experiment
CT Hypoxia_high __Stiff_ECM:0
EXP Hypoxia_high __Stiff_ECM:1
Control sim.: Lv_28382476_Hypoxia_WT_Softer_ECM
Perturb. sim.: Lv_28382476_Hypoxia_WT_Stiff_ECM

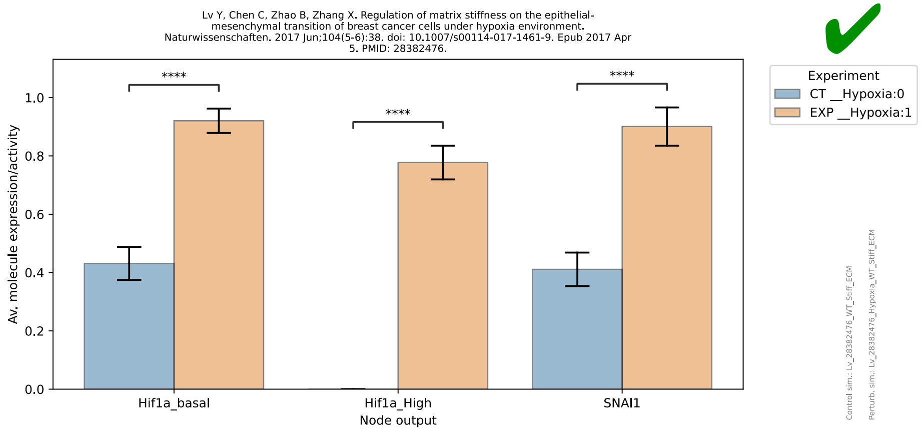
Lv Y, Chen C, Zhao B, Zhang X. Regulation of matrix stiffness on the epithelial-mesenchymal transition of breast cancer cells under hypoxia environment. Naturwissenschaften. 2017 Jun;104(5-6):38. doi: 10.1007/s00114-017-1461-9. Epub 2017 Apr 5. PMID: 28382476.
Av. molecule expression/activity
1.0
0.8
0.6
0.4
0.2
0.0
****
****
****
Hif1a_basal
Hif1a_High
SNAI1
Node output
Experiment
CT __Hypoxia:0
EXP __Hypoxia:1
Control sim.: Lv_28382476_WT_Stiff_ECM
Perturb. sim.: Lv_28382476_Hypoxia_WT_Stiff_ECM

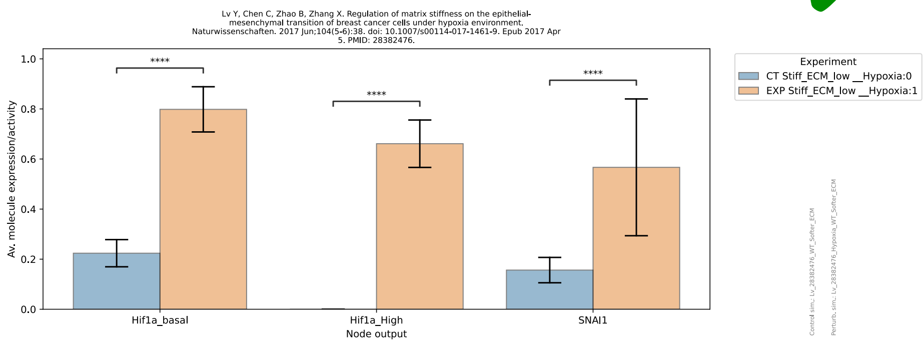
Lv Y, Chen C, Zhao B, Zhang X. Regulation of matrix stiffness on the epithelial-mesenchymal transition of breast cancer cells under hypoxia environment. Naturwissenschaften. 2017 Jun;104(5-6):38. doi: 10.1007/s00114-017-1461-9. Epub 2017 Apr 5. PMID: 28382476.
Av. molecule expression/activity
1.0
0.8
0.6
0.4
0.2
0.0
****
****
****
Hif1a_basal
Hif1a_High
SNAI1
Node output
Experiment
CT Stiff_ECM_low __Hypoxia:0
EXP Stiff_ECM_low __Hypoxia:1
Control sim.: Lv_28382476_WT_Softer_ECM
Perturb. sim.: Lv_28382476_Hypoxia_WT_Softer_ECM

Mismatch

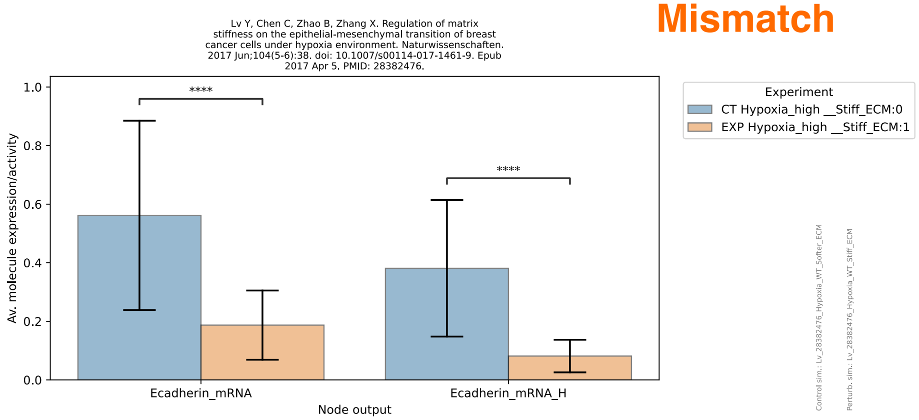
Lv Y, Chen C, Zhao B, Zhang X. Regulation of matrix stiffness on the epithelial-mesenchymal transition of breast cancer cells under hypoxia environment. Naturwissenschaften. 2017 Jun;104(5-6):38. doi: 10.1007/s00114-017-1461-9. Epub 2017 Apr 5. PMID: 28382476.
Av. molecule expression/activity
1.0
0.8
0.6
0.4
0.2
0.0
****
****
Ecadherin_mRNA
Ecadherin_mRNA_H
Node output
Experiment
CT Hypoxia_high __Stiff_ECM:0
EXP Hypoxia_high __Stiff_ECM:1
Control sim.: Lv_28382476_Hypoxia_WT_Softer_ECM
Perturb. sim.: Lv_28382476_Hypoxia_WT_Stiff_ECM

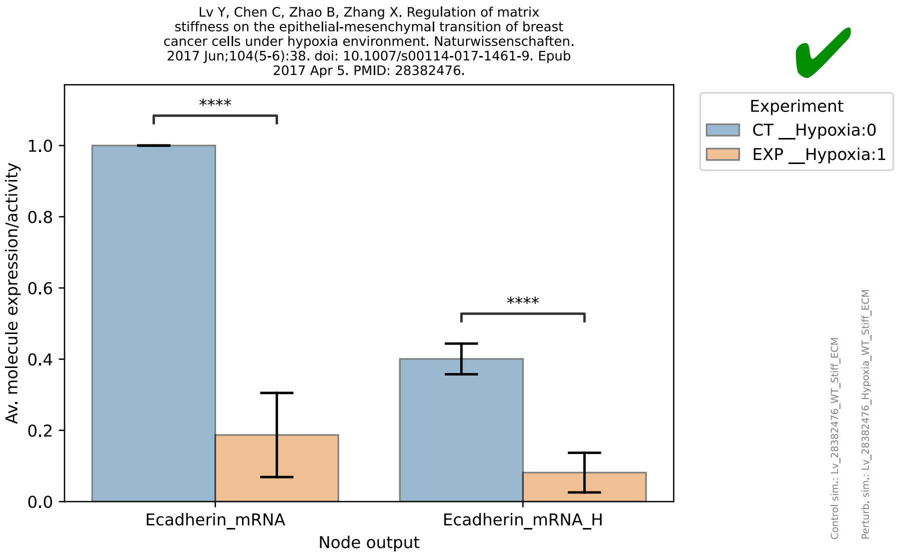
Lv Y, Chen C, Zhao B, Zhang X. Regulation of matrix stiffness on the epithelial-mesenchymal transition of breast cancer cells under hypoxia environment. Naturwissenschaften. 2017 Jun;104(5-6):38. doi: 10.1007/s00114-017-1461-9. Epub 2017 Apr 5. PMID: 28382476.
Av. molecule expression/activity
Node output
Ecadherin_mRNA
Ecadherin_mRNA_H
****
Experiment
CT __Hypoxia:0
EXP __Hypoxia:1
Control sim.: Lv_28382476_WT_Stiff_ECM
Perturb. sim.: Lv_28382476_Hypoxia_WT_Stiff_ECM

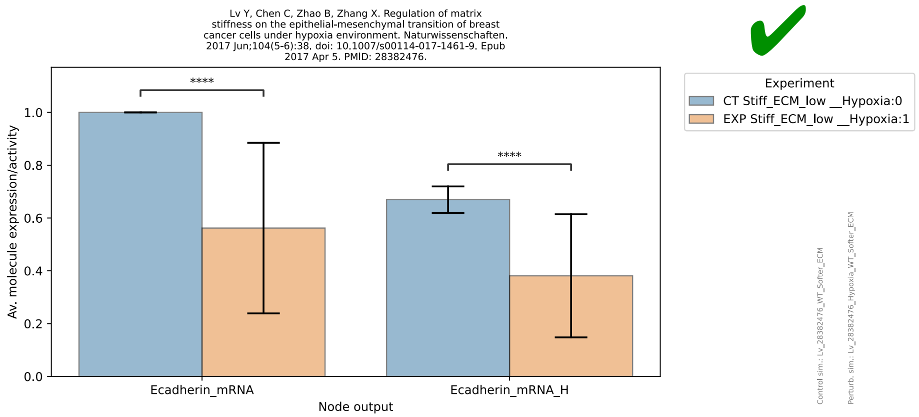
Lv Y, Chen C, Zhao B, Zhang X. Regulation of matrix stiffness on the epithelial-mesenchymal transition of breast cancer cells under hypoxia environment. Naturwissenschaften. 2017 Jun;104(5-6):38. doi: 10.1007/s00114-017-1461-9. Epub 2017 Apr 5. PMID: 28382476.
Av. molecule expression/activity
Node output
Ecadherin_mRNA
Ecadherin_mRNA_H
****
Experiment
CT Stiff_ECM_low __Hypoxia:0
EXP Stiff_ECM_low __Hypoxia:1
Control sim.: Lv_28382476_WT_Softer_ECM
Perturb. sim.: Lv_28382476_Hypoxia_WT_Softer_ECM

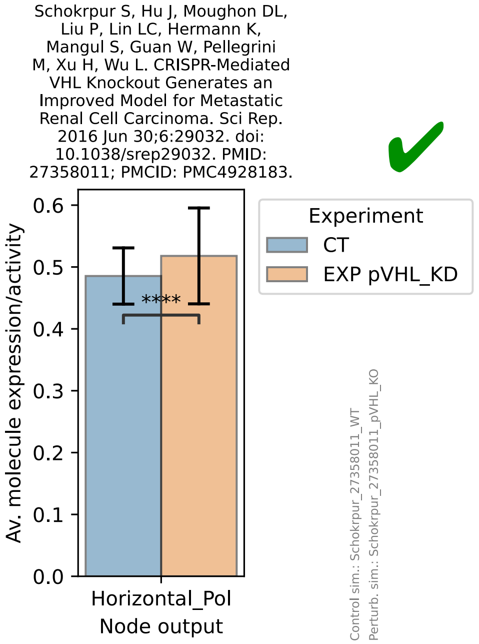
Schokrpur S, Hu J, Moughon DL, Liu P, Lin LC, Hermann K, Mangul S, Guan W, Pellegrini M, Xu H, Wu L. CRISPR-Mediated VHL Knockout Generates an Improved Model for Metastatic Renal Cell Carcinoma. Sci Rep. 2016 Jun 30;6:29032. doi: 10.1038/srep29032. PMID: 27358011; PMCID: PMC4928183.
Av. molecule expression/activity
Horizontal_Pol
Node output
****
Experiment
CT
EXP pVHL_KD
Control sim.: Schokrpur_27358011_WT
Perturb. sim.: Schokrpur_27358011_pVHL_KO

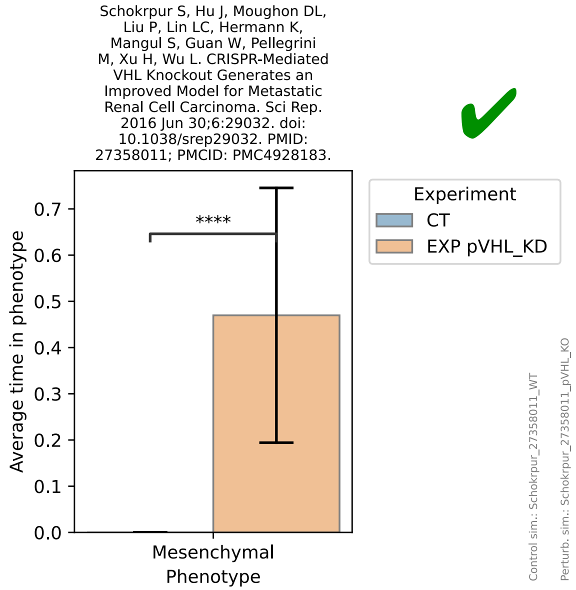
Schokrpur S, Hu J, Moughon DL, Liu P, Lin LC, Hermann K, Mangul S, Guan W, Pellegrini M, Xu H, Wu L. CRISPR-Mediated VHL Knockout Generates an Improved Model for Metastatic Renal Cell Carcinoma. Sci Rep. 2016 Jun 30;6:29032. doi: 10.1038/srep29032. PMID: 27358011; PMCID: PMC4928183.
Average time in phenotype
Mesenchymal
Phenotype
****
Experiment
CT
EXP pVHL_KD
Control sim.: Schokrpur_27358011_WT
Perturb. sim.: Schokrpur_27358011_pVHL_KO

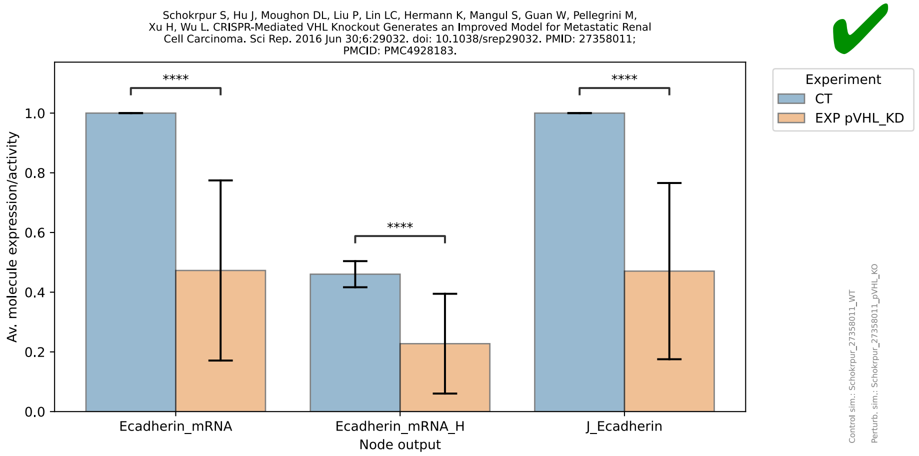
Schokrpur S, Hu J, Moughon DL, Liu P, Lin LC, Hermann K, Mangul S, Guan W, Pellegrini M,
Xu H, Wu L. CRISPR-Mediated VHL Knockout Generates an Improved Model for Metastatic Renal
Cell Carcinoma. Sci Rep. 2016 Jun 30;6:29032. doi: 10.1038/srep29032. PMID: 27358011;
PMCID: PMC4928183.
Av. molecule expression/activity
1.0
0.8
0.6
0.4
0.2
0.0
****
****
****
Ecadherin_mRNA
Ecadherin_mRNA_H
J_Ecadherin
Node output
Experiment
CT
EXP pVHL_KD
✔
Control sim.: Schokrpur_27358011_WT
Perturb. sim.: Schokrpur_27358011_pVHL_KO

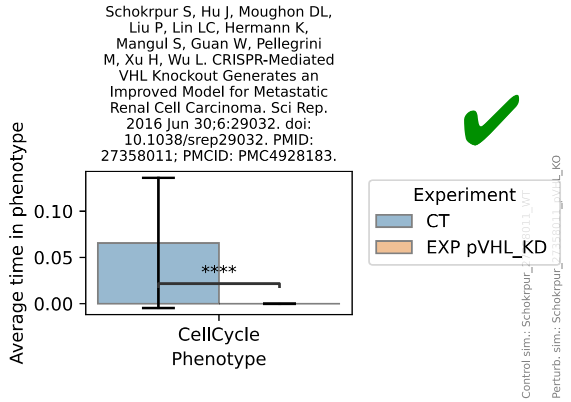
Schokrpur S, Hu J, Moughon DL,
Liu P, Lin LC, Hermann K,
Mangul S, Guan W, Pellegrini
M, Xu H, Wu L. CRISPR-Mediated
VHL Knockout Generates an
Improved Model for Metastatic
Renal Cell Carcinoma. Sci Rep.
2016 Jun 30;6:29032. doi:
10.1038/srep29032. PMID:
27358011; PMCID: PMC4928183.
Average time in phenotype
0.10
0.05
0.00
****
CellCycle
Phenotype
Experiment
CT
EXP pVHL_KD
✔

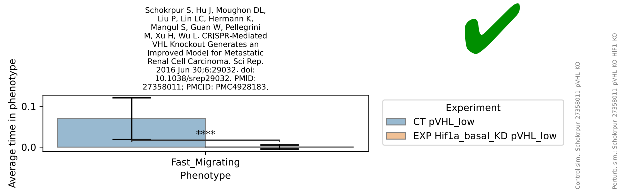
Schokrpur S, Hu J, Moughon DL,
Liu P, Lin LC, Hermann K,
Mangul S, Guan W, Pellegrini
M, Xu H, Wu L. CRISPR-Mediated
VHL Knockout Generates an
Improved Model for Metastatic
Renal Cell Carcinoma. Sci Rep.
2016 Jun 30;6:29032. doi:
10.1038/srep29032. PMID:
27358011; PMCID: PMC4928183.
Average time in phenotype
0.1
0.0
****
Fast_Migrating
Phenotype
Experiment
CT pVHL_low
EXP Hif1a_basal_KD pVHL_low
✔

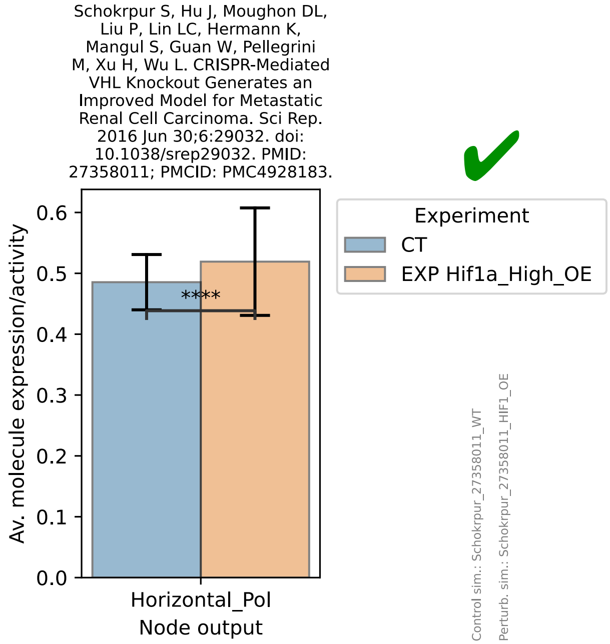
Schokrpur S, Hu J, Moughon DL,
Liu P, Lin LC, Hermann K,
Mangul S, Guan W, Pellegrini
M, Xu H, Wu L. CRISPR-Mediated
VHL Knockout Generates an
Improved Model for Metastatic
Renal Cell Carcinoma. Sci Rep.
2016 Jun 30;6:29032. doi:
10.1038/srep29032. PMID:
27358011; PMCID: PMC4928183.
Av. molecule expression/activity
0.6
0.5
0.4
0.3
0.2
0.1
0.0
****
Horizontal_Pol
Node output
Experiment
CT
EXP Hif1a_High_OE
✔
Control sim.: Schokrpur_27358011_WT
Perturb. sim.: Schokrpur_27358011_HIF1_OE

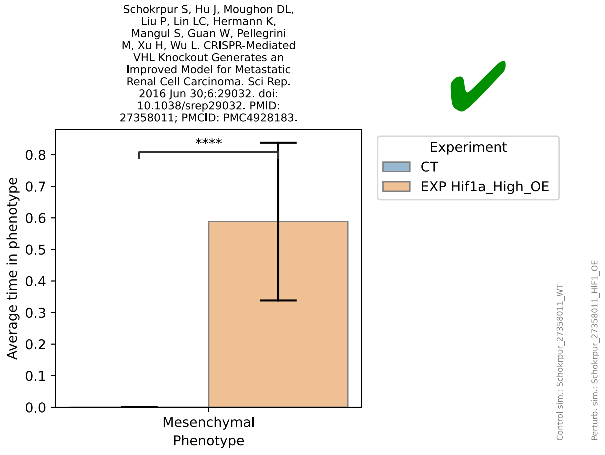
Schokrpur S, Hu J, Moughon DL, Liu P, Lin LC, Hermann K, Mangul S, Guan W, Pellegrini M, Xu H, Wu L. CRISPR-Mediated VHL Knockout Generates an Improved Model for Metastatic Renal Cell Carcinoma. Sci Rep. 2016 Jun 30;6:29032. doi: 10.1038/srep29032. PMID: 27358011; PMCID: PMC4928183.
Average time in phenotype
****
Experiment
CT
EXP Hif1a_High_OE
Mesenchymal Phenotype
Control sim.: Schokrpur_27358011_WT
Perturb. sim.: Schokrpur_27358011_HIF1_OE

Mismatch

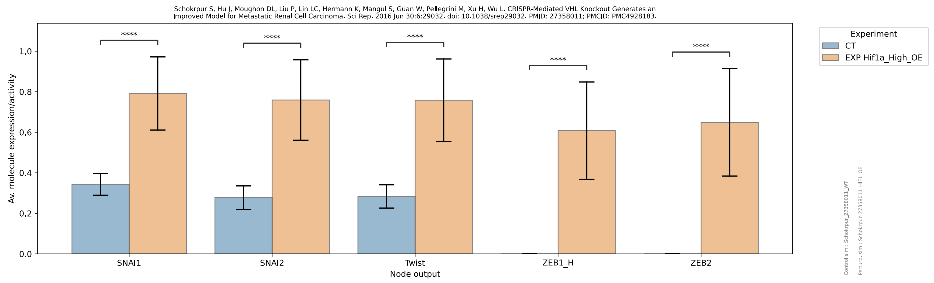
Av. molecule expression/activity
Experiment
CT
EXP Hif1a_High_OE
SNAI1
SNAI2
Twist
ZEB1_H
ZEB2
Node output

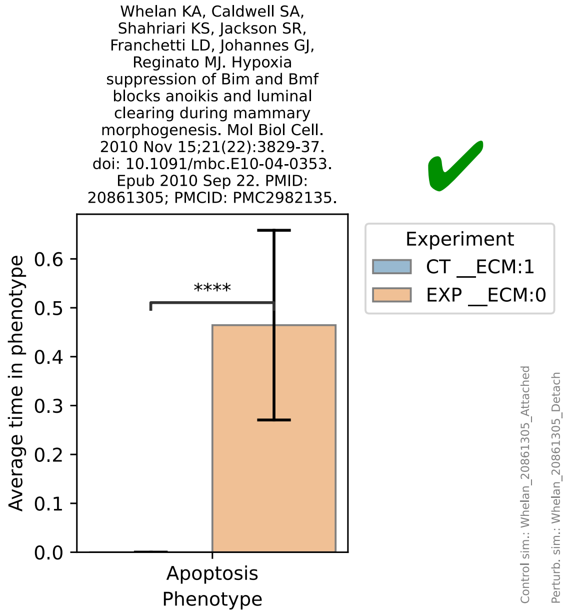
Whelan KA, Caldwell SA, Shahriari KS, Jackson SR, Franchetti LD, Johannes GJ, Reginato MJ. Hypoxia suppression of Bim and Bmf blocks anoikis and luminal clearing during mammary morphogenesis. Mol Biol Cell. 2010 Nov 15;21(22):3829-37. doi: 10.1091/mbc.E10-04-0353. Epub 2010 Sep 22. PMID: 20861305; PMCID: PMC2982135.
Average time in phenotype
****
Experiment
CT __ECM:1
EXP __ECM:0
Apoptosis Phenotype
Control sim.: Whelan_20861305_Attached
Perturb. sim.: Whelan_20861305_Detach

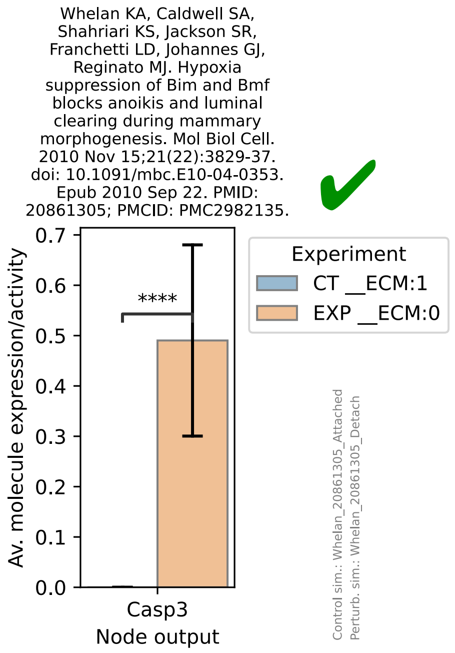
Whelan KA, Caldwell SA, Shahriari KS, Jackson SR, Franchetti LD, Johannes GJ, Reginato MJ. Hypoxia suppression of Bim and Bmf blocks anoikis and luminal clearing during mammary morphogenesis. Mol Biol Cell. 2010 Nov 15;21(22):3829-37. doi: 10.1091/mbc.E10-04-0353. Epub 2010 Sep 22. PMID: 20861305; PMCID: PMC2982135.
Av. molecule expression/activity
****
Experiment
CT __ECM:1
EXP __ECM:0
Casp3 Node output
Control sim.: Whelan_20861305_Attached
Perturb. sim.: Whelan_20861305_Detach

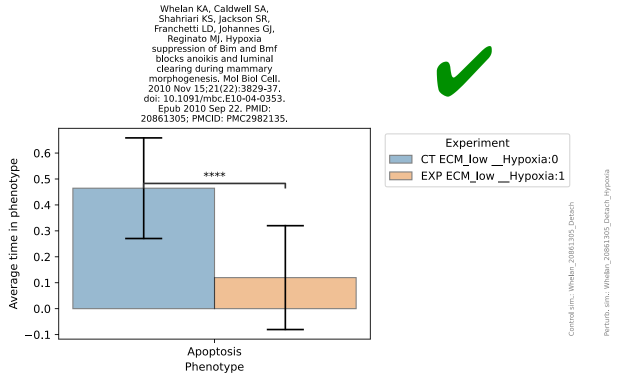
Whelan KA, Caldwell SA, Shahriari KS, Jackson SR, Franchetti LD, Johannes GJ, Reginato MJ. Hypoxia suppression of Bim and Bmf blocks anoikis and luminal clearing during mammary morphogenesis. Mol Biol Cell. 2010 Nov 15;21(22):3829-37. doi: 10.1091/mbc.E10-04-0353. Epub 2010 Sep 22. PMID: 20861305; PMCID: PMC2982135.
Average time in phenotype
****
Apoptosis
Phenotype
Experiment
CT ECM_low __Hypoxia:0
EXP ECM_low __Hypoxia:1
Control sim.: Whelan_20861305_Detach
Perturb. sim.: Whelan_20861305_Detach_Hypoxia

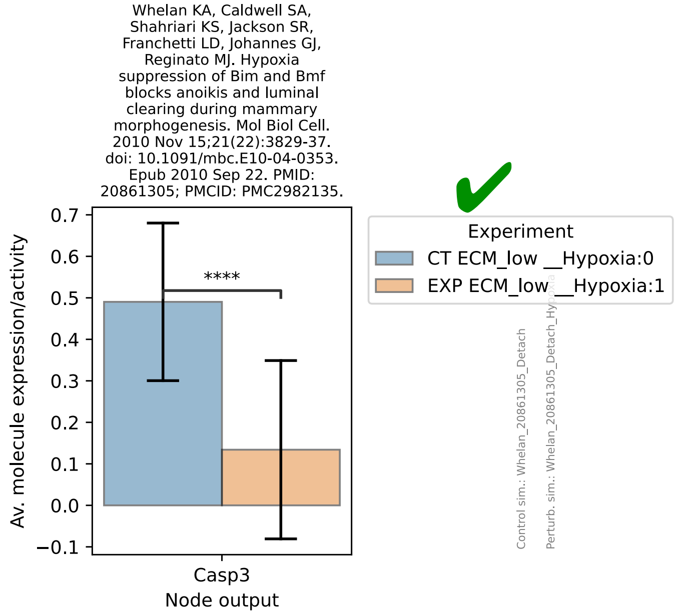
Whelan KA, Caldwell SA, Shahriari KS, Jackson SR, Franchetti LD, Johannes GJ, Reginato MJ. Hypoxia suppression of Bim and Bmf blocks anoikis and luminal clearing during mammary morphogenesis. Mol Biol Cell. 2010 Nov 15;21(22):3829-37. doi: 10.1091/mbc.E10-04-0353. Epub 2010 Sep 22. PMID: 20861305; PMCID: PMC2982135.
Av. molecule expression/activity
****
Casp3
Node output
Experiment
CT ECM_low __Hypoxia:0
EXP ECM_low __Hypoxia:1
Control sim.: Whelan_20861305_Detach

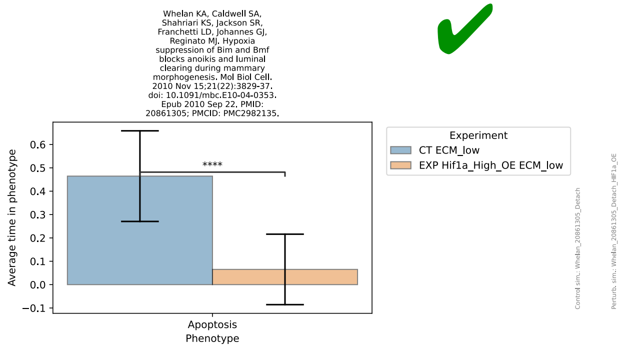
Whelan KA, Caldwell SA, Shahriari KS, Jackson SR, Franchetti LD, Johannes GJ, Reginato MJ. Hypoxia suppression of Bim and Bmf blocks anoikis and luminal clearing during mammary morphogenesis. Mol Biol Cell. 2010 Nov 15;21(22):3829-37. doi: 10.1091/mbc.E10-04-0353. Epub 2010 Sep 22. PMID: 20861305; PMCID: PMC2982135.
Average time in phenotype
****
Apoptosis
Phenotype
Experiment
CT ECM_low
EXP Hif1a_High_OE ECM_low
Control sim.: Whelan_20861305_Detach
Perturb. sim.: Whelan_20861305_Detach_Hif1a_OE

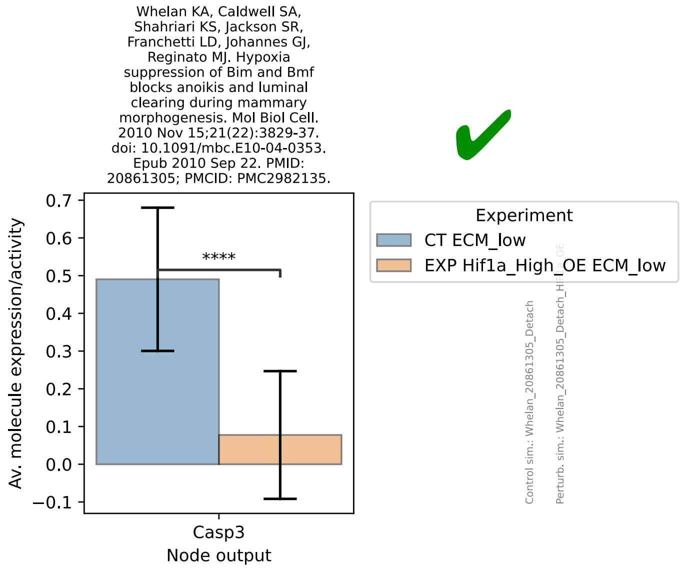
Whelan KA, Caldwell SA, Shahriari KS, Jackson SR, Franchetti LD, Johannes GJ, Reginato MJ. Hypoxia suppression of Bim and Bmf blocks anoikis and luminal clearing during mammary morphogenesis. Mol Biol Cell. 2010 Nov 15;21(22):3829-37. doi: 10.1091/mbc.E10-04-0353. Epub 2010 Sep 22. PMID: 20861305; PMCID: PMC2982135.
Av. molecule expression/activity
****
Casp3
Node output
Experiment
CT ECM_low
EXP Hif1a_High_OE ECM_low
Control sim.: Whelan_20861305_Detach

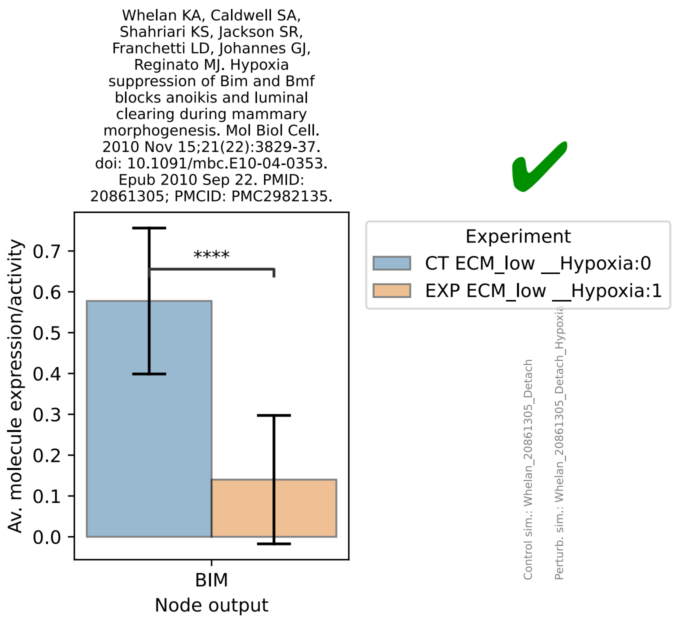
Whelan KA, Caldwell SA, Shahriari KS, Jackson SR, Franchetti LD, Johannes GJ, Reginato MJ. Hypoxia suppression of Bim and Bmf blocks anoikis and luminal clearing during mammary morphogenesis. Mol Biol Cell. 2010 Nov 15;21(22):3829-37. doi: 10.1091/mbc.E10-04-0353. Epub 2010 Sep 22. PMID: 20861305; PMCID: PMC2982135.
Av. molecule expression/activity
0.0
0.1
0.2
0.3
0.4
0.5
0.6
0.7
****
BIM
Node output
Experiment
CT ECM_low __Hypoxia:0
EXP ECM_low __Hypoxia:1
Control sim.: Whelan_20861305_Detach
Perturb. sim.: Whelan_20861305_Detach_Hypoxi

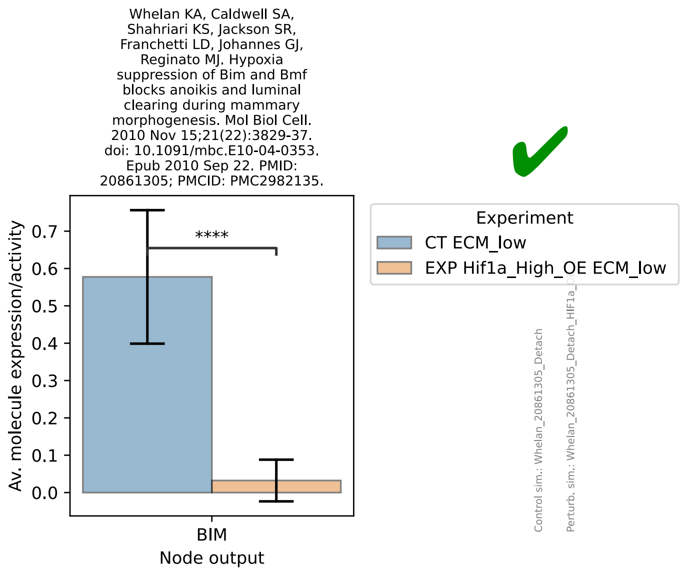
Whelan KA, Caldwell SA, Shahriari KS, Jackson SR, Franchetti LD, Johannes GJ, Reginato MJ. Hypoxia suppression of Bim and Bmf blocks anoikis and luminal clearing during mammary morphogenesis. Mol Biol Cell. 2010 Nov 15;21(22):3829-37. doi: 10.1091/mbc.E10-04-0353. Epub 2010 Sep 22. PMID: 20861305; PMCID: PMC2982135.
Av. molecule expression/activity
0.0
0.1
0.2
0.3
0.4
0.5
0.6
0.7
****
BIM
Node output
Experiment
CT ECM_low
EXP Hif1a_High_OE ECM_low
Control sim.: Whelan_20861305_Detach
Perturb. sim.: Whelan_20861305_Detach_HIF1a_

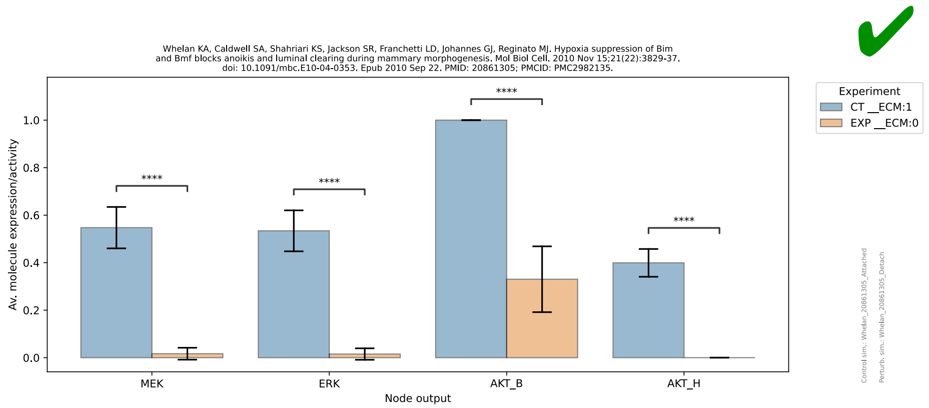
Whelan KA, Caldwell SA, Shahriari KS, Jackson SR, Franchetti LD, Johannes GJ, Reginato MJ. Hypoxia suppression of Bim and Bmf blocks anoikis and luminal clearing during mammary morphogenesis. Mol Biol Cell. 2010 Nov 15;21(22):3829-37. doi: 10.1091/mbc.E10-04-0353. Epub 2010 Sep 22. PMID: 20861305; PMCID: PMC2982135.
Av. molecule expression/activity
0.0
0.2
0.4
0.6
0.8
1.0
****
MEK
ERK
AKT_B
AKT_H
Node output
Experiment
CT __ECM:1
EXP __ECM:0
Control sim.: Whelan_20861305_Attached
Perturb. sim.: Whelan_20861305_Detach

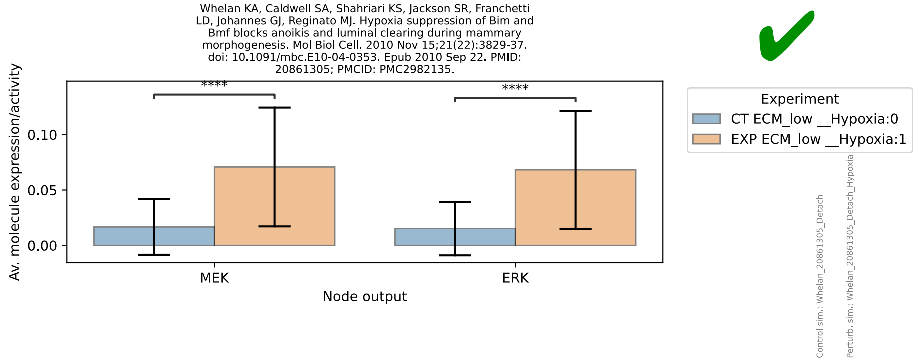
Whelan KA, Caldwell SA, Shahriari KS, Jackson SR, Franchetti LD, Johannes GJ, Reginato MJ. Hypoxia suppression of Bim and Bmf blocks anoikis and luminal clearing during mammary morphogenesis. Mol Biol Cell. 2010 Nov 15;21(22):3829-37. doi: 10.1091/mbc.E10-04-0353. Epub 2010 Sep 22. PMID: 20861305; PMCID: PMC2982135.
Av. molecule expression/activity
0.00
0.05
0.10
****
MEK
ERK
Node output
Experiment
CT ECM_low __Hypoxia:0
EXP ECM_low __Hypoxia:1
Control sim.: Whelan_20861305_Detach
Perturb. sim.: Whelan_20861305_Detach_Hypoxia

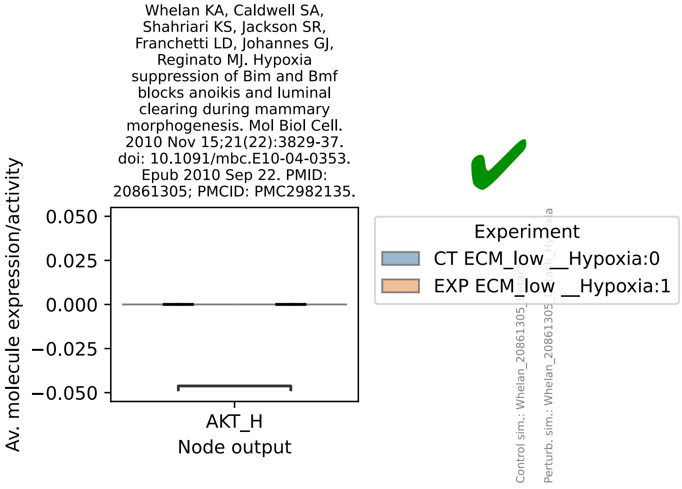

Whelan KA, Caldwell SA, Shahriari KS, Jackson SR, Franchetti LD, Johannes GJ, Reginato MJ. Hypoxia suppression of Bim and Bmf blocks anoikis and luminal clearing during mammary morphogenesis. Mol Biol Cell. 2010 Nov 15;21(22):3829-37. doi: 10.1091/mbc.E10-04-0353. Epub 2010 Sep 22. PMID: 20861305; PMCID: PMC2982135.
Av. molecule expression/activity
0.050
0.025
0.000
−0.025
−0.050
AKT_H
Node output
Experiment
CT ECM_low __Hypoxia:0
EXP ECM_low __Hypoxia:1


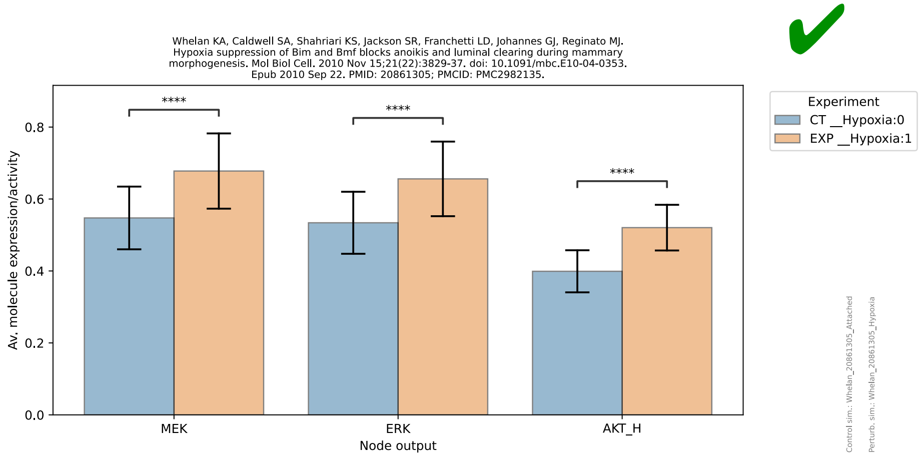

Whelan KA, Caldwell SA, Shahriari KS, Jackson SR, Franchetti LD, Johannes GJ, Reginato MJ. Hypoxia suppression of Bim and Bmf blocks anoikis and luminal clearing during mammary morphogenesis. Mol Biol Cell. 2010 Nov 15;21(22):3829-37. doi: 10.1091/mbc.E10-04-0353. Epub 2010 Sep 22. PMID: 20861305; PMCID: PMC2982135.
Av. molecule expression/activity
0.0
0.2
0.4
0.6
0.8
****
****
****
MEK
ERK
AKT_H
Node output
Experiment
CT __Hypoxia:0
EXP __Hypoxia:1
Control sim.: Whelan_20861305_Attached
Perturb. sim.: Whelan_20861305_Hypoxia


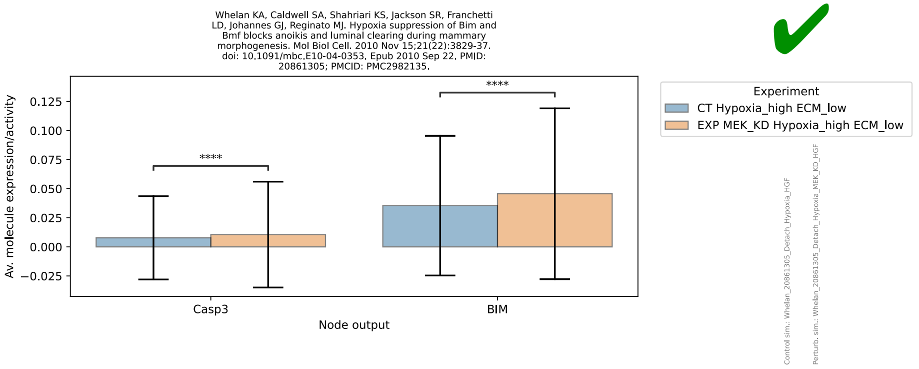

Whelan KA, Caldwell SA, Shahriari KS, Jackson SR, Franchetti LD, Johannes GJ, Reginato MJ. Hypoxia suppression of Bim and Bmf blocks anoikis and luminal clearing during mammary morphogenesis. Mol Biol Cell. 2010 Nov 15;21(22):3829-37. doi: 10.1091/mbc.E10-04-0353. Epub 2010 Sep 22. PMID: 20861305; PMCID: PMC2982135.
Av. molecule expression/activity
0.125
0.100
0.075
0.050
0.025
0.000
−0.025
****
****
Casp3
BIM
Node output
Experiment
CT Hypoxia_high ECM_low
EXP MEK_KD Hypoxia_high ECM_low
Control sim.: Whelan_20861305_Detach_Hypoxia_HGF
Perturb. sim.: Whelan_20861305_Detach_Hypoxia_MEK_KD_HGF


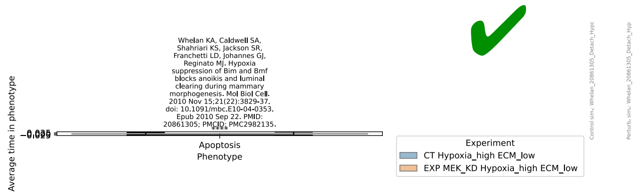

Whelan KA, Caldwell SA, Shahriari KS, Jackson SR, Franchetti LD, Johannes GJ, Reginato MJ. Hypoxia suppression of Bim and Bmf blocks anoikis and luminal clearing during mammary morphogenesis. Mol Biol Cell. 2010 Nov 15;21(22):3829-37. doi: 10.1091/mbc.E10-04-0353. Epub 2010 Sep 22. PMID: 20861305; PMCID: PMC2982135.
Average time in phenotype
Apoptosis
Phenotype
Experiment
CT Hypoxia_high ECM_low
EXP MEK_KD Hypoxia_high ECM_low

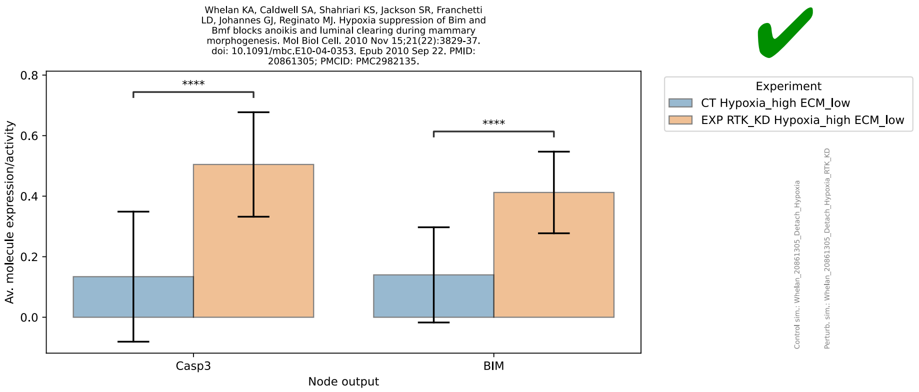
Whelan KA, Caldwell SA, Shahriari KS, Jackson SR, Franchetti LD, Johannes GJ, Reginato MJ. Hypoxia suppression of Bim and Bmf blocks anoikis and luminal clearing during mammary morphogenesis. Mol Biol Cell. 2010 Nov 15;21(22):3829-37. doi: 10.1091/mbc.E10-04-0353. Epub 2010 Sep 22. PMID: 20861305; PMCID: PMC2982135.
Av. molecule expression/activity
****
****
Casp3
BIM
Node output
Experiment
CT Hypoxia_high ECM_low
EXP RTK_KD Hypoxia_high ECM_low
Control sim.: Whelan_20861305_Detach_Hypoxia
Perturb. sim.: Whelan_20861305_Detach_Hypoxia_RTK_KD

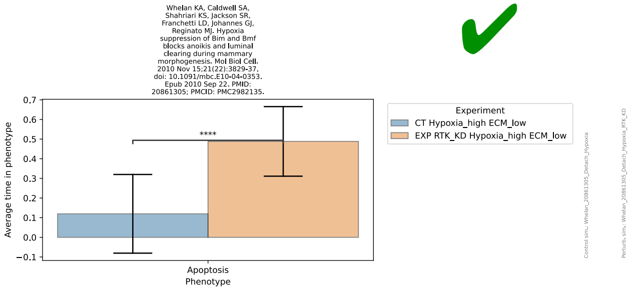
Whelan KA, Caldwell SA, Shahriari KS, Jackson SR, Franchetti LD, Johannes GJ, Reginato MJ. Hypoxia suppression of Bim and Bmf blocks anoikis and luminal clearing during mammary morphogenesis. Mol Biol Cell. 2010 Nov 15;21(22):3829-37. doi: 10.1091/mbc.E10-04-0353. Epub 2010 Sep 22. PMID: 20861305; PMCID: PMC2982135.
Average time in phenotype
****
Apoptosis
Phenotype
Experiment
CT Hypoxia_high ECM_low
EXP RTK_KD Hypoxia_high ECM_low

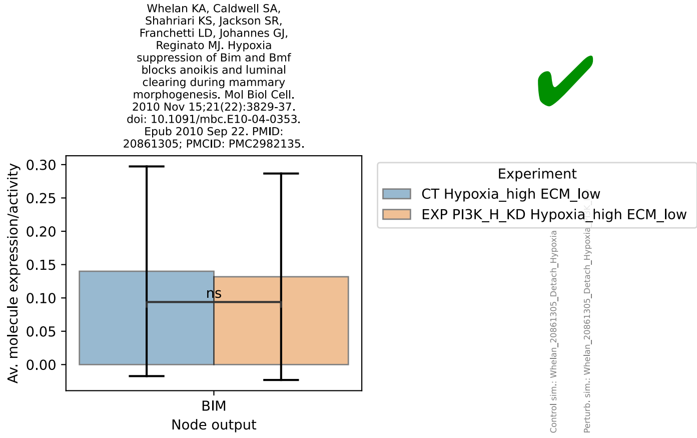
Whelan KA, Caldwell SA, Shahriari KS, Jackson SR, Franchetti LD, Johannes GJ, Reginato MJ. Hypoxia suppression of Bim and Bmf blocks anoikis and luminal clearing during mammary morphogenesis. Mol Biol Cell. 2010 Nov 15;21(22):3829-37. doi: 10.1091/mbc.E10-04-0353. Epub 2010 Sep 22. PMID: 20861305; PMCID: PMC2982135.
Av. molecule expression/activity
ns
BIM
Node output
Experiment
CT Hypoxia_high ECM_low
EXP PI3K_H_KD Hypoxia_high ECM_low
Control sim.: Whelan_20861305_Detach_Hypoxia
Perturb. sim.: Whelan_20861305_Detach_Hypoxia

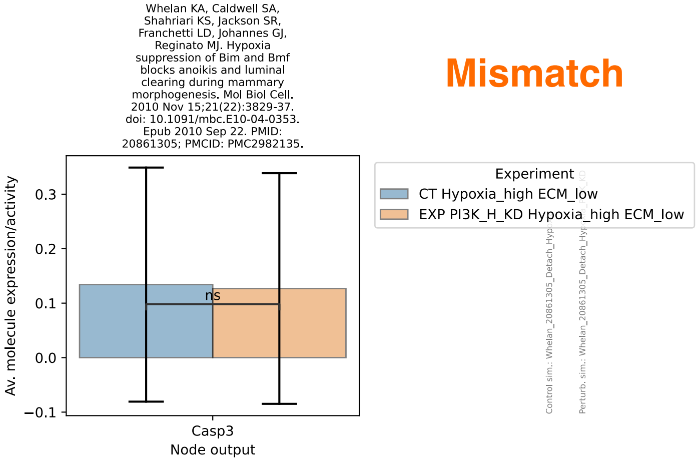
Whelan KA, Caldwell SA, Shahriari KS, Jackson SR, Franchetti LD, Johannes GJ, Reginato MJ. Hypoxia suppression of Bim and Bmf blocks anoikis and luminal clearing during mammary morphogenesis. Mol Biol Cell. 2010 Nov 15;21(22):3829-37. doi: 10.1091/mbc.E10-04-0353. Epub 2010 Sep 22. PMID: 20861305; PMCID: PMC2982135.
Mismatch
Av. molecule expression/activity
ns
Casp3
Node output
Experiment
CT Hypoxia_high ECM_low
EXP PI3K_H_KD Hypoxia_high ECM_low

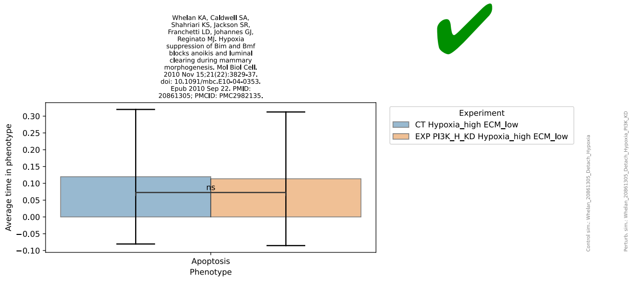
Whelan KA, Caldwell SA, Shahriari KS, Jackson SR, Franchetti LD, Johannes GJ, Reginato MJ. Hypoxia suppression of Bim and Bmf blocks anoikis and luminal clearing during mammary morphogenesis. Mol Biol Cell. 2010 Nov 15;21(22):3829-37. doi: 10.1091/mbc.E10-04-0353. Epub 2010 Sep 22. PMID: 20861305; PMCID: PMC2982135.
Average time in phenotype
0.30
0.25
0.20
0.15
0.10
0.05
0.00
−0.05
−0.10
ns
Apoptosis
Phenotype
Experiment
CT Hypoxia_high ECM_low
EXP PI3K_H_KD Hypoxia_high ECM_low
Control sim.: Whelan_20861305_Detach_Hypoxia
Perturb. sim.: Whelan_20861305_Detach_Hypoxia_PI3K_KD

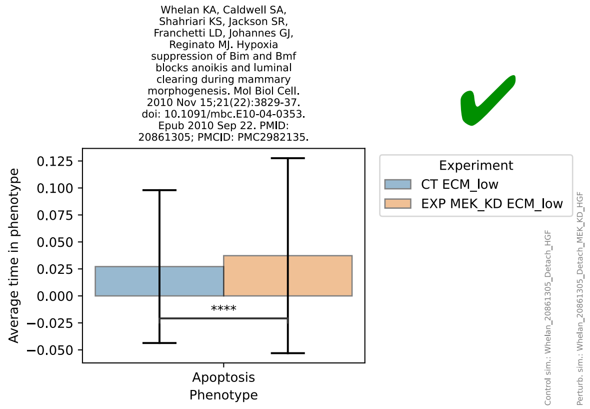
Whelan KA, Caldwell SA, Shahriari KS, Jackson SR, Franchetti LD, Johannes GJ, Reginato MJ. Hypoxia suppression of Bim and Bmf blocks anoikis and luminal clearing during mammary morphogenesis. Mol Biol Cell. 2010 Nov 15;21(22):3829-37. doi: 10.1091/mbc.E10-04-0353. Epub 2010 Sep 22. PMID: 20861305; PMCID: PMC2982135.
Average time in phenotype
0.125
0.100
0.075
0.050
0.025
0.000
−0.025
−0.050
****
Apoptosis
Phenotype
Experiment
CT ECM_low
EXP MEK_KD ECM_low
Control sim.: Whelan_20861305_Detach_HGF
Perturb. sim.: Whelan_20861305_Detach_MEK_KD_HGF

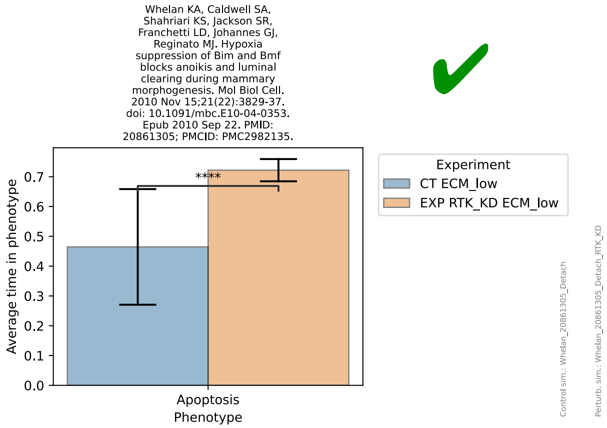
Whelan KA, Caldwell SA, Shahriari KS, Jackson SR, Franchetti LD, Johannes GJ, Reginato MJ. Hypoxia suppression of Bim and Bmf blocks anoikis and luminal clearing during mammary morphogenesis. Mol Biol Cell. 2010 Nov 15;21(22):3829-37. doi: 10.1091/mbc.E10-04-0353. Epub 2010 Sep 22. PMID: 20861305; PMCID: PMC2982135.
Average time in phenotype
0.7
0.6
0.5
0.4
0.3
0.2
0.1
0.0
****
Apoptosis
Phenotype
Experiment
CT ECM_low
EXP RTK_KD ECM_low
Control sim.: Whelan_20861305_Detach
Perturb. sim.: Whelan_20861305_Detach_RTK_KD

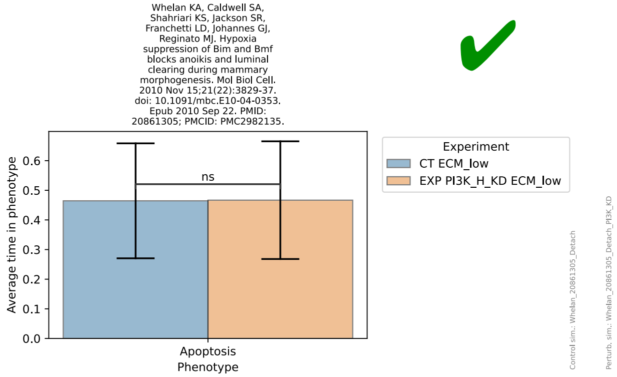
Whelan KA, Caldwell SA, Shahriari KS, Jackson SR, Franchetti LD, Johannes GJ, Reginato MJ. Hypoxia suppression of Bim and Bmf blocks anoikis and luminal clearing during mammary morphogenesis. Mol Biol Cell. 2010 Nov 15;21(22):3829-37. doi: 10.1091/mbc.E10-04-0353. Epub 2010 Sep 22. PMID: 20861305; PMCID: PMC2982135.
Average time in phenotype
0.6
0.5
0.4
0.3
0.2
0.1
0.0
ns
Apoptosis
Phenotype
Experiment
CT ECM_low
EXP PI3K_H_KD ECM_low
Control sim.: Whelan_20861305_Detach
Perturb. sim.: Whelan_20861305_Detach_PI3K_KD

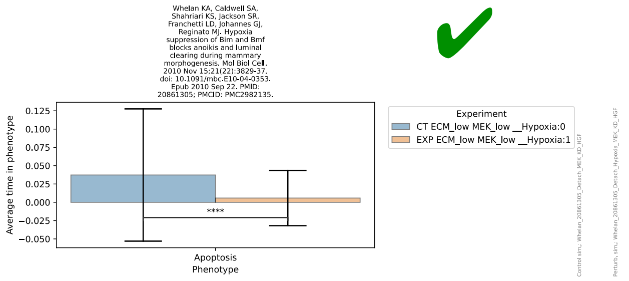
Whelan KA, Caldwell SA, Shahriari KS, Jackson SR, Franchetti LD, Johannes GJ, Reginato MJ. Hypoxia suppression of Bim and Bmf blocks anoikis and luminal clearing during mammary morphogenesis. Mol Biol Cell. 2010 Nov 15;21(22):3829-37. doi: 10.1091/mbc.E10-04-0353. Epub 2010 Sep 22. PMID: 20861305; PMCID: PMC2982135.
Average time in phenotype
0.125
0.100
0.075
0.050
0.025
0.000
−0.025
−0.050
****
Apoptosis
Phenotype
Experiment
CT ECM_low MEK_low __Hypoxia:0
EXP ECM_low MEK_low __Hypoxia:1
Control sim.: Whelan_20861305_Detach_MEK_KD_HGF
Perturb. sim.: Whelan_20861305_Detach_Hypoxia_MEK_KD_HGF

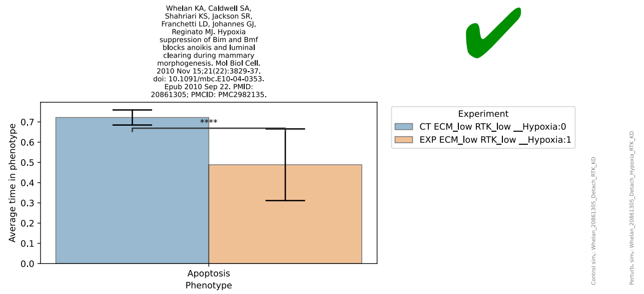
Whelan KA, Caldwell SA, Shahriari KS, Jackson SR, Franchetti LD, Johannes GJ, Reginato MJ. Hypoxia suppression of Bim and Bmf blocks anoikis and luminal clearing during mammary morphogenesis. Mol Biol Cell. 2010 Nov 15;21(22):3829-37. doi: 10.1091/mbc.E10-04-0353. Epub 2010 Sep 22. PMID: 20861305; PMCID: PMC2982135.
Average time in phenotype
0.7
0.6
0.5
0.4
0.3
0.2
0.1
0.0
****
Apoptosis
Phenotype
Experiment
CT ECM_low RTK_low __Hypoxia:0
EXP ECM_low RTK_low __Hypoxia:1
Control sim.: Whelan_20861305_Detach_RTK_KD
Perturb. sim.: Whelan_20861305_Detach_Hypoxia_RTK_KD

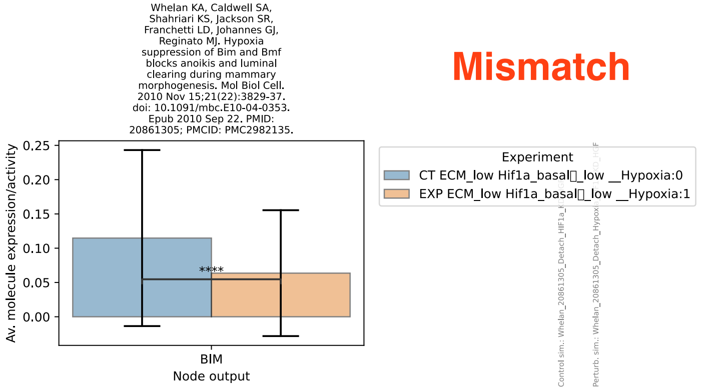
Whelan KA, Caldwell SA, Shahriari KS, Jackson SR, Franchetti LD, Johannes GJ, Reginato MJ. Hypoxia suppression of Bim and Bmf blocks anoikis and luminal clearing during mammary morphogenesis. Mol Biol Cell. 2010 Nov 15;21(22):3829-37. doi: 10.1091/mbc.E10-04-0353. Epub 2010 Sep 22. PMID: 20861305; PMCID: PMC2982135.
Av. molecule expression/activity
0.25
0.20
0.15
0.10
0.05
0.00
****
BIM
Node output
Experiment
CT ECM_low Hif1a_basal□_low __Hypoxia:0
EXP ECM_low Hif1a_basal□_low __Hypoxia:1

Mismatch

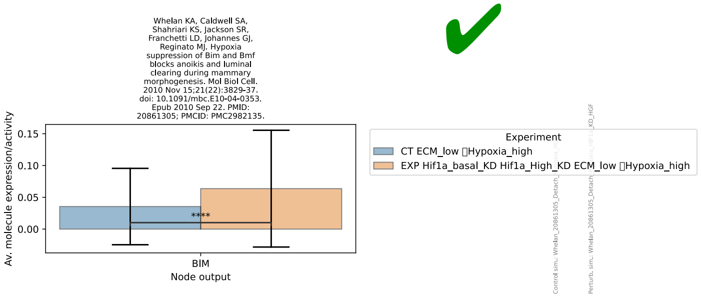
Whelan KA, Caldwell SA, Shahriari KS, Jackson SR, Franchetti LD, Johannes GJ, Reginato MJ. Hypoxia suppression of Bim and Bmf blocks anoikis and luminal clearing during mammary morphogenesis. Mol Biol Cell. 2010 Nov 15;21(22):3829-37. doi: 10.1091/mbc.E10-04-0353. Epub 2010 Sep 22. PMID: 20861305; PMCID: PMC2982135.
Av. molecule expression/activity
0.15
0.10
0.05
0.00
****
BIM
Node output
Experiment
CT ECM_low □Hypoxia_high
EXP Hif1a_basal_KD Hif1a_High_KD ECM_low □Hypoxia_high

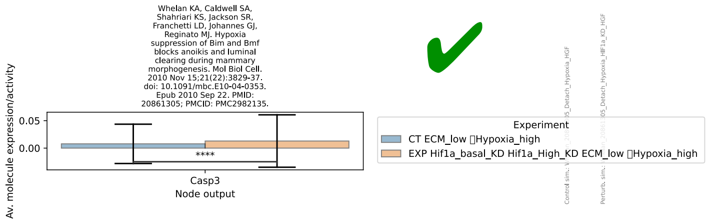
Whelan KA, Caldwell SA, Shahriari KS, Jackson SR, Franchetti LD, Johannes GJ, Reginato MJ. Hypoxia suppression of Bim and Bmf blocks anoikis and luminal clearing during mammary morphogenesis. Mol Biol Cell. 2010 Nov 15;21(22):3829-37. doi: 10.1091/mbc.E10-04-0353. Epub 2010 Sep 22. PMID: 20861305; PMCID: PMC2982135.
Av. molecule expression/activity
0.05
0.00
****
Casp3
Node output
Experiment
CT ECM_low Hypoxia_high
EXP Hif1a_basal_KD Hif1a_High_KD ECM_low Hypoxia_high

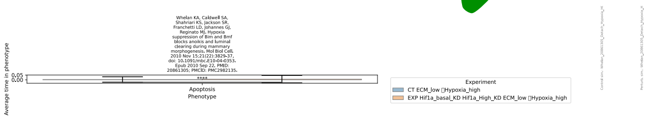
Whelan KA, Caldwell SA, Shahriari KS, Jackson SR, Franchetti LD, Johannes GJ, Reginato MJ. Hypoxia suppression of Bim and Bmf blocks anoikis and luminal clearing during mammary morphogenesis. Mol Biol Cell. 2010 Nov 15;21(22):3829-37. doi: 10.1091/mbc.E10-04-0353. Epub 2010 Sep 22. PMID: 20861305; PMCID: PMC2982135.
Average time in phenotype
0.05
0.00
****
Apoptosis
Phenotype
Experiment
CT ECM_low Hypoxia_high
EXP Hif1a_basal_KD Hif1a_High_KD ECM_low Hypoxia_high

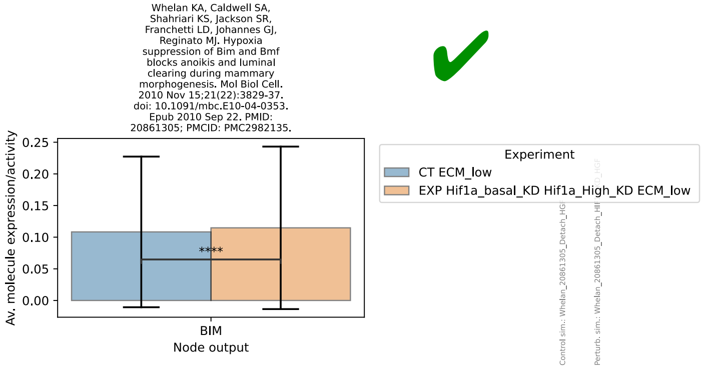
Whelan KA, Caldwell SA, Shahriari KS, Jackson SR, Franchetti LD, Johannes GJ, Reginato MJ. Hypoxia suppression of Bim and Bmf blocks anoikis and luminal clearing during mammary morphogenesis. Mol Biol Cell. 2010 Nov 15;21(22):3829-37. doi: 10.1091/mbc.E10-04-0353. Epub 2010 Sep 22. PMID: 20861305; PMCID: PMC2982135.
Av. molecule expression/activity
0.25
0.20
0.15
0.10
0.05
0.00
****
BIM
Node output
Experiment
CT ECM_low
EXP Hif1a_basal_KD Hif1a_High_KD ECM_low

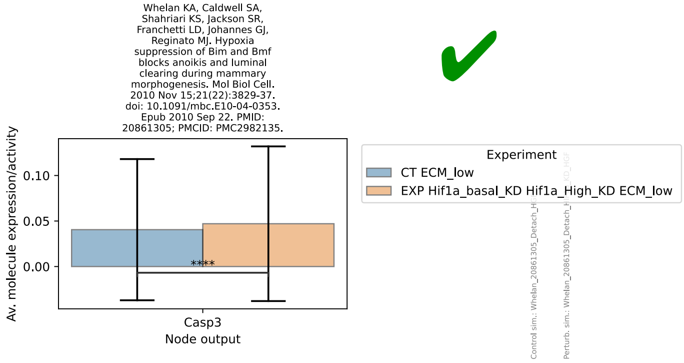
Whelan KA, Caldwell SA, Shahriari KS, Jackson SR, Franchetti LD, Johannes GJ, Reginato MJ. Hypoxia suppression of Bim and Bmf blocks anoikis and luminal clearing during mammary morphogenesis. Mol Biol Cell. 2010 Nov 15;21(22):3829-37. doi: 10.1091/mbc.E10-04-0353. Epub 2010 Sep 22. PMID: 20861305; PMCID: PMC2982135.
Av. molecule expression/activity
0.10
0.05
0.00
****
Casp3
Node output
Experiment
CT ECM_low
EXP Hif1a_basal_KD Hif1a_High_KD ECM_low

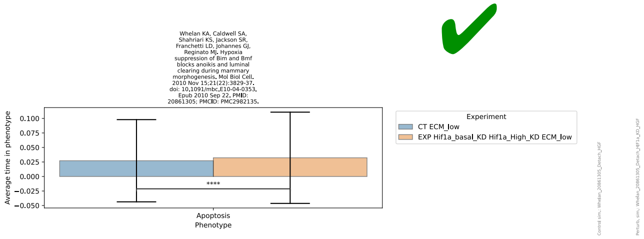
Whelan KA, Caldwell SA, Shahriari KS, Jackson SR, Franchetti LD, Johannes GJ, Reginato MJ. Hypoxia suppression of Bim and Bmf blocks anoikis and luminal clearing during mammary morphogenesis. Mol Biol Cell. 2010 Nov 15;21(22):3829-37. doi: 10.1091/mbc.E10-04-0353. Epub 2010 Sep 22. PMID: 20861305; PMCID: PMC2982135.
Average time in phenotype
0.100
0.075
0.050
0.025
0.000
−0.025
−0.050
****
Apoptosis
Phenotype
Experiment
CT ECM_low
EXP Hif1a_basal_KD Hif1a_High_KD ECM_low
Control sim.: Whelan_20861305_Detach_HGF
Perturb. sim.: Whelan_20861305_Detach_HIF1a_KD_HGF

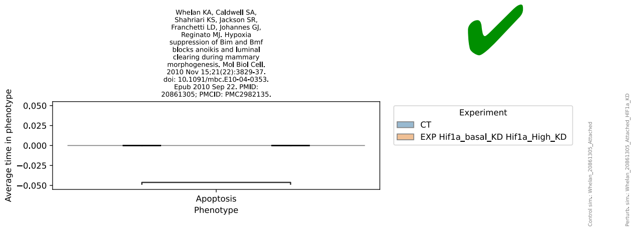
Whelan KA, Caldwell SA, Shahriari KS, Jackson SR, Franchetti LD, Johannes GJ, Reginato MJ. Hypoxia suppression of Bim and Bmf blocks anoikis and luminal clearing during mammary morphogenesis. Mol Biol Cell. 2010 Nov 15;21(22):3829-37. doi: 10.1091/mbc.E10-04-0353. Epub 2010 Sep 22. PMID: 20861305; PMCID: PMC2982135.
Average time in phenotype
0.050
0.025
0.000
−0.025
−0.050
Apoptosis
Phenotype
Experiment
CT
EXP Hif1a_basal_KD Hif1a_High_KD
Control sim.: Whelan_20861305_Attached
Perturb. sim.: Whelan_20861305_Attached_HIF1a_KD

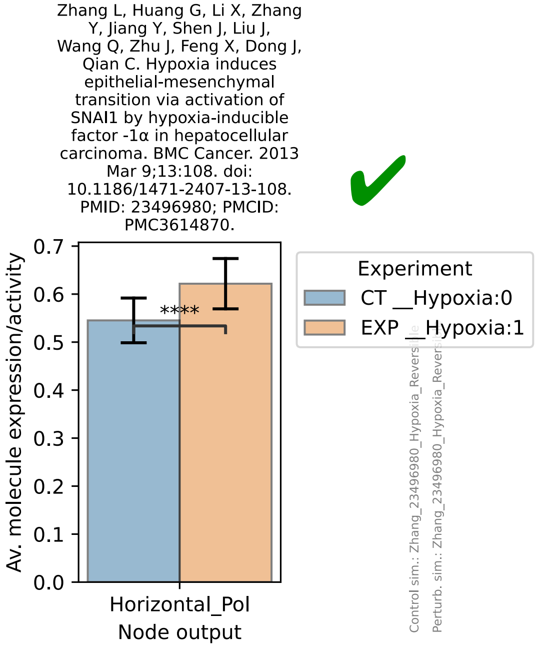
Zhang L, Huang G, Li X, Zhang Y, Jiang Y, Shen J, Liu J, Wang Q, Zhu J, Feng X, Dong J, Qian C. Hypoxia induces epithelial-mesenchymal transition via activation of SNAI1 by hypoxia-inducible factor -1α in hepatocellular carcinoma. BMC Cancer. 2013 Mar 9;13:108. doi: 10.1186/1471-2407-13-108. PMID: 23496980; PMCID: PMC3614870.
Av. molecule expression/activity
0.7
0.6
0.5
0.4
0.3
0.2
0.1
0.0
****
Horizontal_Pol
Node output
Experiment
CT __Hypoxia:0
EXP __Hypoxia:1
Control sim.: Zhang_23496980_Hypoxia_Revers
Perturb. sim.: Zhang_23496980_Hypoxia_Rever

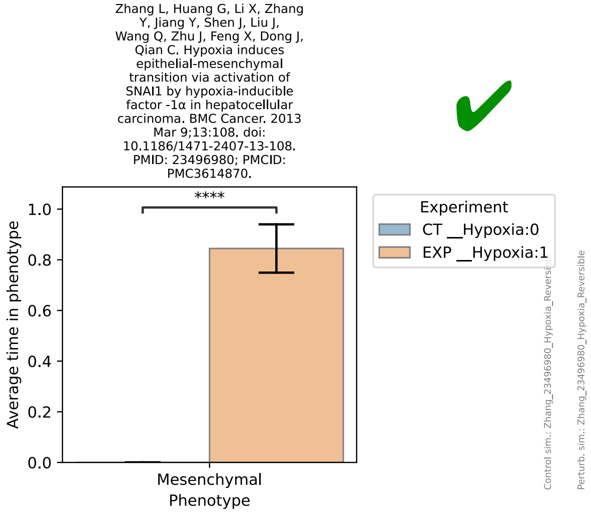
Zhang L, Huang G, Li X, Zhang Y, Jiang Y, Shen J, Liu J, Wang Q, Zhu J, Feng X, Dong J, Qian C. Hypoxia induces epithelial-mesenchymal transition via activation of SNAI1 by hypoxia-inducible factor -1α in hepatocellular carcinoma. BMC Cancer. 2013 Mar 9;13:108. doi: 10.1186/1471-2407-13-108. PMID: 23496980; PMCID: PMC3614870.
Average time in phenotype
1.0
0.8
0.6
0.4
0.2
0.0
****
Mesenchymal
Phenotype
Experiment
CT __Hypoxia:0
EXP __Hypoxia:1
Control sim.: Zhang_23496980_Hypoxia_Reversi
Perturb. sim.: Zhang_23496980_Hypoxia_Reversible

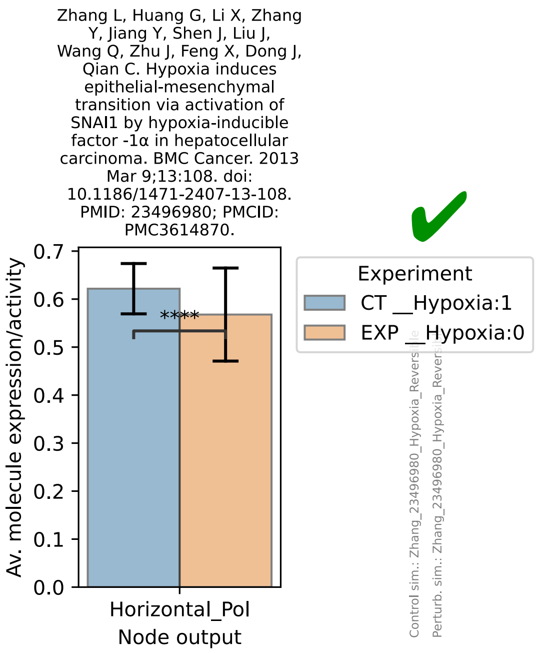
Zhang L, Huang G, Li X, Zhang Y, Jiang Y, Shen J, Liu J, Wang Q, Zhu J, Feng X, Dong J, Qian C. Hypoxia induces epithelial-mesenchymal transition via activation of SNAI1 by hypoxia-inducible factor -1α in hepatocellular carcinoma. BMC Cancer. 2013 Mar 9;13:108. doi: 10.1186/1471-2407-13-108. PMID: 23496980; PMCID: PMC3614870.
Av. molecule expression/activity
0.7
0.6
0.5
0.4
0.3
0.2
0.1
0.0
****
Horizontal_Pol
Node output
Experiment
CT __Hypoxia:1
EXP __Hypoxia:0
Control sim.: Zhang_23496980_Hypoxia_Revers
Perturb. sim.: Zhang_23496980_Hypoxia_Rever

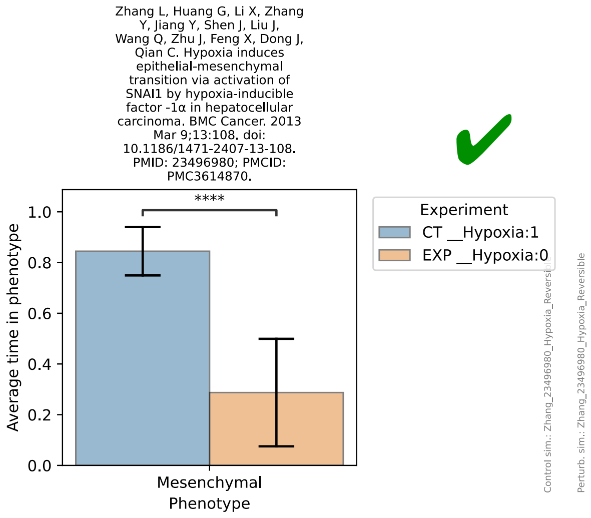
Zhang L, Huang G, Li X, Zhang Y, Jiang Y, Shen J, Liu J, Wang Q, Zhu J, Feng X, Dong J, Qian C. Hypoxia induces epithelial-mesenchymal transition via activation of SNAI1 by hypoxia-inducible factor -1α in hepatocellular carcinoma. BMC Cancer. 2013 Mar 9;13:108. doi: 10.1186/1471-2407-13-108. PMID: 23496980; PMCID: PMC3614870.
Average time in phenotype
1.0
0.8
0.6
0.4
0.2
0.0
****
Mesenchymal
Phenotype
Experiment
CT __Hypoxia:1
EXP __Hypoxia:0
Control sim.: Zhang_23496980_Hypoxia_Reversi
Perturb. sim.: Zhang_23496980_Hypoxia_Reversible

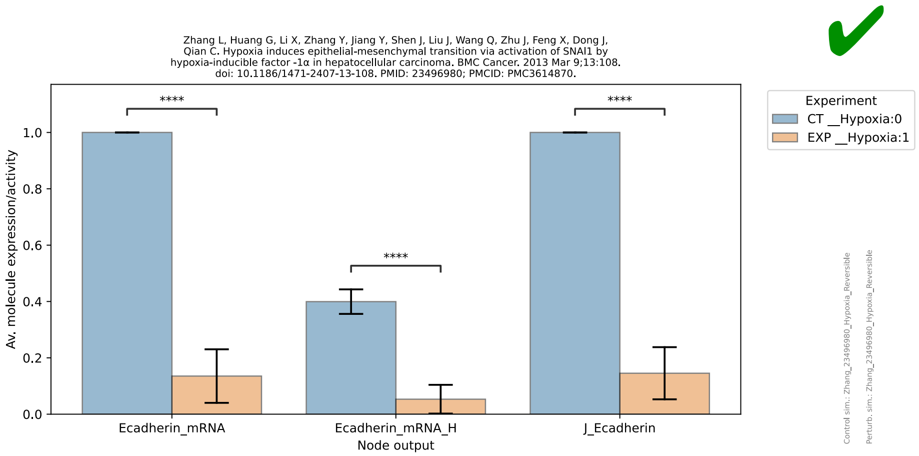
Zhang L, Huang G, Li X, Zhang Y, Jiang Y, Shen J, Liu J, Wang Q, Zhu J, Feng X, Dong J, Qian C. Hypoxia induces epithelial-mesenchymal transition via activation of SNAI1 by hypoxia-inducible factor -1α in hepatocellular carcinoma. BMC Cancer. 2013 Mar 9;13:108. doi: 10.1186/1471-2407-13-108. PMID: 23496980; PMCID: PMC3614870.
Av. molecule expression/activity
1.0
0.8
0.6
0.4
0.2
0.0
****
****
****
Ecadherin_mRNA
Ecadherin_mRNA_H
J_Ecadherin
Node output
Experiment
CT __Hypoxia:0
EXP __Hypoxia:1

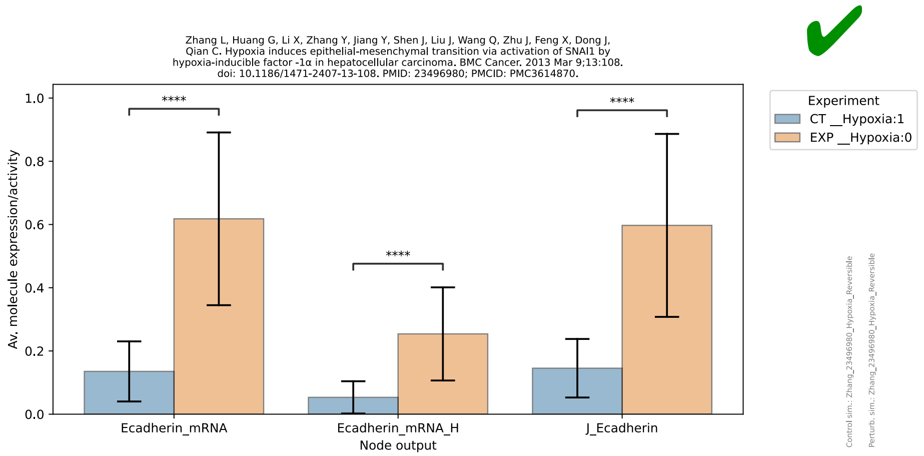
Zhang L, Huang G, Li X, Zhang Y, Jiang Y, Shen J, Liu J, Wang Q, Zhu J, Feng X, Dong J, Qian C. Hypoxia induces epithelial-mesenchymal transition via activation of SNAI1 by hypoxia-inducible factor -1α in hepatocellular carcinoma. BMC Cancer. 2013 Mar 9;13:108. doi: 10.1186/1471-2407-13-108. PMID: 23496980; PMCID: PMC3614870.
Av. molecule expression/activity
1.0
0.8
0.6
0.4
0.2
0.0
****
****
****
Ecadherin_mRNA
Ecadherin_mRNA_H
J_Ecadherin
Node output
Experiment
CT __Hypoxia:1
EXP __Hypoxia:0

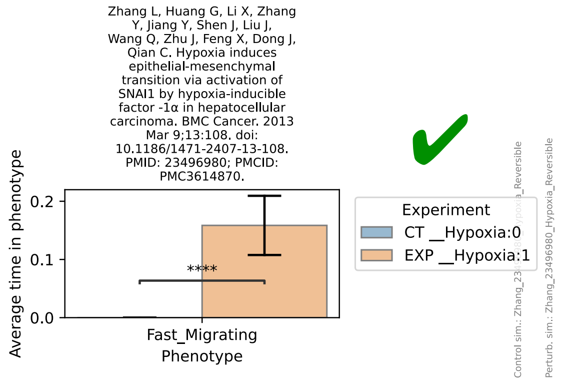
Zhang L, Huang G, Li X, Zhang Y, Jiang Y, Shen J, Liu J, Wang Q, Zhu J, Feng X, Dong J, Qian C. Hypoxia induces epithelial-mesenchymal transition via activation of SNAI1 by hypoxia-inducible factor -1α in hepatocellular carcinoma. BMC Cancer. 2013 Mar 9;13:108. doi: 10.1186/1471-2407-13-108. PMID: 23496980; PMCID: PMC3614870.
Average time in phenotype
0.2
0.1
0.0
****
Fast_Migrating
Phenotype
Experiment
CT __Hypoxia:0
EXP __Hypoxia:1
Perturb. sim.: Zhang_23496980_Hypoxia_Reversible

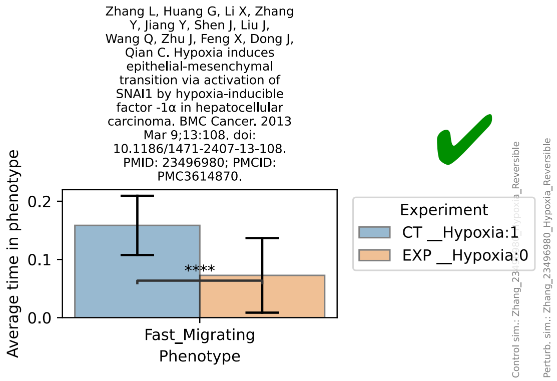
Zhang L, Huang G, Li X, Zhang Y, Jiang Y, Shen J, Liu J, Wang Q, Zhu J, Feng X, Dong J, Qian C. Hypoxia induces epithelial-mesenchymal transition via activation of SNAI1 by hypoxia-inducible factor -1α in hepatocellular carcinoma. BMC Cancer. 2013 Mar 9;13:108. doi: 10.1186/1471-2407-13-108. PMID: 23496980; PMCID: PMC3614870.
Average time in phenotype
0.2
0.1
0.0
****
Fast_Migrating
Phenotype
Experiment
CT __Hypoxia:1
EXP __Hypoxia:0
Perturb. sim.: Zhang_23496980_Hypoxia_Reversible

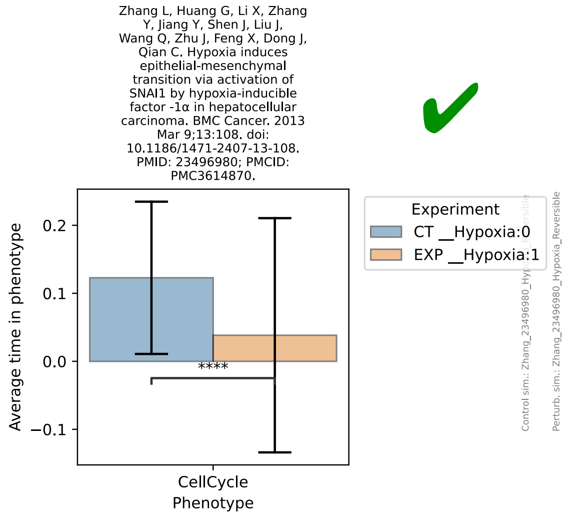
Zhang L, Huang G, Li X, Zhang Y, Jiang Y, Shen J, Liu J, Wang Q, Zhu J, Feng X, Dong J, Qian C. Hypoxia induces epithelial-mesenchymal transition via activation of SNAI1 by hypoxia-inducible factor -1α in hepatocellular carcinoma. BMC Cancer. 2013 Mar 9;13:108. doi: 10.1186/1471-2407-13-108. PMID: 23496980; PMCID: PMC3614870.
Average time in phenotype
0.2
0.1
0.0
−0.1
****
CellCycle
Phenotype
Experiment
CT __Hypoxia:0
EXP __Hypoxia:1
Perturb. sim.: Zhang_23496980_Hypoxia_Reversible

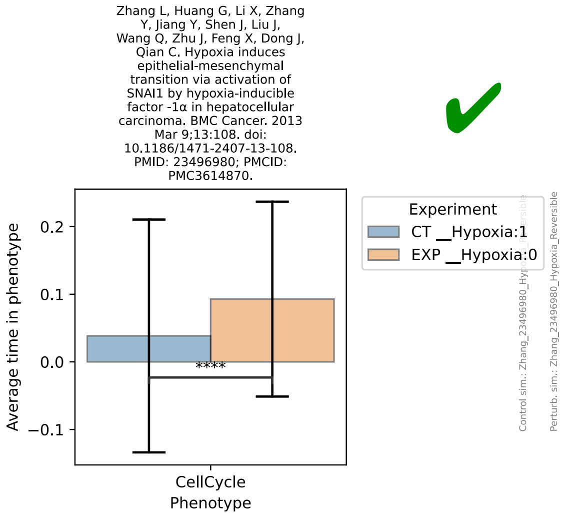
Zhang L, Huang G, Li X, Zhang Y, Jiang Y, Shen J, Liu J, Wang Q, Zhu J, Feng X, Dong J, Qian C. Hypoxia induces epithelial-mesenchymal transition via activation of SNAI1 by hypoxia-inducible factor -1α in hepatocellular carcinoma. BMC Cancer. 2013 Mar 9;13:108. doi: 10.1186/1471-2407-13-108. PMID: 23496980; PMCID: PMC3614870.
Average time in phenotype
0.2
0.1
0.0
−0.1
****
CellCycle
Phenotype
Experiment
CT __Hypoxia:1
EXP __Hypoxia:0
Perturb. sim.: Zhang_23496980_Hypoxia_Reversible

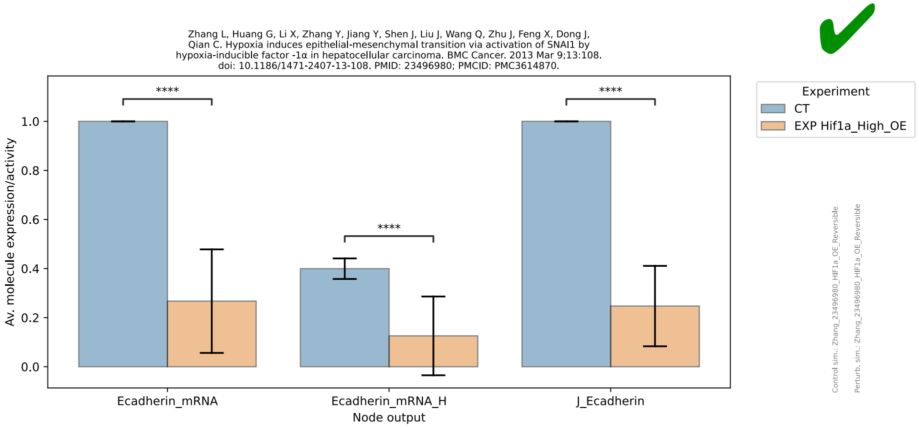
Zhang L, Huang G, Li X, Zhang Y, Jiang Y, Shen J, Liu J, Wang Q, Zhu J, Feng X, Dong J, Qian C. Hypoxia induces epithelial-mesenchymal transition via activation of SNAI1 by hypoxia-inducible factor -1α in hepatocellular carcinoma. BMC Cancer. 2013 Mar 9;13:108. doi: 10.1186/1471-2407-13-108. PMID: 23496980; PMCID: PMC3614870.
Av. molecule expression/activity
Ecadherin_mRNA
Ecadherin_mRNA_H
J_Ecadherin
Node output
Experiment
CT
EXP Hif1a_High_OE
Control sim.: Zhang_23496980_HIF1a_OE_Reversible
Perturb. sim.: Zhang_23496980_HIF1a_OE_Reversible

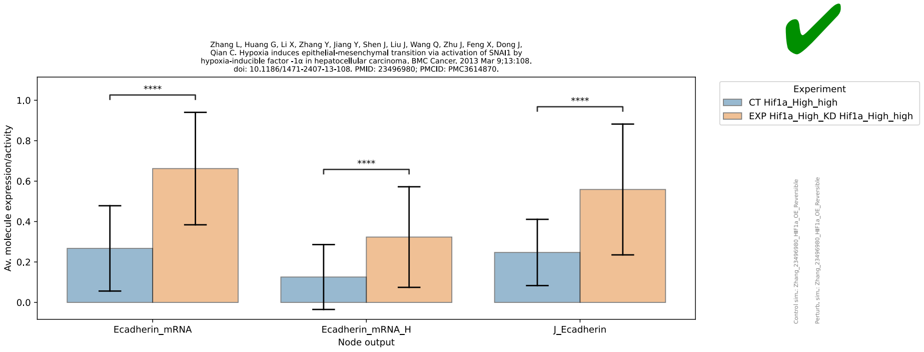
Zhang L, Huang G, Li X, Zhang Y, Jiang Y, Shen J, Liu J, Wang Q, Zhu J, Feng X, Dong J, Qian C. Hypoxia induces epithelial-mesenchymal transition via activation of SNAI1 by hypoxia-inducible factor -1α in hepatocellular carcinoma. BMC Cancer. 2013 Mar 9;13:108. doi: 10.1186/1471-2407-13-108. PMID: 23496980; PMCID: PMC3614870.
Av. molecule expression/activity
Ecadherin_mRNA
Ecadherin_mRNA_H
J_Ecadherin
Node output
Experiment
CT Hif1a_High_high
EXP Hif1a_High_KD Hif1a_High_high

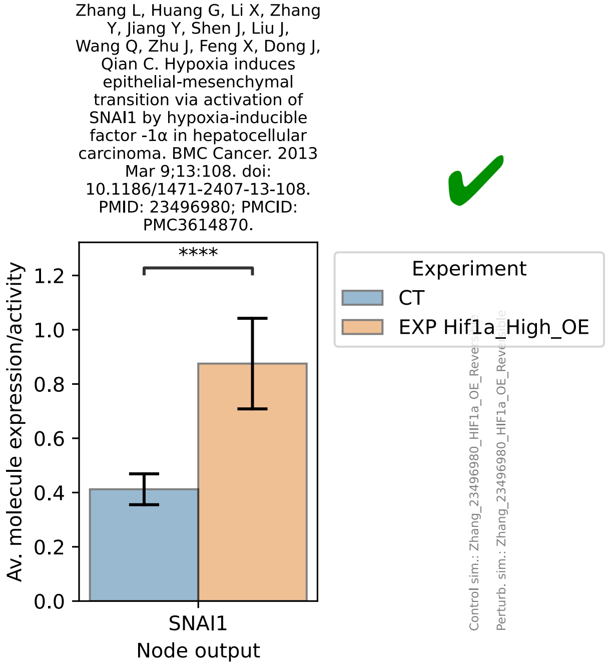
Zhang L, Huang G, Li X, Zhang Y, Jiang Y, Shen J, Liu J, Wang Q, Zhu J, Feng X, Dong J, Qian C. Hypoxia induces epithelial-mesenchymal transition via activation of SNAI1 by hypoxia-inducible factor -1α in hepatocellular carcinoma. BMC Cancer. 2013 Mar 9;13:108. doi: 10.1186/1471-2407-13-108. PMID: 23496980; PMCID: PMC3614870.
Av. molecule expression/activity
SNAI1
Node output
Experiment
CT
EXP Hif1a_High_OE

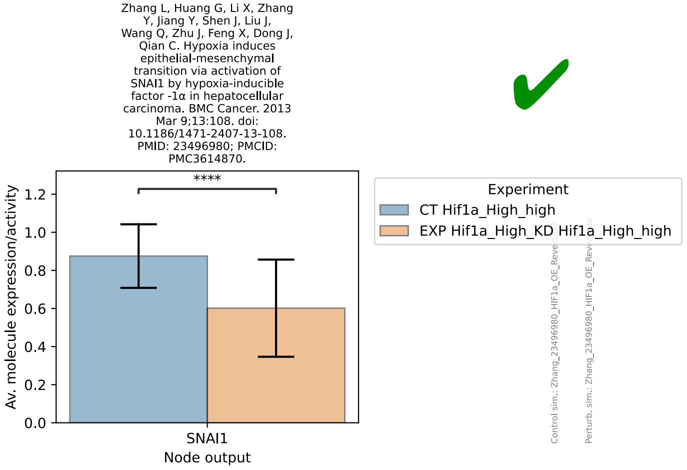
Zhang L, Huang G, Li X, Zhang Y, Jiang Y, Shen J, Liu J, Wang Q, Zhu J, Feng X, Dong J, Qian C. Hypoxia induces epithelial-mesenchymal transition via activation of SNAI1 by hypoxia-inducible factor -1α in hepatocellular carcinoma. BMC Cancer. 2013 Mar 9;13:108. doi: 10.1186/1471-2407-13-108. PMID: 23496980; PMCID: PMC3614870.
Av. molecule expression/activity
SNAI1
Node output
Experiment
CT Hif1a_High_high
EXP Hif1a_High_KD Hif1a_High_high

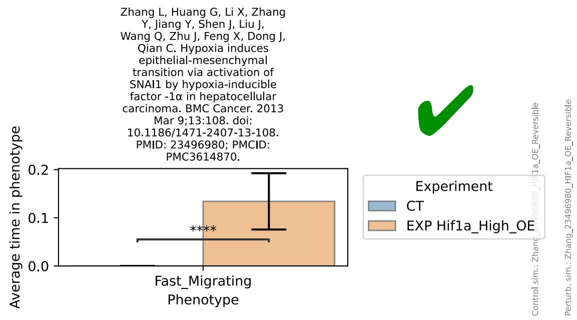
Zhang L, Huang G, Li X, Zhang Y, Jiang Y, Shen J, Liu J, Wang Q, Zhu J, Feng X, Dong J, Qian C. Hypoxia induces epithelial-mesenchymal transition via activation of SNAI1 by hypoxia-inducible factor -1α in hepatocellular carcinoma. BMC Cancer. 2013 Mar 9;13:108. doi: 10.1186/1471-2407-13-108. PMID: 23496980; PMCID: PMC3614870.
Average time in phenotype
0.2
0.1
0.0
****
Fast_Migrating
Phenotype
Experiment
CT
EXP Hif1a_High_OE
Control sim.: Zhang_23496980_HIF1a_OE_Reversible
Perturb. sim.: Zhang_23496980_HIF1a_OE_Reversible

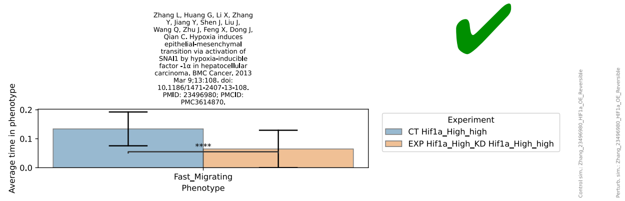
Zhang L, Huang G, Li X, Zhang Y, Jiang Y, Shen J, Liu J, Wang Q, Zhu J, Feng X, Dong J, Qian C. Hypoxia induces epithelial-mesenchymal transition via activation of SNAI1 by hypoxia-inducible factor -1α in hepatocellular carcinoma. BMC Cancer. 2013 Mar 9;13:108. doi: 10.1186/1471-2407-13-108. PMID: 23496980; PMCID: PMC3614870.
Average time in phenotype
0.2
0.1
0.0
****
Fast_Migrating
Phenotype
Experiment
CT Hif1a_High_high
EXP Hif1a_High_KD Hif1a_High_high
Control sim.: Zhang_23496980_HIF1a_OE_Reversible
Perturb. sim.: Zhang_23496980_HIF1a_OE_Reversible
